# The blueprint of human functional architecture shifts from cognition to anatomy during perturbations of consciousness

**DOI:** 10.64898/2026.06.07.730661

**Authors:** Andrea I. Luppi, Dragana Manasova, Justine Y. Hansen, Zhen-Qi Liu, Asa Farahani, Yonatan Sanz Perl, Jakub Vohryzek, Daniel Golkowski, Andreas Ranft, Rüdiger Ilg, Denis Jordan, Vincent Bonhomme, Audrey Vanhaudenhuyse, Athena Demertzi, Oceane Jaquet, Mohamed Ali Bahri, Naji L. N. Alnagger, Paolo Cardone, Lorina Naci, Adrian M. Owen, John D. Pickard, Guy B. Williams, Judith Allanson, Enrico Amico, Danilo Bzdok, Jacobo D. Sitt, David K. Menon, Emmanuel A. Stamatakis, Bratislav Misic

**Affiliations:** Department of Psychiatry, University of Oxford, Oxford, UK; Division of Information Engineering and St John’s College, University of Cambridge, Cambridge, UK; Montréal Neurological Institute, McGill University, Montréal, QC, Canada; Institute for Medical Engineering and Science, Massachusetts Institute of Technology, Cambridge, Massachusetts, USA; Children’s Hospital of Philadelphia, University of Pennsylvania, Philadelphia, PA, USA; Pompeu Fabra University, Barcelona, Spain; Technical University of Munich, Munich, Germany; Department of Neurology, University Hospital Heidelberg, Heidelberg, Germany; Department of Anesthesia and Intensive Care Medicine, Liège University Hospital, Liège, Belgium; Anesthesia and Perioperative Neuroscience Laboratory, GIGA-Consciousness, GIGA Institute, Liège University, Liège, Belgium; Interdisciplinary Algology Center, Liège University Hospital, Liège, Belgium; Conscious Care Lab, GIGA Consciousness, GIGA Institute, University of Liège, Belgium; Physiology of Cognition Lab, GIGA-Cyclotron Research Center In Vivo Imaging, University of Liège, Liège, Belgium; GIGA-Cyclotron Research Center Human Imaging, University of Liège, Liège, Belgium; Coma Science Group, GIGA-Consciousness, GIGA Institute, University of Liège, Liège, Belgium; School of Psychology, Trinity College Dublin, Dublin, Ireland; Department of Physiology and Pharmacology, and Department of Psychology, Western University, London, ON, Canada; Department of Clinical Neurosciences, University of Cambridge, Cambridge, UK; University of Birmingham, Birmingham, UK; MILA-Quebec Artificial Intelligence Institute, Montréal, QC, Canada; Paris Brain Institute, Paris, France; Division of Anaesthesia, University of Cambridge, Cambridge, UK

## Abstract

Consciousness and cognition arise from the ongoing interactions between brain regions. Synchronous fluctuations of fMRI signals may indicate that two brain regions perform similar cognitive functions, but neural interactions are also constrained by anatomical connectivity and regions’ molecular, cytoarchitectonic, and metabolic profiles. Here we disentangle the respective contributions of ongoing cognition and multimodal neurobiological constraints in shaping functional connectivity. We jointly contextualise haemodynamic FC against eight distinct multimodal representations of the human connectome: (i) structural connectivity from diffusion tractography; (ii) spatial embedding; (iii) similarity of transcriptional profiles from gene expression; (iv) similarity of receptor profiles from Positron Emission Tomography; (v) laminar profile similarity from histology; (vi) correlated electrophysiological activity from magnetoencephalography; (vii) correlated metabolic activity from PET glucose uptake; (viii) coordinated activation across 123 cognitive operations from the NeuroSynth meta-analytic engine. We demonstrate that cognitive co-activation is the dominant predictor of inter-regional fMRI synchrony in the awake human brain, even when quantified using intracranial electrical stimulation. Crucially, this predominance of cognitive co-activation for shaping functional connectivity is systematically obliterated across five datasets of pharmacological and pathological perturbations of consciousness (chronic disorders of consciousness; anaesthesia with sevoflurane, propofol, or ketamine) when cognition is disconnected from the environment or altogether abolished. Altogether, we show that multimodal predictors of functional architecture shift away from cognitive co-activation and toward anatomical-molecular constraints during pharmacological and pathological perturbations of consciousness.

## INTRODUCTION

How brain organisation enables conscious experience is a question of enduring neuroscientific interest. Noninvasive neuroimaging with functional MRI has revealed a complex functional architecture, with brain regions continuously forming and dissolving coalitions of synchronous haemodynamic activity [1–3]. Synchronous co-fluctuations are taken to indicate that two regions are involved in similar cognitive functions: hence the name of ‘functional’ connectivity (FC) [1–3]. Indeed, patterns of haemodynamic FC bear predictive value for cognition, behaviour, pathology, and individual identity [4–10]. Even in the absence of explicit tasks, FC is far from random, instead recapitulating task-induced coactivations [11–16]. Coordinated activity across tasks reflects which regions are consistently recruited by the same cognitive operations—such as attention, memory, emotion or language—thereby indicating similarity of cognitive function. Further supporting the cognitive relevance of haemodynamic FC, when consciousness is lost due to general anaesthesia or disorders of consciousness, FC reverts to the underlying scaffold of anatomical connections between regions [17–22], and information about individual identity is lost [23].

At the same time, a rich literature has identified overlapping macro- and microarchitectural constraints on functional connectivity [24, 25]. In addition to the role of direct and indirect structural connectivity [26–36], functional interactions between regions also depend on whether they exhibit similar profiles of neurotransmitter receptors [37–40]; similar metabolic demand [41– 43] and preferred neurophysiological frequencies [44– 49]; spatial embedding [50]; and on the similarity of their broader molecular [51–55] and microarchitectural fingerprints [24, 56–62]. Each of these features of brain organisation provides a distinct, complementary way to understand how brain structure informs and constrains function [24–26], and how the brain is reorganised following injury [63]. Yet previous studies have not directly compared these contraposed accounts of FC.

The question arises: What are the respective contributions of brain architecture and ongoing cognition for shaping functional connectivity? Here we pursue this question with a two-pronged strategy. To go beyond conventional univariate comparisons, we adopt a multivariate approach to jointly contextualise haemodynamic FC against eight distinct ways of characterising structural and functional relationships between cortical regions (Fig. 1). Specifically, we consider: (i) spatial distance between regions; (ii) the structural white matter connectome reconstructed from diffusion MRI tractography; (iii) covariance of regional gene expression profiles from spatial transcriptomics [52]; (iv) similarity of regional receptor profiles from *in vivo* Positron Emission Tomography (PET) in *>* 1 200 participants [38]; (v) laminar profile similarity from *ex vivo* histological cell staining [62, 64]; (vi) correlated electrophysiological activity from source-resolved magnetoencephalography (MEG) [45]; (vii) correlated metabolic activity measured with dynamic fluorodeoxyglucose uptake (FDG-PET) [41]; and finally (viii) co-activation across 123 cognitive operations (Table 1) from NeuroSynth metaanalysis, aggregated over 14 000 neuroimaging studies (cognitive co-activations, for short) [12, 14, 15, 65]. These multimodal anatomical and functional networks [24] provide a comprehensive characterisation of the relationships between human cortical regions (Fig. 1).

**Figure 1.**
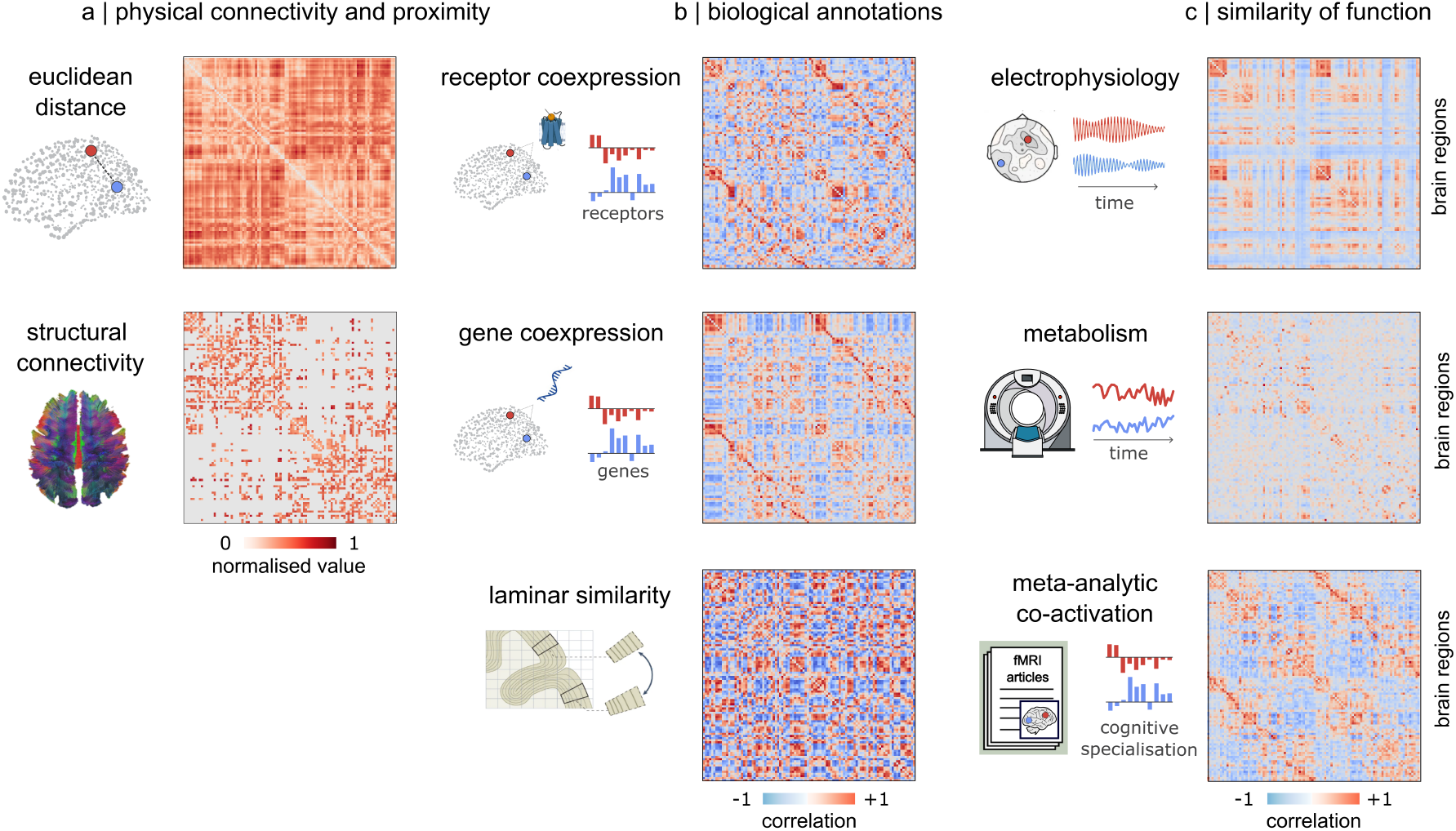
Structural and functional networks of the human brain. (**a**) Physical relationships between brain regions: Euclidean distance between regions (mm), and structural connectivity (number of white matter fibers reconstructed from *in vivo* diffusion tractography). For visualisation purposes, values are normalised to the maximum. (**b**) Local biological similarity: correlation of regions’ receptor expression from Positron Emission Tomography; correlation of regions’ gene expression from spatial transcriptomics; and correlation of laminar profiles from histology. (**c**) Similarity of function: correlated metabolic activity from FDG-PET; correlated electrodynamic activity from MEG; and correlation of involvement across 123 cognitive operations from NeuroSynth meta-analysis. The networks are available at https://github.com/netneurolab/hansen_many_networks/tree/v1.0.0.

**TABLE 1.** NeuroSynth terms. Terms that overlapped between the NeuroSynth database [65] and the Cognitive Atlas [177] were included in the analysis.

|  |  |  |  |  |
| --- | --- | --- | --- | --- |
| action | adaptation | addiction | anticipation | anxiety |
| arousal | association | attention | autobiographical memory | balance |
| belief | categorization | cognitive control | communication | competition |
| concept | consciousness | consolidation | context | coordination |
| decision | decision making | detection | discrimination | distraction |
| eating | efficiency | effort | emotion | emotion regulation |
| empathy | encoding | episodic memory | expectancy | expertise |
| extinction | face recognition | facial expression | familiarity | fear |
| fixation | focus | gaze | goal | hyperactivity |
| imagery | impulsivity | induction | inference | inhibition |
| insight | integration | intelligence | intention | interference |
| judgment | knowledge | language | language comprehension | learning |
| listening | localization | loss | maintenance | manipulation |
| meaning | memory | memory retrieval | mental imagery | monitoring |
| mood | morphology | motor control | movement | multisensory |
| naming | navigation | object recognition | pain | perception |
| planning | priming | psychosis | reading | reasoning |
| recall | recognition | rehearsal | reinforcement learning | response inhibition |
| response selection | retention | retrieval | reward anticipation | rhythm |
| risk | rule | salience | search | selective attention |
| semantic memory | sentence comprehension | skill | sleep | social cognition |
| spatial attention | speech perception | speech production | strategy | strength |
| stress | sustained attention | task difficulty | thought | uncertainty |
| updating | utility | valence | verbal fluency | visual attention |
| visual perception | word recognition | working memory |  |  |

Next, we employ general anaesthesia as an experimental paradigm to disentangle the respective contributions of anatomy and cognition to haemodynamic FC. General anaesthesia is an acute pharmacological manipulation that profoundly alters or altogether abolishes cognition, as indicated by complete cessation of the organism’s ability to process and respond to stimuli in the environment. Crucially, anaesthesia leaves anatomy intact (unlike e.g. a brain lesion), providing an ideal means to disentangle whether anatomical contributions to haemodynamic connectivity change in the presence versus absence of ongoing cognition. Specifically, we consider datasets of general anaesthesia with propofol [66], sevoflurane [67], and a high dose of ketamine [68], all inducing disconnection from the environment with complete loss of behavioural responsiveness. We then generalise results beyond pharmacology, in two datasets of pathological perturbation of consciousness due to brain injury (chronic disorders of consciousness) [63, 66]. To foreshadow our main results, our multimodal approach reveals that pharmacological and pathological perturbations of consciousness do not only increase the contribution of anatomical-molecular constraints for predicting fMRI functional connectivity, but also induce a shift away from cognitive co-activation as predictor of functional architecture.

## RESULTS

### Cognitive co-activation is the best predictor of haemodynamic functional connectivity

Here we ask how perturbation of consciousness and disconnection from the environment induced by injury or pharmacology - whereby cognition is profoundly perturbed or altogether abolished - reshape the relationship between different types of functional and structural networks in the human brain. To disambiguate the contributions of each structural and functional network to FC, we use dominance analysis, a multivariate approach that optimally distributes the fit (variance explained) of a multiple regression model across individual predictors. This procedure is well suited to the present analysis because unlike methods based on regression coefficients or univariate correlations, it estimates predictor importance while accounting for shared variance among correlated predictors [69, 70].

The first discovery is that across all datasets, by far the best predictor of haemodynamic functional connectivity is the network of regions’ co-activation across cognitive operations, derived from NeuroSynth meta-analysis. Similarity across NeuroSynth maps consistently accounts for *>* 50% of the explained variance in the awake human brain’s BOLD FC (Fig. 2). It is remarkable that restingstate coordinated haemodynamics are better recapitulated by task-related co-activations aggregated from the literature, than by coordinated metabolic activity from FDG-PET, or coordinated electrodynamics from MEG. This is surprising because one might have expected that these alternative functional networks would be the most relevant predictors of BOLD FC. First, because both FDG-PET FC and MEG-FC are obtained directly from time-series of regional activity, like BOLD FC but *unlike* metaanalytic co-activation. Second, FDG-PET FC and MEG-FC both reflect the same kind of process as BOLD FC: synchronous activity over time, in the absence of any task. Instead, we find that regions’ propensity for exhibiting spontaneous fMRI co-fluctuations is best predicted by their propensity for activating together in response to a variety of tasks. This observation is consistent with observations that the spontaneous fluctuations of the BOLD signal are not random, instead recapitulating cognitively-relevant co-activations [11–16].

**Figure 2.**
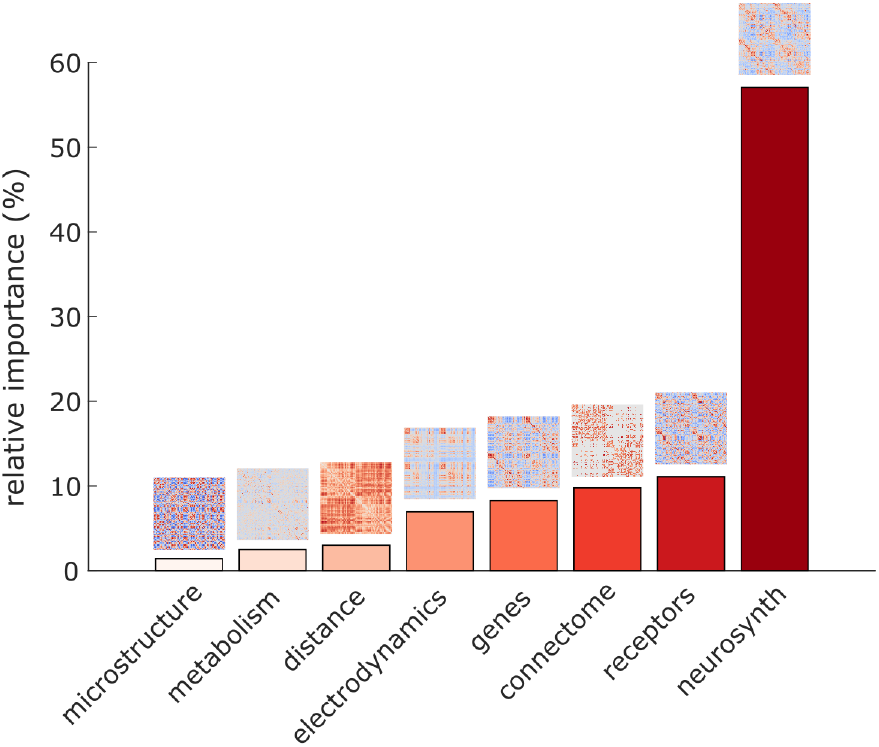
Meta-analytic co-activation is the dominant predictor of fMRI functional connectivity. Bars indicate the mean percentage of relative importance of each network for predicting fMRI functional connectivity, averaged across the baseline conditions of all datasets. See Fig. 3 for dataset-by-dataset breakdown.

### Pharmacological and pathological perturbations of consciousness induce a shift away from cognitive co-activation

This picture is drastically altered upon pharmacological or pathological perturbations of consciousness. Across anaesthesia with propofol, sevoflurane, and a high dose of ketamine (all inducing complete loss of behavioural responsiveness) and also in DOC patients, we find that cognitive co-activations from NeuroSynth significantly and consistently reduce their relative importance as predictors of FC, compared with baseline (Fig. 3a-d). This effect is then reversed upon recovery of responsiveness from both sevoflurane and propofol anaesthesia (Fig. 3b-d and Fig. S1). In other words, when behavioural responsiveness is lost, and cognition is disconnected from the environment or altogether abolished, we see that the spontaneous co-fluctuations of human haemodynamics cease to mirror regions’ co-involvement across cognitive operations (Fig. 3b-d and Fig. S1). We confirm this by showing that the network of meta-analytic co-activations is the most impacted by anaesthesia and DOC, in terms of exhibiting the greatest magnitude of change in relative importance (Fig. 3e).

**Figure 3.**
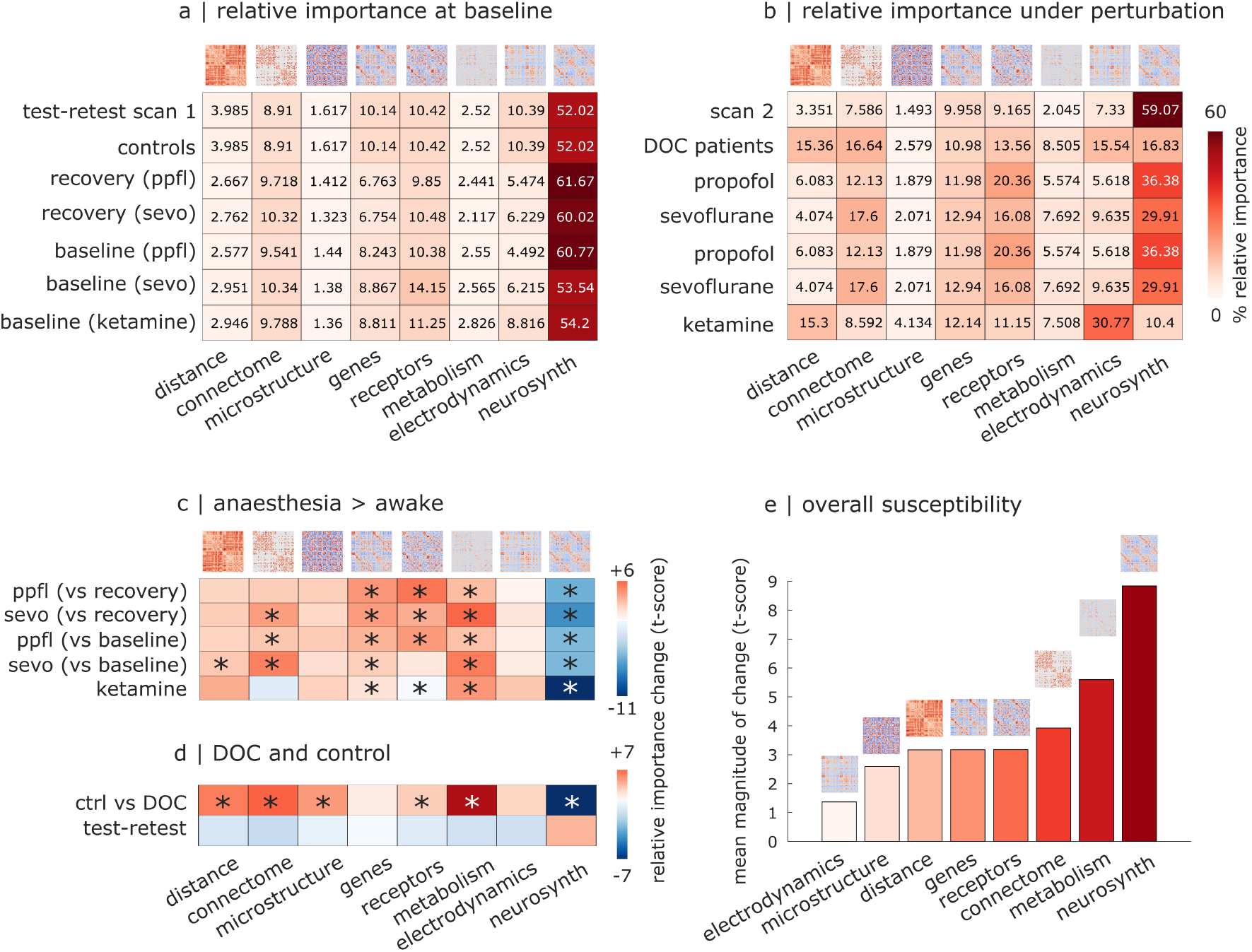
Contribution of cognitively-relevant co-activations to functional connectivity is consistent across datasets but reduced under perturbed consciousness. (**a**) Mean percent relative importance from dominance analysis for predicting FC from structural and functional brain networks, for the baseline scans (pre-anaesthetic wakefulness, post-anaesthetic recovery of responsiveness, and healthy controls). (**b**) Mean percent relative importance from dominance analysis for predicting FC from structural and functional brain networks, for the perturbed consciousness datasets (anaesthesia and DOC patients). For comparison, the second scan of the test-retest dataset is also included. (**c**) Change in percentage of relative importance under anaesthesia. (**d**) Change in percentage of relative importance in DOC patients or in a second scan of the same healthy controls. *, *p* < 0.05 (head motion is included as covariate of no interest). For (c) and (d), red color-scale indicates perturbed > baseline; blue color-scale indicates baseline > perturbed. See Tables S1-S2 for statistical reporting. Note that the first scan of the test-retest dataset is used as the control group for the DOC patients. For the sevoflurane and propofol datasets, the same anaesthetised scan is compared against both baseline wakefulness and post-anaesthetic recovery of responsiveness. (**e**) Mean magnitude of change (t-score) across contrasts from the anaesthesia and DOC datasets.

Based on the literature, we might expect to observe an increased contribution from the structural connectome in DOC patients as well as sevoflurane and propofol anaesthesia [17, 18, 21]—which is indeed what we find. However, we do not observe this for ketamine anaesthesia. The ketamine dose induced complete loss of behavioural responsiveness, with Ramsay score of 5-6 [68] comparable to the propofol and sevoflurane datasets [66, 67]. However, participants in the ketamine dataset were not unconscious [68, 71]. Indeed, the present results are consistent with a recent report in the same ketamine dataset, which also found that unlike propofol and DOC, ketamine does not increase structure-function coupling in the human brain [71]. Notably, we also find that under perturbations of consciousness, haemodynamic FC becomes better predicted by regional coexpression of genes and receptors, and especially by the slow metabolic co-fluctuations measured by FDG-PET (Fig. 3b,c). These observations are consistent across pharmacological and pathological perturbations, including ketamine, and are not driven by differences in head motion, which we include as covariate of no interest. No changes in predictive performance are observed for either microstructural profile similarity (except in DOC) or MEG electrodynamic similarity. Spatial distance also does not exhibit consistent changes. Thus, under perturbations of consciousness we see consistent increases in the coupling between FC and different aspects of brain anatomy. In contrast, no significant changes are observed in a test-retest dataset (Fig. 3d), further confirming that the relative contributions of different brain networks to FC are not only highly consistent across datasets, but also stable over time within the same dataset.

We also ask whether the dominance of meta-analytic co-activation for predicting fMRI FC, and the impact of anaesthesia and DOC, are primarily driven by connections involving specific cortical territories. We divide the cortex in unimodal and transmodal regions, according to the canonical map of [72]. We then repeat the same analysis using only connections between unimodal regions; or only connections between transmodal regions; or only connections involving one unimodal and one transmodal region (Fig. S2). In all three cases, we find that meta-analytic co-activation is the dominant predictor of haemodynamic FC at baseline (Fig. S2) and also the predictor most affected by anaesthesia and DOC (Fig. S3 and Fig. S4). However, both the relative dominance of meta-analytic co-activation and the extent to which it is affected by anaesthesia and DOC are most pronounced for connections involving transmodal cortex (especially transmodal-transmodal, where results mostly resemble the whole-brain analysis), and least pronounced for unimodal-unimodal connections (Fig. S2 and Fig. S4). These results indicate that the relevance of cognitive co-activations for tracking state of consciousness is especially prominent for the cortical types most involved in higher-order cognition.

### Multivariate classification of consciousness and prognosis

To complement the main group-level analysis we show that the relative contributions of different structural and functional networks in predicting haemodynamic FC can be used to classify perturbed versus unperturbed consciousness at the single-participant level. We train a machine learning classifier (support vector machine, SVM) using the subject-wise patterns of networks’ relative contributions to FC from controls, DOC patients, and anaesthesia (including pre-anaesthetic baseline and anaesthesia, but excluding post-anaesthetic recovery). The SVM is trained to distinguish unperturbed baseline from perturbed consciousness, using leave-one-out crossvalidation. This machine learning model achieves 91% accuracy: significantly better performance than chance (*p* < 0.001; permutation test with 5,000 repetitions with shuffled labels; Fig. 4a), comparing favourably with recent DOC classification efforts [63]. The SVM trained on the relative importance of different networks is also significantly better than the performance (79% accuracy) obtained by following exactly the same training regime, but using the univariate correlations between FC and each brain network, instead of the relative importance from dominance analysis (*p* = 0.002; *χ*^2^ = 9.39 from McNemar test) (Fig. S5).

**Figure 4.**
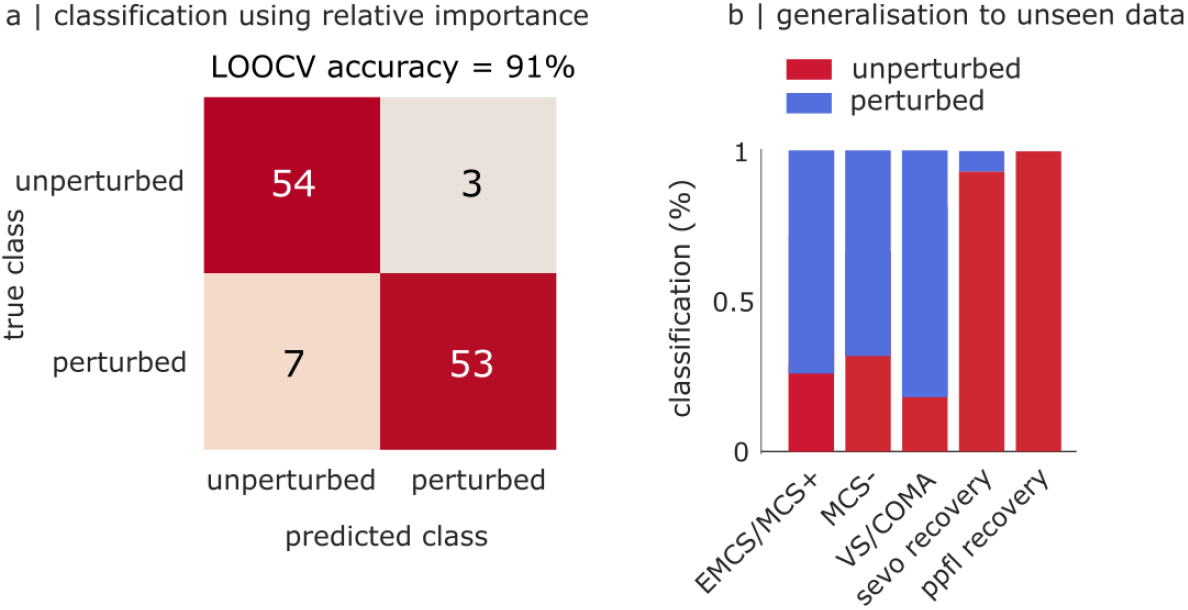
Machine learning classification of consciousness from multimodal network contributions. **(a)** Confusion matrix for the classification of pharmacological and pathological perturbations of consciousness using the relative importance of multimodal networks, achieving 91% accuracy (p < 0.001) after training a support vector machine using leave-one-out cross-validation. **(b)** Classification of unseen data: the recovery scans from the sevoflurane and deep propofol anaesthesia datasets; and three different groups of DOC patients from the Paris dataset: emerged from minimally conscious (EMCS) and MCS+; MCS-; and coma and vegetative state (also known as unresponsive wakefulness syndrome).

Finally, we consider out-of-sample validation for the model. First, we use the trained SVM to classify the recovery data from the sevoflurane and propofol datasets, which were not included in the data used to train the SVM. In all but one case, individuals (who were all responsive at this stage) are correctly assigned to unperturbed consciousness, despite the lingering presence of anaesthetic in their bloodstream (Fig. 4b). Then, we use the same SVM to classify an independent sample of DOC patients (Paris dataset, N=51 patients). The group comprises a broader range of DOC patients, including not only vegetative state/unresponsive wakefulness syndrome and minimally conscious state (further subdivided into MCS ‘plus’ and ‘minus’), like the Cambridge dataset, but also coma and emergence from MCS (EMCS). We find that using the patients’ patterns of relative importance from dominance analysis, the SVM classifies 71% (10*/*14) of the EMCS and MCS+ patients as being in a state of perturbed consciousness (i.e., more similar to anaesthetised than to awake individuals, and therefore presumed unconscious), and similarly 66% (10*/*15) of the MCS-, reaching an even higher proportion (81%, or 18*/*22) for the VS/coma patients, whose disorder is most severe. Further complementing this approach of binary classification, we apply multiple regression to predict the Paris DOC patients’ Glasgow Outcome Scale - Extended (GOSE) scores at 6 months. We find that the subject-wise patterns of networks’ relative contributions to fMRI FC can account for 34% of the variance in DOC patients’ GOSE scores after 6 months, significantly more than would be expected by chance (*p* = 0.011; Fig. S6).

### Cross-modal validation with causal functional circuits from electrical stimulation

The network of meta-analytic co-activations reflects which regions are consistently recruited by the same cognitive operations: it identifies the macroscale circuits that support different facets of cognition in the human brain, indicating similarity of function. However, it is based on results from functional MRI, the same modality used to quantify the prediction target of haemodynamic connectivity. Nonetheless, it should be noted that NeuroSynth is not based on resting-state fMRI, but rather task-fMRI. In fact, the NeuroSynth meta-analytic maps are not directly computed from any task-fMRI data; rather, each map is obtained by aggregating the co-occurrence of terms and coordinates across articles in the published literature [65]. Hence the extent to which haemodynamic FC and NeuroSynth meta-analytic co-activation are ‘the same modality’ is at best indirect.

To corroborate this theoretical argument with empirical data, and dispel the possibility that the dominance of cognitive co-activations may be driven by use of the same imaging technique, we replicate our main analysis using a different, complementary strategy to quantify similarity of function: elicitation of similar subjective experiences in response to direct intracranial electrical stimulation, across a large sample of patients undergoing pre-surgical mapping [73]. Experiences elicited by direct electrical stimulation were grouped into eight categories: somatomotor, visual, olfactory, vestibular, emotion, language, memory, and physiological (Fig. 5a). Different regions elicit these categories with different probabilities upon direct electrical stimulation, and we obtain a co-elicitation network by correlating regions’ profiles of elicitation probabilities across the eight categories (Fig. 5a).

**Figure 5.**
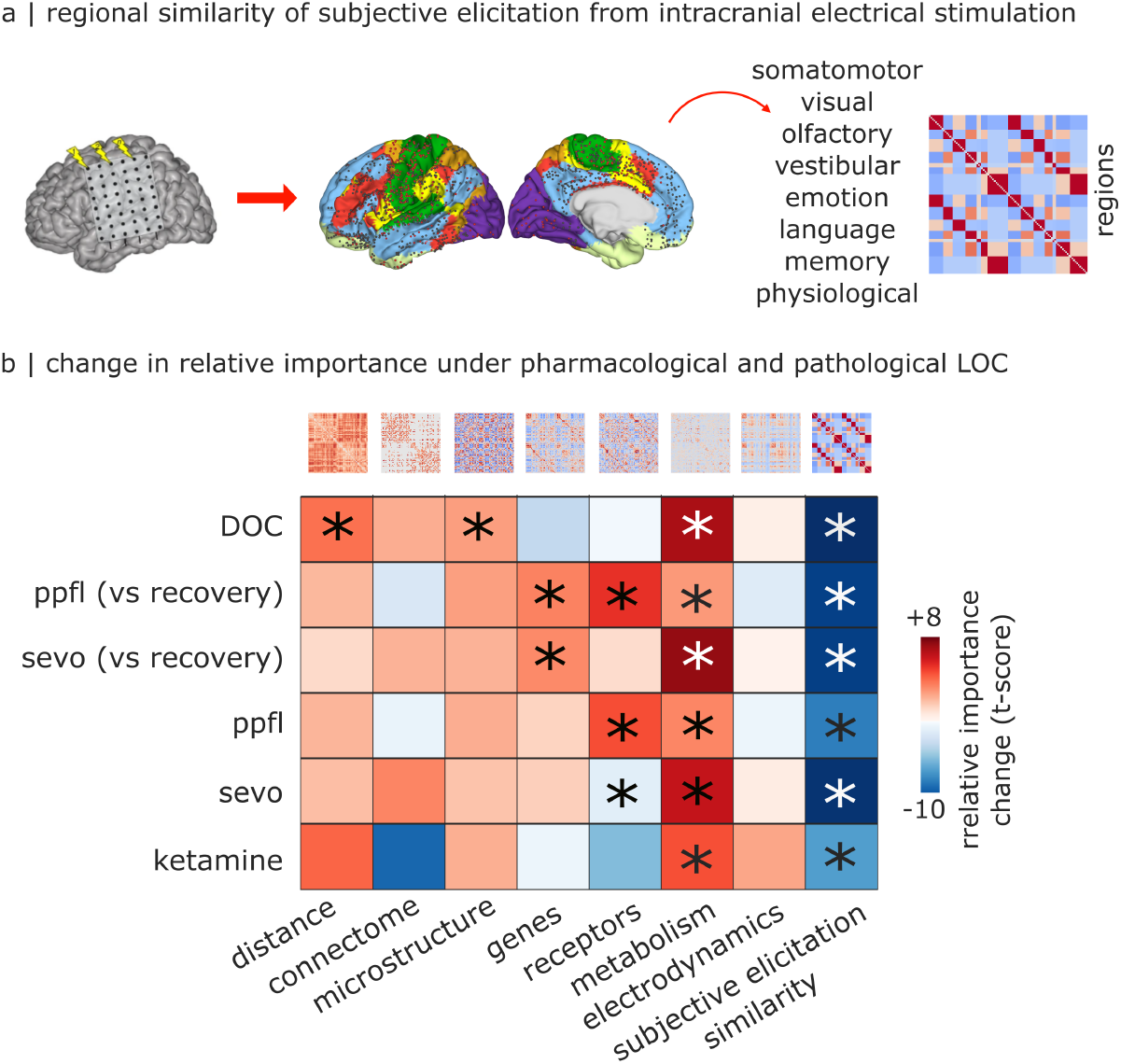
Reduced contribution of meta-analytic co-activation to functional connectivity is replicated with the network of similarity of subjective experiences elicited by intracranial electrical stimulation. *, p < 0.05 with head motion included as covariate of no interest. See Tables S3-S4 for statistical reporting. Red color-scale indicates altered > baseline. Blue color-scale indicates baseline > altered.

This co-elicitation approach is complementary to meta-analytic co-activation in several respects. First, co-elicitation is based on similarity of effects across direct experimental manipulations. Second, the direction of inference between brain regions and cognition is different. Instead of starting from a cognitive operation (i.e., a term for the term-based meta-analysis) and asking which regions co-activate in studies that mention this term, co-elicitation starts from an area of cortex and asks which subjective experiences it elicits upon direct intracranial electrical stimulation. Third, the modality is different: co-elicitation is quantified based on subjective reports following intracranial electrical stimulation, rather than activation from functional MRI tasks. Fourth, co-elicitation does not rely on meta-analytic data-aggregation, but rather it is based on the results of a single study from hundreds of patients–thereby providing stronger experimental confidence [73, 74].

Despite all these differences with the meta-analytic approach, use of electrical co-elicitation reproduces our main findings. First, we find that when defined in terms of subjective co-elicitation, the network of cognitive similarity accounts for 26-37% of the variance in functional connectivity, thereby remaining the predictor with highest relative importance (Fig. S7). Second, we also find that the relative importance of the network of cognitive similarity is significantly reduced under pathological and pharmacological loss of consciousness (Fig. 5b), as low as 6% in the case of DOC patients (Fig. S7). Also in accordance with the main analysis, use of the networks’ distribution of relative importances from dominance analysis allows for high classification accuracy (89%) of conscious versus unconscious individuals (Fig. S8). Altogether, similarity of subjective experiences elicited by direct intracranial stimulation reflects regions’ shared cognitive roles, without use of fMRI. This analysis therefore allowed us to disentangle whether the network of metaanalytic co-activations predicts haemodynamic FC due to its cognitive relevance, or merely due to the shared modality of fMRI. We found that what matters is the similarity of regions’ cognitive roles, not the modality through which they are assessed.

### Robustness and sensitivity

We replicate the main results using a different parcellation (the Desikan-Killiany atlas [75], which is defined anatomically instead of functionally). We find consistent results for the predominant importance of metaanalytic co-activation for predicting haemodynamic FC, and how it is affected by anaesthesia and DOC (Fig. S9 and Fig. S10). Indeed, there is a significant correlation between the changes in relative importance observed in the main analysis, and in this replication (Spearman *ρ* = 0.70, *p* < 0.001).

Using the same anatomical atlas, we also replicate the main results with a different method for quantifying meta-analytic co-activation: using the expert-curated BrainMap database [12, 76], instead of NeuroSynth which is an automated tool (Fig. S11). We find that this alternative meta-analytic approach is less powerful than NeuroSynth at recapitulating FC in the awake healthy brain, accounting for only 25 − 30% of the variance explained (compared with 45 − 55% for NeuroSynth, for both Schaefer and Desikan-Killiany atlases)—possibly due to the smaller number of studies included, being a manually-curated database (Fig. S13). Nevertheless, meta-analytic co-activation remains the most important predictor of haemodynamic FC at baseline, and using BrainMap we again find that perturbations of consciousness induce prominent reductions in the contribution (relative importance) of cognitively-relevant coactivation to FC (Fig. S12). This effect fails to meet statistical significance for the ketamine anaesthesia dataset and for recovery-versus-propofol, after controlling for head motion. We also observe that the counterbalancing increase is primarily observed for gene expression similarity, rather than receptor expression or metabolic similarity, which is what we had observed in the main results. Despite these minor discrepancies, we find a significant positive correlation between the changes in relative importance observed in the main analysis with NeuroSynth, and in the BrainMap replication (Spearman *ρ* = 0.63, *p* < 0.001).

We also replicate the main results using gene expression data obtained from RNA-seq rather than microarray. The relative importance of gene co-expression for explaining FC in the awake brain is similar across the two modalities, around 7-10% (Fig. S14). Consistent with the main results using microarray data, we also find that during unconsciousness, the contribution of metaanalytic co-activation to FC is significantly and consistently reduced, and there is instead an increased contribution of gene co-expression and to some extent receptor co-expression and metabolic similarity (Fig. S15). Indeed, we find a significant correlation between the changes in relative importance observed in the main analysis with microarray, and in the RNA-seq replication (Spearman *ρ* = 0.79, *p* < 0.001).

As a further robustness check, we replicate the main results using an alternative approach for revealing the anatomical connectivity between regions: instead of diffusion tractography (which is non-invasive but also not causal), we use the amplitude of cortico-cortical evoked potentials between pairs of regions, as estimated from *>* 34 354 direct electrical stimulations in 780 patients with epilepsy from the F-TRACT database [77] (Fig. S16). The rationale for this approach is that, if direct electrical stimulation in region *i* induces an electrical evoked potential in region *j* (as recorded by stereo-EEG), then this is causal evidence that region *i* is directly or indirectly connected to region *j*. Being predicated on a region’s capacity to exert causal influence on another, this method arguably reflects effective connectivity, as opposed to structural connectivity [78]. Although this method of estimating anatomical connectivity between pairs of regions is conceptually very different from non-invasive diffusion tractography, we find highly consistent results between the two, including a predominant role of the NeuroSynth co-activation at baseline (Fig. S17), and a reduction of its relative importance under loss of consciousness (Fig. S18). The consistency of results is confirmed quantitatively by the significant positive correlation between the changes in relative importance observed in the main analysis with diffusion tractography, and in the F-TRACT replication (Spearman *ρ* = 0.63, *p* < 0.001).

We also show that cognitively-relevant co-activation remains consistently the most dominant predictor of haemodynamic FC in the healthy brain, when only specific frequency bands are used for computing electrodynamic connectivity from MEG: delta (2–4 Hz), theta (5– 7 Hz), alpha (8–12 Hz), beta (15–29 Hz), low gamma (30–59 Hz), and high gamma (60–90 Hz) (Fig. S19). Results exhibit some frequency-specificity: MEG connectivity becomes the second-best predictor when alpha-band is considered, narrowly outperforming molecular architecture and anatomical connectivity. It may seem surprising that the highest performance of MEG connectivity for predicting haemodynamic FC is observed for frequencies in the alpha rather than delta band, which is the slowest (2-4Hz) and therefore closest to the timescale of fMRI. However, it was already shown at the univariate level that the correspondence between fMRI and MEG functional connectivity does not peak in the delta-band [45], which the present multivariate results corroborate.

Through the analysis of subjective experiences elicited by direct stimulation, we demonstrated that the capacity of meta-analytic co-activation to predict coordinated haemodynamics is not due to the shared modality (fMRI), which direct intracranial stimulation does not share. Rather, what matters is the similarity of regions’ cognitive roles, which NeuroSynth co-activation and direct intracranial stimulation both capture, through very different avenues. For our final analysis, we show that the converse also holds: shared modality alone is not the main predictor of how different regions coordinate their haemodynamic activity. To do so, we replicate results after adding an additional predictor network: similarity of blood perfusion from Arterial Spin Labelling (ASL). Although the BOLD signal is based on haemodynamics, which is closely related to blood perfusion, we find that co-perfusion is only the second-best predictor, accounting for approximately 25% of explained variance in fMRI FC (Fig. S20). In contrast, meta-analytic cognitive coactivation remains by far the most important predictor of awake BOLD FC, accounting for 40 − 45% of the explained variance (Fig. S20). As with the main analysis, meta-analytic co-activation is also the predictor that is most affected by anaesthesia and DOC, occurring consistently in each dataset, even when similarity of blood perfusion is included (Fig. S21).

Altogether, the present findings are observed across multiple datasets (propofol, sevoflurane, and ketamine anaesthesia); they remain stable across test-retest; and they generalise to an unseen cohort of DOC patients. Results are neither driven by head motion, nor by simple Euclidean distance between parcels. We also provide evidence that results are robust to the choice of parcellation (anatomical or functional), to the use of a different gene expression database (RNA-seq), to the choice of neurophysiological frequency band, and to the use of a different method for connectome reconstruction (corticocortical evoked potentials from direct electrical stimulation). We further demonstrate that haemodynamic FC in the awake brain is better captured by meta-analytic coactivation across cognitive operations, than by the mere similarity of regions’ blood perfusion. Finally, the finding of reduced contribution from the network of cognitively-relevant co-activation replicates not only when using an alternative meta-analytic approach (BrainMap), but also when defining cognitive similarity in terms of similar subjective experiences elicited by direct intracranial electrical stimulation: bypassing the use of fMRI altogether, and ensuring that our results are not driven by any modality bias but rather by the cognitive similarity that NeuroSynth and intracranial stimulation both capture.

## DISCUSSION

Here, we investigated the relationship between spontaneous haemodynamic networks (functional connectivity) and other anatomical and functional networks [24], in the awake healthy brain and across pharmacological and pathological perturbations of consciousness. The present findings are two-fold. First, we show that ongoing cognition is the dominant predictor of how brain regions spontaneously synchronise on a timescale of seconds. The network of cognitively relevant co-activations from NeuroSynth meta-analysis is by far the main contributor for predicting the fMRI functional connectivity of the human brain. While the structural connectome and molecular architecture of genes and receptors also make notable contributions in the order of 10% each, in line with their well established univariate relationships with FC [26, 27, 36, 51, 79], nonetheless cognitive co-activation alone accounts for equal or more variance than all the other seven structural and functional networks combined. This finding is reproducible across all datasets, and it is also highly stable over time. Second, we find that the predominance of cognitively-relevant coactivations is systematically reduced across pathological and pharmacological perturbations of consciousness, including disorders of consciousness (due to cardiac arrest or traumatic injury), but also anaesthesia with propofol, sevoflurane, or a high dose of ketamine that rendered participants unresponsive and disconnected from the environment. This effect is consistently accompanied by increased contributions from networks of metabolic co-fluctuation and gene and receptor expression similarity. The result is a coherent pattern of increases and decreases in the relative contributions of structural and functional networks to FC.

It is an implicit assumption underpinning rs-fMRI research that synchronous fluctuations should indicate that two regions are involved in the same cognitive functions: hence the name of ‘functional’ connectivity. This assumption found support in early studies indicating that even at ‘rest’, FC is not random, instead recapitulating task-induced co-activations [11, 13–15]. However, it is also widely observed that FC retains some structure even during sleep or anaesthesia, when cognition is absent, and is partly determined by the relative distance and physical connectivity between regions, as well as their molecular and cytoarchitectonic profiles [26, 27]. Previous studies have not directly compared these different accounts of FC.

Here, we used dominance analysis to arbitrate between these two contraposed accounts of haemodynamic functional connectivity. Results consistently indicate that the network of meta-analytic co-activations is the most dominant predictor of FC—eclipsing the contributions of spatial distance or even physical connectivity, and also blood perfusion (which is measured using the same imaging modality). The network of meta-analytic coactivations reflects which regions are consistently involved in the same cognitive operations—such as attention, memory, emotion or language. It identifies the macroscale circuits that support different facets of cognition in the human brain, indicating sameness of function. In other words, to know whether regions *i* and *j* will exhibit synchronous haemodynamics, the most relevant piece of information is how consistently *i* and *j* are jointly recruited across the spectrum of human cognition. While dominance analysis does not establish that ongoing cognition causally determines FC, the present report indicates that regions’ cognitive co-activation is up to 10 times more important as a predictor of their FC than knowing how close the two regions are in space, and up to 5 times more important than knowing whether they are directly monosynaptically connected.

Notably, the predictors of coordinated haemodynamics included coordinated electrodynamics from MEG [45], and coordinated metabolic activity from FDG-PET [41]. One might have expected these modalities to be the most relevant predictors of BOLD FC, because they reflect the same kind of process: synchronous activity over time, in the absence of any task. However, it is worth noting that the three modalities differ substantially not only in terms of what signals they reflect, but also in terms of temporal resolution, spanning several orders of magnitude: from millisecond (MEG neurophysiology) to 1-2s (fMRI haemodynamics) and approximately 15s (glucose consumption from FDG-PET). Among them, we speculate that fMRI may occupy a privileged timescale for capturing spontaneous cognition in the resting state. Intuitively, one’s internal monologue is certainly faster than FDG-PET’s 15s, but for many purposes it may be slower than the millisecond-level timescale where M/EEG excels. Indeed, the present results remained robust regardless of the chosen MEG frequency. This line of argument suggests that although electrophysiology remains the undisputed gold standard for capturing the brain’s reactions to external stimuli, and for examining the broadest range of temporal scales [80], fMRI may be well-suited to capturing ongoing, internally-driven cognition. This would be consistent with the present evidence that haemodynamic FC bears little resemblance to either electrodynamic or metabolic similarity after other modalities have been accounted for, and is instead closely related to regions’ co-involvement across cognitive functions.

Crucially, we also replicated results about cognitive co-activations using an entirely different data modality: similarity of subjective experiences elicited by direct intracranial electrical stimulation [73], as opposed to meta-analytic aggregation from the fMRI literature. This drastically different approach provided fully consistent results with the main NeuroSynth analysis, demonstrating that the present results do not pertain to a specific modality (fMRI) but rather to regions’ joint recruitment across cognitive operations. Convergent evidence comes from the control analysis with blood perfusion, which is outperformed by meta-analytic co-activation as a predictor of haemodynamic FC.

Although a decrease in the relative contribution of cognitively-relevant co-activations necessitates some increase in the relative contributions of other predictors, such increase could take many forms. For example, different predictors could increase their contributions heterogeneously across individuals, which would not produce a statistically significant increase at the group level. Or a different predictor could increase its contribution in each dataset, which would produce a statistically significant increase, but without manifesting as a consistent pattern across different datasets. Alternatively, each predictor could increase its contribution in approximately equal amounts. Again such a scenario would not be in line with the present observation that increases are both statistically significant within datasets, and consistent across datasets in terms of which predictors are involved. Therefore, the present results of consistent increases in the contribution of specific networks cannot be simply discounted as a side-effect of the reduced contribution of NeuroSynth meta-analytic patterns.

The present account of functional connectivity being shaped by ongoing cognition is further corroborated by the results that we provide about anaesthesia and disorders of consciousness. Both anaesthesia and DOC are characterised not only by behavioural unresponsiveness, but also by either the complete cessation of cognition due to unconsciousness, or its disconnection from the goings-on in the external environment. In each of these cases, brain regions cease to coordinate their activity in a manner that is consistent with the cognitive operations of the healthy brain. Instead, we find greater contributions from SC and molecular architecture (genes and receptors), as well as metabolic similarity. The exception is ketamine, where in accordance with previous findings [71] we do not observe an increased contribution of SC– possibly because participants’ consciousness was disconnected from the environment, but not altogether abolished. The traditional account for the increased coupling between FC and SC observed under loss of consciousness [17, 18, 21, 81] is that the brain loses the flexibility to visit functional configurations beyond the dictates of anatomical connectivity. The present results add an important caveat. It is not just *any* functional configuration that becomes less accessible: rather, the brain ceases to spontaneously self-organise according to *cognitively relevant* functional patterns.

In addition to this decoupling between FC and cognition, we also see that increased structure-function coupling between SC and FC is not the full story. Rather, multiple additional aspects of brain anatomy increase their hold on FC. Specifically, we consistently observed greater contributions of gene and receptor co-expression. Neurotransmitter receptors govern the responsiveness of neurons to neuromodulatory influences, and gene expression shapes the cellular and molecular composition of each region. Together, these properties are major determinants of local circuit dynamics. It is therefore unsurprising that regions with similar local circuitry, attuned to the same neuromodulatory influences, should exhibit time-locked fluctuations in their activity. Indeed, under perturbed consciousness spontaneous haemodynamics also become more in line with the similarity of regions’ metabolic demands, which are likely to be largely driven by intrinsic cytoarchitecture, in the absence of specific cognitive demands imposed on a region.

Altogether, by considering a broader range of structural and functional networks across different perturbations of consciousness we can see that increased coupling with the SC is indeed observed (except for ketamine anaesthesia) in accordance with prior literature, but it is not the full story. Rather, contextualising BOLD FC with respect to multimodal anatomical and functional networks of the human brain reveals additional, consistent reconfigurations across pathological and pharmacological perturbations of consciousness. The present approach complements recent efforts that used multiple modalities for each DOC patient, including functional and diffusion MRI but also FDG-PET and electrophysiology, showing the added diagnostic and prognostic value of additional modalities [63]. Here we used instead a single modality per individual (fMRI), using multiple modalities to interpret it. Future work may seek to combine the two approaches, using patient-specific multimodal networks to characterise patient-specific BOLD FC.

There are a number of limitations that should be borne in mind when interpreting the results. All networks are obtained from different populations of varying ages, sex ratios, handedness, and sample size (e.g., only 6 donors for gene expression [52], and one donor for histology [62])—which is inevitable since the histology and gene expression datasets are both post-mortem. Subcortical regions are not included in the present analysis, due to notable cortical-subcortical differences for the receptor data [38], and outright unavailability for the electrical stimulation and electrophysiology datasets [45, 73]. However, future work should seek ways to extend the present results to the subcortex, since structures such as the thalamus, basal ganglia, and brainstem all play prominent roles in the mechanisms of anaesthesia and disorders of consciousness [82–92]. Each dataset also comes with its own specific limitations including instances of false positives and negatives in diffusion tractography [93], nonspecific binding for some PET tracers [94], and heterogeneous levels of signal-to-noise ratio across all imaging types [24]. Spatial and temporal resolutions also vary. Results may therefore be influenced by differences in how the data are acquired. The functionally defined Schaefer atlas [95] used for all main analyses may reflect functional networks (e.g., metabolic connectivity, electrophysiological connectivity, meta-analytic activation from fMRI) better than anatomical and molecular networks. We sought to mitigate these challenges by running extensive sensitivity analyses, including use of a different parcellation (Desikan-Killiany [75], which is anatomically defined). We also replicated our results using different data modalities: instead of automated meta-analysis of task-fMRI studies with NeuroSynth, we used expert-curated meta-analysis with BrainMap [96], and an entirely different modality, intracranial electrical stimulation [73]; instead of microarray gene expression we used RNA-seq [52]; and instead of diffusion tractography for revealing anatomical connections, we used cortico-cortical evoked potentials from the F-TRACT database [77]. The anaesthesia and DOC datasets are also heterogeneous, being acquired at different sites and with different acquisition parameters [66–68]. We sought to mitigate this concern by using uniform procedures for preprocessing and denoising, to ensure consistency. The fact that we found consistent results despite this heterogeneity is also a testament to the robustness of the present findings.

Altogether, this work combined multimodal and multiscale perspectives on structure-function relationships between cortical connectivity networks, including structural connectivity and geometry but also gene expression, receptor density, cytoarchitecture, electrodynamics at different frequencies, slow metabolic expenditure, and meta-analytic activation of cognitive patterns. We demonstrated that ongoing cognition is the dominant predictor of inter-regional fMRI synchrony at rest, and that the contribution of cognitive co-activations to FC is systematically reduced when cognition is abolished by pathological and pharmacological perturbations of consciousness.

## MATERIALS AND METHODS

### Haemodynamic connectivity datasets

#### Sevoflurane anaesthesia dataset

The sevoflurane data included here have been published before [67], and we refer the reader to the original publication for details. The ethics committee of the medical school of the Technische Universitat Munchen (Munchen, Germany) approved the current study, which was conducted in accordance with the Declaration of Helsinki. Written informed consent was obtained from volunteers at least 48 h before the study session. Twenty healthy adult men (20 to 36 years of age; mean, 26 years) were recruited through campus notices and personal contact, and compensated for their participation in the study. Before inclusion in the study, detailed information was provided about the protocol and risks, and medical history was reviewed to assess any previous neurologic or psychiatric disorder. A focused physical examination was performed, and a resting electrocardiogram was recorded. Further exclusion criteria were the following: physical status other than American Society of Anesthesiologists physical status I, chronic intake of medication or drugs, hardness of hearing or deafness, absence of fluency in German, known or suspected disposition to malignant hyperthermia, acute hepatic porphyria, history of halothane hepatitis, obesity with a body mass index more than 30 kg/m2, gastrointestinal disorders with a disposition for gastroesophageal regurgitation, known or suspected difficult airway, and presence of metal implants. Data acquisition took place between June and December 2013.

Sevoflurane concentrations were chosen so that participants tolerated artificial ventilation (reached at 2.0 vol%) and that burst-suppression (BS) was reached in all participants (around 4.4 vol%). To make group comparisons feasible, an intermediate concentration of 3.0 vol% was also used. In the MRI scanner, participants were in a resting state with eyes closed for 700s. Since EEG data were simultaneously acquired during MRI scanning [67] (though they are not analysed in the present study), visual online inspection of the EEG was used to verify that participants did not fall asleep during the pre-anaesthesia baseline scan. Sevoflurane mixed with oxygen was administered via a tight-fitting facemask using an fMRI-compatible anaesthesia machine (Fabius Tiro, Drager, Germany). Standard American Society of Anesthesiologists monitoring was performed: concentrations of sevoflurane, oxygen and carbon dioxide, were monitored using a cardiorespiratory monitor (DatexaS, General electric, USA). After administering an end-tidal sevoflurane concentration (etSev) of 0.4 vol% for 5 min, sevoflurane concentration was increased in a stepwise fashion by 0.2 vol% every 3 min until the participant became unconscious, as judged by the loss of responsiveness (LOR) to the repeatedly spoken command “squeeze my hand” two consecutive times. Sevoflurane concentration was then increased to reach an end-tidal concentration of approximately 3 vol%. When clinically indicated, ventilation was managed by the physician and a laryngeal mask suitable for fMRI (I-gel, Intersurgical, United Kingdom) was inserted. The fraction of inspired oxygen was then set at 0.8, and mechanical ventilation was adjusted to maintain end-tidal carbon dioxide at steady concentrations of 33±1.71 mmHg during BS, 34±1.12 mmHg during 3 vol%, and 33±1.49 mmHg during 2 vol% (throughout this article, mean ± SD). Norepinephrine was given by continuous infusion (0.1±0.01*µg* kg-1 min-1) through an intravenous catheter in a vein on the dorsum of the hand, to maintain the mean arterial blood pressure close to baseline values (baseline, 96±9.36 mmHg; BS, 88±7.55 mmHg; 3 vol%, 88±8.4 mmHg; 2 vol%, 89±9.37 mmHg; follow-up, 98±9.41 mmHg). After insertion of the laryngeal mask airway, sevoflurane concentration was gradually increased until the EEG showed burst-suppression with suppression periods of at least 1,000 ms and about 50% suppression of electrical activity (reached at 4.34±0.22 vol%), which is characteristic of deep anaesthesia. At that point, another 700s of electroencephalogram and fMRI was recorded. Further 700s of data were acquired at steady end-tidal sevoflurane concentrations of 3 and 2 vol%, respectively (corresponding to Ramsay scale level 6, the deepest), each after an equilibration time of 15 min. In a final step, etSev was reduced to two times the concentration at LOR. However, most of the participants moved or did not tolerate the laryngeal mask any more under this condition: therefore, this stage was not included in the analysis. Sevoflurane administration was then terminated, and the scanner table was slid out of the MRI scanner to monitor post-anaesthetic recovery. The participants were manually ventilated until spontaneous ventilation returned. The laryngeal mask was removed as soon as the patient opened his mouth on command. The physician regularly asked the participant to squeeze their hand: recovery of responsiveness was noted to occur as soon as the command was followed. Fifteen minutes after the time of recovery of responsiveness, the Brice interview was administered to assess for awareness during sevoflurane exposure; the interview was repeated on the phone the next day. After a total of 45 min of recovery time, another resting-state combined fMRI-EEG scan was acquired (with eyes closed, as for the baseline scan). When participants were alert, oriented, cooperative, and physiologically stable, they were taken home by a family member or a friend appointed in advance.

Although the original study acquired both functional MRI (fMRI) and electroencephalographic (EEG) data, in the present work we only considered the fMRI data. Data acquisition was carried out on a 3-Tesla magnetic resonance imaging scanner (Achieva Quasar Dual 3.0T 16CH, The Netherlands) with an eight-channel, phased-array head coil. The data were collected using a gradient echo planar imaging sequence (echo time = 30 ms, repetition time (TR) = 1.838 s, flip angle = 75 deg, field of view = 220×220 mm2, matrix = 72 × 72, 32 slices, slice thickness = 3 mm, and 1 mm interslice gap; 700-s acquisition time, resulting in 350 functional volumes). The anatomical scan was acquired before the functional scan using a T1-weighted MPRAGE sequence with 240 × 240 × 170 voxels (1 1 1 mm voxel size) covering the whole brain. A total of 16 volunteers completed the full protocol and were included in our analyses; one participant was excluded due to high motion, leaving N=15 for analysis.

#### Propofol anaesthesia dataset

The propofol data were collected between May and November 2014 at the Robarts Research Institute, Western University, London, Ontario (Canada), and have been published before [66, 97]. The study received ethical approval from the Health Sciences Research Ethics Board and Psychology Research Ethics Board of Western University (Ontario, Canada). Healthy volunteers (n=19) were recruited (18–40 years; 13 males). Volunteers were right-handed, native English speakers, and had no history of neurological disorders. In accordance with relevant ethical guidelines, each volunteer provided written informed consent, and received monetary compensation for their time. Due to equipment malfunction or physiological impediments to anaesthesia in the scanner, data from n=3 participants (1 male) were excluded from analyses, leaving a total n=16 for analysis.

Resting-state fMRI data were acquired at different propofol levels: no sedation (Awake), Deep anaesthesia (corresponding to Ramsay score of 5) and also during post-anaesthetic recovery. As previously reported [66], for each condition fMRI acquisition began after two anaesthesiologists and one anaesthesia nurse independently assessed Ramsay level in the scanning room. The anaesthesiologists and the anaesthesia nurse could not be blinded to experimental condition, since part of their role involved determining the participants’ level of anaesthesia. Note that the Ramsay score is designed for critical care patients, and therefore participants did not receive a score during the Awake condition before propofol administration: rather, they were required to be fully awake, alert and communicating appropriately. To provide a further, independent evaluation of participants’ level of responsiveness, they were asked to perform two tasks: a test of verbal memory recall, and a computer-based auditory target-detection task. Wakefulness was also monitored using an infrared camera placed inside the scanner. Propofol was administered intravenously using an AS50 auto syringe infusion pump (Baxter Healthcare, Singapore); an effect-site/plasma steering algorithm combined with the computer-controlled infusion pump was used to achieve step-wise sedation increments, followed by manual adjustments as required to reach the desired target concentrations of propofol according to the TIVA Trainer (European Society for Intravenous Anaeesthesia, eurosiva.eu) pharmacokinetic simulation program. This software also specified the blood concentrations of propofol, following the Marsh 3-compartment model, which were used as targets for the pharmacokinetic model providing target-controlled infusion. After an initial propofol target effect-site concentration of 0.6 *µ*g mL^−1^, concentration was gradually increased by increments of 0.3 *µ*g mL^−1^, and Ramsay score was assessed after each increment: a further increment occurred if the Ramsay score was lower than 5. The mean estimated effect-site and plasma propofol concentrations were kept stable by the pharmacokinetic model delivered via the TIVA Trainer infusion pump. Ramsay level 5 was achieved when participants stopped responding to verbal commands, were unable to engage in conversation, and were rousable only to physical stimulation. Once both anaesthesiologists and the anaesthesia nurse all agreed that Ramsay sedation level 5 had been reached, and participants stopped responding to both tasks, data acquisition was initiated. The mean estimated effect-site propofol concentration was 2.48 (1.82-3.14) *µ*g mL-1, and the mean estimated plasma propofol concentration was 2.68 (1.92-3.44) *µ*g mL-1. Mean total mass of propofol administered was 486.58 (373.30-599.86) mg. These values of variability are typical for the pharmacokinetics and pharmacodynamics of propofol. Oxygen was titrated to maintain SpO2 above 96%. At Ramsay 5 level, participants remained capable of spontaneous cardiovascular function and ventilation. However, the sedation procedure did not take place in a hospital setting; therefore, intubation during scanning could not be used to ensure airway security during scanning. Consequently, although two anaesthesiologists closely monitored each participant, scanner time was minimised to ensure return to normal breathing following deep sedation. No state changes or movement were noted during the deep sedation scanning for any of the participants included in the study. Propofol was discontinued following the deep anaesthesia scan, and participants reached level 2 of the Ramsay scale approximately 11 minutes afterwards, as indicated by clear and rapid responses to verbal commands. This corresponds to the “recovery” period. As previously reported [66], once in the scanner participants were instructed to relax with closed eyes, without falling asleep. Resting-state functional MRI in the absence of any tasks was acquired for 8 minutes for each participant. A further scan was also acquired during auditory presentation of a plot-driven story through headphones (5-minute long). Participants were instructed to listen while keeping their eyes closed. The present analysis focuses on the resting-state data only; the story scan data have been published separately [98] and will not be discussed further here.

As previously reported [66], MRI scanning was performed using a 3-Tesla Siemens Tim Trio scanner (32-channel coil), and 256 functional volumes (echo-planar images, EPI) were collected from each participant, with the following parameters: slices = 33, with 25% inter-slice gap; resolution = 3mm isotropic; TR = 2000ms; TE = 30ms; flip angle = 75 degrees; matrix size = 64 × 64. The order of acquisition was interleaved, bottomup. Anatomical scanning was also performed, acquiring a high-resolution T1-weighted volume (32-channel coil, 1mm isotropic voxel size) with a 3D MPRAGE sequence, using the following parameters: TA = 5min, TE = 4.25ms, 240 × 256 matrix size, 9 degrees flip angle [66].

#### Ketamine anaesthesia dataset

The anaesthetic ketamine data used here have been published previously [68], and we refer the reader to the original publication for further details. The original study was approved by the ethics committee of the Medical School of the University of Liège (University Hospital, Liège, Belgium) and registered at EudraCT (2010-023016-13). Fourteen right-handed volunteers were recruited via advertisements on an Internet forum (5 women; median age [range], 25 [19–31] years). Each participant provided written informed consent and underwent a medical interview and physical examination prior to participation.

Volunteers were instructed to fast for at least 6 h from solids and 2 h from liquids prior to the experimental session. After acquisition of structural MRI images, participants were removed from the MRI scanner and fitted with 64 scalp electroencephalogram (EEG) electrodes to enable simultaneous EEG–fMRI recording. In the present study, only functional MRI data were analyzed. An 18-gauge intravenous catheter (BD Insyte-W; Becton Dickinson Infusion Therapy Systems Inc., USA) was inserted into a vein of the left forearm and infused with normal saline at a rate of 20 ml/h. This line was used for ketamine infusion and potential administration of rescue medications. Additionally, a 20-gauge arterial catheter (Arrow International Inc., USA) was placed in the left radial artery under strict sterile conditions after local anesthesia with 3 ml of 1% lidocaine. The arterial catheter was connected to a monitoring set (TruWave, Edwards Lifesciences, Dominican Republic) for arterial blood sampling and gas analysis. Standard MRI-compatible anesthesia monitoring equipment (Magnitude 3150M; Invivo Research Inc., USA) was used to continuously record electrocardiogram, heart rate, blood pressure, pulse oximetry (SpO_2_), and respiratory rate throughout scanning and recovery. Supplemental oxygen was delivered at 5 l/min via a loosely fitting plastic facemask, while spontaneous breathing was maintained. One certified anesthesiologist and one neurologist were present throughout the experiment. After all equipment and monitoring devices were set up, volunteers were positioned comfortably in the MRI tray in a supine position to minimize discomfort or painful stimulation. All participants wore earplugs to reduce scanner noise and headphones to allow communication with investigators. One investigator remained in the MRI scan room at all times.

Ketamine was administered using a computer-controlled intravenous infusion system operated via a dedicated laptop computer. A 50-ml syringe was filled with normal saline containing racemic ketamine (Ketalar, Pfizer Ltd., Turkey) at a concentration of 10 mg/ml. The infusion pump was driven by the Domino pharmacokinetic model, which has demonstrated acceptable predictive performance. The system adjusted infusion rates to target precise plasma and effect-site ketamine concentrations based on biometric parameters. Following each change in target concentration, a 5-min equilibration period was allowed to ensure compartmental equilibration. Sedation depth was assessed using the Ramsay Scale (RS) and the University of Michigan Sedation Scale (UMSS), immediately before and after each fMRI acquisition. Volunteers were instructed to squeeze the investigator’s hand forcefully, with the command repeated twice. An investigator remained inside the MRI room throughout the experiment to ensure close monitoring. An initial fMRI acquisition was performed without ketamine infusion (awake state, W1). Ketamine infusion was then initiated, and target concentration was increased in steps of 0.5 *µ*g/ml until light sedation was achieved (RS 3–4 or UMSS 1–2; S1). After the 5-min equilibration period, an identical fMRI acquisition sequence was performed. Target concentration was sub-sequently increased in additional steps of 0.5 *µ*g/ml until deep sedation was reached (RS 5–6 or UMSS 4; S2), followed by another identical acquisition sequence. Due to ketamine’s long elimination half-life and to limit time spent in the scanner, the order of clinical states was not randomized, and a recovery condition was not acquired. After completion of data acquisition, ketamine infusion was stopped and participants were removed from the scanner for recovery. The occurrence of dreaming during ketamine infusion was assessed via a telephone interview conducted after the experimental session [68]. In the present analysis, only the awake and deep sedation conditions were considered.

MRI data were acquired on a 3 T Siemens Allegra scanner (Siemens AG, Germany) using an echo-planar imaging (EPI) sequence with 32 slices (repetition time = 2460 ms; echo time = 40 ms; field of view = 220 mm; voxel size = 3.45 × 3.45 × 3 mm; matrix size = 64 × 64 × 32). For each volunteer and condition, 300 functional volumes were acquired. A high-resolution structural T1-weighted image was obtained at the beginning of the experiment for coregistration with functional data. Six participants were excluded from further analysis due to excessive agitation and movement (5 participants) or voluntary withdrawal (1 participant), leaving data from eight participants for final analysis [68].

#### Cambridge DOC dataset

The disorders of consciousness (DOC) patient data used in this study have been published previously. For clarity and consistency of reporting, where applicable we use the same wording as in our earlier publications using these data [66, 99]. Briefly, 71 patients with DOC were recruited from specialised long-term care centres between January 2010 and December 2015. Ethical approval was provided by the National Research Ethics Service (National Health Service, UK; LREC reference 99/391).

Patients were eligible for inclusion if they had a diagnosis of a chronic disorder of consciousness, written informed consent was obtained from their legal representative, and they could be safely transported to Addenbrooke’s Hospital (Cambridge, UK). Exclusion criteria included any medical condition rendering participation unsafe, as judged by clinical personnel blinded to the specific aims of the study, or any contraindication to the MRI environment (e.g. non-MRI-safe implants). Patients were also excluded if they had significant pre-existing mental health problems or insufficient fluency in English prior to their injury.

Following admission to Addenbrooke’s Hospital, each patient underwent a comprehensive programme of clinical and neuroimaging assessments, including task-based, resting-state, and anatomical MRI scans. Patients spent a total of five days in hospital, including arrival and departure days. Neuroimaging data were acquired at the Wolfson Brain Imaging Centre (Addenbrooke’s Hospital, Cambridge, UK), and all prescribed medications were maintained throughout scanning.

The Coma Recovery Scale–Revised (CRS-R) was administered at least once daily throughout each patient’s admission. Patients who did not exhibit behavioural evidence of awareness at any assessment were classified as being in an Unresponsive Wakefulness Syndrome (UWS). Patients were classified as being in a Minimally Conscious State (MCS) if they demonstrated behavioural evidence of awareness, such as simple automatic motor behaviours (e.g. scratching or pulling bed sheets), visual fixation or pursuit, or localisation to noxious stimulation [100, 101]. Due to the limited sample size, MCS patients were not further subdivided into MCS − and MCS+.

Because the present study focused on whole-brain properties, sufficient brain coverage was required. Consistent with our previous studies [66, 99], patients were excluded prior to analysis if an expert neuroanatomist blinded to diagnosis judged that there was excessive focal brain damage (greater than one third of one hemisphere), if brain damage resulted in suboptimal segmentation or spatial normalisation, or if excessive head motion was observed during scanning (exceeding 3 mm translation or 3^◦^ rotation). Forty-one patients were excluded due to excessive brain damage or distortion preventing satisfactory segmentation and normalisation, and a further eight were excluded due to excessive head motion. The final sample comprised 22 adult patients (14 males, 8 females; age range 17–70 years; mean time post-injury: 13 months), including 10 patients diagnosed with UWS and 12 with MCS due to acquired brain injury [66].

Resting-state fMRI data were acquired for 10 min (300 volumes; TR = 2000 ms) using a Siemens Trio 3 T scanner (Siemens, Erlangen, Germany). Functional images were acquired using an echo-planar imaging (EPI) sequence with 32 slices and the following parameters: voxel size 3 × 3 × 3.75 mm, TR = 2000 ms, TE = 30 ms, and flip angle = 78^◦^. High-resolution anatomical images were acquired using a T1-weighted MPRAGE sequence (TR = 2300 ms; TE = 2.47 ms; 150 slices; voxel size 1 × 1 × 1 mm) [66].

#### Paris DOC dataset

The Pairs DOC dataset has been published before, and we refer to the original articles for full details [18, 63]. This research was approved by the ethical committee of the Pitie-Salpetriere under the French label of ‘routine care research’ (Comité de Protection des Personnes no 2013-A01385-40, Ile de France 1, Paris, France under the code ‘Recherche en soins courants’, protocol numbers NEURO-DoC/HAO-006/20130409 and M-NEURO-DoC/NCT04534777) [18, 63]. Briefly, in this DOC cohort, three MRI modalities were acquired: anatomical T1-weighted MRI (aMRI) and resting-state functional MRI (RS-fMRI), with functional scans lasting approximately five minutes. Two distinct acquisition protocols were used to collect structural and functional images.

In the first protocol, imaging was performed using a 3 T General Electric Signa system (GE HealthCare Technologies Inc., Chicago, IL, USA). Resting-state fMRI data consisted of T2^∗^-weighted whole-brain images acquired using a gradient-echo echo-planar imaging (EPI) sequence in axial orientation (200 volumes, 48 slices, slice thickness = 3 mm, TR/TE = 2400/30 ms, voxel size = 3.4375 × 3.4375 × 3.4375 mm, flip angle = 90^◦^, field of view (FOV) = 220 mm^2^). A T1-weighted anatomical image was acquired during the same session using a magnetization-prepared rapid acquisition gradient echo (MPRAGE) sequence (154 slices, slice thickness = 1.2 mm, TR/TE = 7.112/3.084 ms, voxel size = 1 × 1 × 1 mm, flip angle = 15^◦^). In the second protocol, functional data were acquired using a 3 T Siemens Skyra system (Siemens Healthineers, Erlangen, Germany) with a gradient-echo EPI sequence (180 volumes, 62 slices, slice thickness = 2.5 mm, TR/TE = 2000/30 ms, voxel size = 2 × 2 × 2 mm, flip angle = 90^◦^, FOV = 240 mm^2^, multiband factor = 2). Structural T1-weighted MPRAGE images were also collected in the same session (208 slices, slice thickness = 1.2 mm, TR/TE = 1800/2.35 ms, voxel size = 0.85 × 0.85 × 0.85 mm, flip angle = 8^◦^). Patients who had received medications known to significantly alter vigilance levels (e.g. propofol or benzodiazepines) prior to scanning were excluded from the fMRI analyses [18, 63].

#### Test-retest dataset

The test-retest data included here have been published before [99, 102]. Briefly, right-handed healthy participants (*N* = 22; age range, 19–57 years; mean age, 35.0 years; SD, 11.2; female-to-male ratio, 9:13) were recruited via advertisements in the Cambridge area and received financial compensation for participation. The study was approved by the Cambridgeshire 2 Research Ethics Committee (LREC 08/H0308/246), and all participants provided written informed consent prior to inclusion. Exclusion criteria included a National Adult Reading Test (NART) score < 70, Mini-Mental State Examination (MMSE) score < 23, left-handedness, history of drug or alcohol abuse, history of psychiatric or neurological disorders, contraindications to MRI scanning, use of medications that could affect cognitive performance or that were prescribed for depression, and any physical impairment that could interfere with completion of the experimental procedures. The study comprised two experimental visits separated by 2–4 weeks.

During each visit, resting-state functional MRI data were acquired for 5 min 20 s using a Siemens Trio 3 T scanner (Siemens, Erlangen, Germany). Functional images were collected using an echo-planar imaging (EPI) sequence with the following parameters: repetition time (TR) = 2000 ms; echo time (TE) = 30 ms; flip angle = 78^◦^; field of view (FOV) = 192 × 192 mm^2^; in-plane resolution = 3.0 × 3.0 mm; 32 slices with slice thickness of 3.0 mm and an inter-slice gap of 0.75 mm. A high-resolution three-dimensional structural image was also acquired using a magnetization-prepared rapid acquisition gradient echo (MPRAGE) sequence with the following parameters: TR = 2300 ms; TE = 2.98 ms; flip angle = 9^◦^; FOV = 256 × 256 mm^2^. Task-based fMRI data were additionally collected and have been analyzed previously to address separate experimental questions [99, 102]. Eighteen participants had usable data for both restingstate fMRI sessions and were included in the present analysis.

#### FMRI preprocessing and denoising

For the Cambridge DOC, test-retest, and anaesthesia datasets, we applied a standard preprocessing pipeline in accordance with our previous publications with anaesthesia and DOC data [66]. Preprocessing was performed using the *CONN* toolbox, version 17f (CONN; http://www.nitrc.org/projects/conn) [103], implemented in MATLAB 2016a. The pipeline involved the following steps: removal of the first 10s, to achieve steady-state magnetization; motion correction; slice-timing correction; identification of outlier volumes for subsequent scrubbing by means of the quality assurance/artifact rejection software *art* (http://www.nitrc.org/projects/artifact_detect); normalisation to Montreal Neurological Institute (MNI-152) standard space (2 mm isotropic re-sampling resolution), using the segmented grey matter image from each participant’s T1-weighted anatomical image, together with an a priori grey matter template.

Denoising was also performed using the CONN toolbox, using the same approach as in our previous publications with pharmaco-MRI datasets [66]. Pharmacological agents can induce alterations in physiological parameters (heart rate, breathing rate, motion) or neurovascular coupling. The anatomical CompCor (aCompCor) method removes physiological fluctuations by extracting principal components from regions unlikely to be modulated by neural activity; these components are then included as nuisance regressors [104]. Following this approach, five principal components were extracted from white matter and cerebrospinal fluid signals (using individual tissue masks obtained from the T1-weighted structural MRI images) [103]; and regressed out from the functional data together with six individual-specific realignment parameters (three translations and three rotations) as well as their first-order temporal derivatives; followed by scrubbing of outliers identified by ART, using Ordinary Least Squares regression [103]. Finally, the denoised BOLD signal timeseries were linearly detrended and band-pass filtered to eliminate both low-frequency drift effects and high-frequency noise, thus retaining frequencies between 0.008 and 0.09 Hz. The step of global signal regression (GSR) has received substantial attention in the literature as a denoising method [105–107]. However, recent work has demonstrated that the global signal contains behaviourally relevant information [108] and, crucially, information about states of consciousness, across pharmacological and pathological perturbations [109]. Therefore, in line with ours and others’ previous studies, here we avoided GSR in favour of the aCompCor denoising procedure, which is among those recommended.

Finally, denoised BOLD signals were parcellated into 100 cortical regions-of-interest (ROIs) from the Schaefer atlas [95]. We also replicated our results with the 68-ROI anatomical Desikan-Killiany cortical parcellation [110]. Functional connectivity was estimated for each individual and each condition, as the Pearson correlation between between pairs of denoised and parcellated BOLD timeseries.

The Paris DOC dataset was processed as follows using `fMRIPrep` version 22.0.2. For participants with visible lesions on T1-weighted anatomical images, manual lesion masks were created and provided as inputs to the preprocessing pipeline. For each BOLD run identified per subject (across all tasks and sessions), the following steps were applied. First, a reference volume and its skull-stripped version were generated using a custom `fMRIPrep` procedure. Head-motion parameters relative to the BOLD reference (rigid-body transformation matrices and the corresponding six rotation and translation parameters) were estimated prior to any spatiotemporal filtering using `mcflirt` (FSL 6.0.5.1; [111]). BOLD runs were slice-time corrected to 0.958 s (half of the slice acquisition range, 0–1.92 s) using `3dTshift` from AFNI. The BOLD time series (including slice-time correction when applied) were resampled into their original native space by applying the estimated head-motion correction transforms. These data are referred to as the preprocessed BOLD time series in native space. The BOLD reference image was co-registered to the T1-weighted (T1w) anatomical reference using `mri_coreg` (FreeSurfer), followed by `flirt` (FSL 6.0.5.1) employing the boundary-based registration cost function. Coregistration was configured with six degrees of freedom. Several confounding time series were computed from the preprocessed BOLD data, including framewise displacement (FD), DVARS, and three global signals extracted from cerebrospinal fluid (CSF), white matter (WM), and whole-brain masks. FD and DVARS were computed for each functional run using Nipype implementations following the definitions in [106].

Post-preprocessing steps included regression of 24 head-motion parameters as confounds, removal of the first four volumes, and temporal bandpass filtering (0.01–0.1 Hz). Physiological noise regressors were estimated using component-based noise correction (aCompCor; [104]). Principal components were extracted after high-pass filtering the preprocessed BOLD time series with a discrete cosine filter (128 s cutoff). For aCompCor, probabilistic masks of CSF, WM, and combined CSF+WM were generated in anatomical space. Unlike the original implementation by [104], masks were not eroded in BOLD space. Instead, voxels likely containing gray matter were excluded by subtracting a gray-matter partial-volume mask thresholded at 0.05, ensuring that components were not extracted from voxels containing even a minimal fraction of gray matter. These masks were then resampled into BOLD space and binarized at a threshold of 0.99, consistent with the original implementation. Components were also estimated separately within CSF and WM masks. For each CompCor decomposition, the smallest number of components explaining at least 50% of variance within the corresponding nuisance mask was retained; remaining components were discarded. Head-motion estimates were included in the confounds file, and confound time series derived from motion parameters and global signals were expanded to include temporal derivatives and quadratic terms [112]. Volumes exceeding thresholds of 0.5 mm FD or 1.5 standardized DVARS were flagged as motion outliers. Additional nuisance regressors were estimated using principal component analysis of signal extracted from a thin band of voxels surrounding the brain boundary, following [113].

The denoised BOLD time series were resampled into standard space, yielding preprocessed BOLD data in MNI152NLin2009cAsym space. All resamplings were performed in a single interpolation step by composing all relevant transformations, including head-motion correction, susceptibility distortion correction when available, and co-registrations to anatomical and output spaces. Volumetric resampling was performed using `antsApplyTransforms` (ANTs) with Lanczos interpolation to minimize smoothing effects. Finally, denoised BOLD signals were parcellated into 100 cortical regions-of-interest (ROIs) from the Schaefer atlas [95]. We also replicated our results with the 68-ROI anatomical Desikan-Killiany cortical parcellation [110].

Functional connectivity was estimated for each individual and each condition, as the Pearson correlation between between pairs of denoised and parcellated BOLD timeseries. Pearson correlation is not the only method for obtaining a network of functional connectivity between fMRI time-series (or indeed between electrophysiological or metabolic time-series). On the contrary, hundreds of statistical and information-theoretic relationships exist [10] and may highlight different relationships between regions. Here, we made the pragmatic choice of using Pearson correlation as the most widely used definition of functional connectivity, which also ensures methodological consistency with recent work using the same multi-modal brain networks [24].

### Multimodal anatomical and functional brain networks

#### Metabolic connectivity from dynamic FDG-PET

Metabolic connectivity indexes how similarly two cortical regions metabolize glucose over time and therefore how similarly two cortical regions consume energy [24]. Here we followed the same procedures as in [24]. Briefly, volumetric 4D PET images of [F^18^]-fluordoxyglucose (FDG, a glucose analogue) tracer uptake over time were obtained from Jamadar et al. [114]. Specifically, 26 healthy participants (77% female, 18–23 years old) were recruited from the general population and underwent a 95 minute simultaneous MR-PET scan in a Siemens (Erlangen) Biograph 3-Tesla molecular MR scanner. Participants were positioned supine in the scanner bore with their head in a 16-channel radiofrequency head coil and were instructed to lie as still as possible with eyes open and think of nothing in particular. FDG (average dose 233 MBq) was infused over the course of the scan at a rate of 36 mL/h using a BodyGuard 323 MR-compatible infusion pump (Caesarea Medical Electronics, Caesarea, Israel). Infusion onset was locked to the onset of the PET scan. This data has been validated and analyzed previously in [115, 116].

PET images were reconstructed and preprocessed according to [116]. Specifically, the 5700-second PET time-series for each subject was binned into 356 3D sinogram frames each of 16-second intervals. The attenuation for all required data was corrected via the pseudo-CT method [117]. Ordinary Poisson-Ordered Subset Expectation Maximization algorithm (3 iterations, 21 subsets) with point spread function correction was used to reconstruct 3D volumes from the sinogram frames. The reconstructed DICOM slices were converted to NIFTI format with size 344 × 344 × 127 (voxel size: 2.09 × 2.09 × 2.03 mm^3^) for each volume. A 5 mm FWHM Gaussian postfilter was applied to each 3D volume. All 3D volumes were temporally concatenated to form a 4D (344 × 344 × 127×356) NIFTI volume. A guided motion correction method using simultaneously acquired MRI was applied to correct the motion during the PET scan. 225 16-second volumes were retained commencing for further analyses.

Next, the 225 PET volumes were motion corrected (FSL MCFLIRT [111]) and the mean PET image was brain extracted and used to mask the 4D data. The fPET data were further processed using a spatiotemporal gradient filter to remove the accumulating effect of the radiotracer and other low-frequency components of the signal [118]. Finally, each time point of the PET volumetric time-series were registered to MNI152 template space using Advanced Normalization Tools in Python (ANTSpy, https://github.com/ANTsX/ANTsPy), parcellated to 100 cortical regions of the Schaefer atlas [95] (or 68 cortical regions according to the Desikan-Killiany atlas, for the robustness analyses) and time-series at pairs of cortical regions were correlated (Pearson’s *r*) to construct a metabolic connectivity matrix for each subject. A group-averaged metabolic connectome was obtained by averaging connectivity across subjects.

#### Neurophysiological connectivity from magnetoencephalogrphy

Electrophysiological connectivity was measured using magnetoencephalography (MEG) recordings, which tracks the magnetic field produced by neural currents [24]. Resting state MEG data was acquired for *n* = 33 unrelated healthy young adults (age range 22–35 years) from the Human Connectome Project (S900 release [119]). The data includes resting state scans of approximately 6 minutes long and noise recording for all participants. MEG anatomical data and 3T structural MRI of all participants were also obtained for MEG preprocessing.

The present MEG data was first processed and used by Hansen et al. [24], Shafiei et al. [45]. Resting state MEG data was preprocessed using the opensource software, Brainstorm (https://neuroimage.usc.edu/brainstorm/ [120]), following the online tutorial for the HCP dataset (https://neuroimage.usc.edu/brainstorm/Tutorials/HCP-MEG). MEG recordings were registered to individual structural MRI images before applying the following preprocessing steps. First, notch filters were applied at 60, 120, 180, 240, and 300 Hz, followed by a high-pass filter at 0.3 Hz to remove slow-wave and DC-offset artifacts. Next, bad channels from artifacts (including heartbeats, eye blinks, saccades, muscle movements, and noisy segments) were removed using Signal-Space Projections (SSP).

Pre-processed sensor-level data was used to construct a source estimation on HCP’s fsLR4k cortex surface for each participant. Head models were computed using overlapping spheres and data and noise covariance matrices were estimated from resting state MEG and noise recordings. Linearly constrained minimum variance (LCMV) beamformers were used to obtain the source activity for each participant. Data covariance regularization was performed and the estimated source variance was normalised by the noise covariance matrix to reduce the effect of variable source depth. All eigenvalues smaller than the median eigenvalue of the data covariance matrix were replaced by the median. This helps avoid instability of data covariance inversion caused by the smallest eigenvalues and regularizes the data covariance matrix. Source orientations were constrained to be normal to the cortical surface at each of the 8 000 vertex locations on the cortical surface, then parcellated according to the Schaefer atlas [95] (or Desikan-Killiany atlas, for the robustness analyses).

After preprocessing and parcellating the data, amplitude envelope correlations were performed between time-series at each pair of brain regions, for six canonical frequency bands separately: delta (2–4 Hz), theta (5–7 Hz), alpha (8–12 Hz), beta (15–29 Hz), low gamma (30– 59 Hz), and high gamma (60–90 Hz). Amplitude envelope correlation is applied instead of directly correlating the time-series because of the high sampling rate (2034.5 Hz) of the MEG recordings. An orthogonalization process was applied to correct for the spatial leakage effect by removing all shared zero-lag signals [121]. The composite electrophysiological connectivity matrix is the first principal component of all six connectivity matrices, as per [24].

#### Co-perfusion networks from Arterial Spin Labelling

The blood perfusion data comes from arterial spin labeling (ASL) magnetic resonance imaging (MRI). In this imaging technique, the magnetization of arterial blood in the neck is flipped (or “labeled”) using a train of short block radiofrequency pulses. Images of the brain containing the magnetic “label” are then acquired following a delay that allows the labeled blood to reach and perfuse brain tissue (i.e., post-labeling delay; PLD). In addition, a “control” image with no magnetization inversion is also acquired. The difference between the “label” images and the “control” image is proportional to cerebral blood perfusion and is not affected by the spins. Through this imaging, arterial transit time (ATT) which represents the time it takes for labeled blood to reach brain tissue is also estimated [122, 123].

Here we include ASL data from a total of 1 305 participants in the HCP Lifespan studies (2.0 Release) [124– 126]. Specifically, data from 627 participants (337 females) from HCP-D (5–22 years) and 678 participants (381 females) from HCP-A (36–100 years) are included. All study procedures of the HCP protocol are approved by the Institutional Review Board at Washington University in St. Louis (HCP-D IRB ID#: 201603135; HCP-A IRB ID#: 201603117). All HCP Lifespan imaging data is acquired using a 3.0 Tesla Prisma scanner (Siemens; Erlangen, Germany) and a 32–channel head coil. In HCP Lifespan studies, ASL data acquisition is based on a pseudo-continuous arterial spin labeling (pCASL) and 2D multiband (MB)-echo-planar imaging (EPI) sequence. Pseudo-continuous ASL data are acquired with labeling duration of 1 500 *ms* and five post-labeling delays of 200 *ms*, 700 *ms*, 1 200 *ms*, 1 700 *ms*, and 2 200 *ms*, containing 6, 6, 6, 10, 15 control-label image pairs, respectively. To calibrate perfusion measurements into units of *ml/100g/min*, two PD-weighted M0 calibration images (TR > 8s) are acquired at the end of the pCASL scan. For susceptibility distortion correction, two phase-encoding-reversed spin-echo images are also acquired.

The ASL data preprocessing is conducted following the HCP ASL Pipeline available at https://github.com/physimals/hcp-asl/ [123]. To run this pipeline we use QuNex platform (singularity container, version 0.99.1) [127]. Briefly, raw ASL and calibration images are first registered to participant anatomical space. This registration transform is merged with motion correction, susceptibility distortion, and gradient distortion transforms and applied simultaneously to the raw ASL and calibration images using the `regtricks` library. This approach minimizes repeated interpolations and thus reduces partial volume effects. Next, intensity bias and banding corrections are applied to the structurally-aligned ASL data. In the presence of head motion, voxels traveling between neighboring slices during the acquisition receive differing intensity scaling during the banding corrections, which can lead to spurious signal after label-control subtraction. To address this effect, general linear model (GLM)-based approach for motion-aware subtraction of banded and background-suppressed ASL data, previously introduced by Suzuki et al. [128], is used. The subtracted time-series data is then used for perfusion estimation. Cerebral blood perfusion and arterial transit time (ATT) are estimated using a variational Bayesian method via the `oxford_asl` script [129]. The `aslrest` Buxton model with cerebral blood perfusion, ATT and macrovascular components is used [130, 131]. A normal distribution prior with a mean of 1 300 *ms* is imposed on ATT and an automatic relevancy determination prior is used on macrovascular perfusion to remove this component from non-arterial voxels. Slice-timing correction is performed by adjusting post-labeling delays in each voxel according to its slice timing offset, and perfusion is converted from arbitrary units into *ml/100g/min* using the mean signal value of cerebral blood flow in the lateral ventricles from the calibration image [132]. Lastly, partial volume effect (PVE) is corrected using a spatial variational Bayes method implemented in `oxford_asl` [131]; the required partial volume estimates are obtained using Toblerone operating on the FreeSurfer-derived cortical surfaces and subcortical segmentations [133, 134]. For PVE correction, a normal distribution prior with a mean of 1 300 *ms* for grey matter ATT and a mean of 1 600 *ms* for white matter ATT is used.

Volumetric cerebral blood perfusion and ATT maps from `oxford_asl` are produced in both ASL-gridded T1-weighted space and MNI152–2 mm template space, via the FNIRT-based registration produced by the HCP structural processing pipeline. To generate output maps in greyordinates space (CIFTI format), the volumetric outputs of `oxford_asl` are projected onto the individual’s native cortical surface using the HCP ribbon-constrained method, registered with MSMAll multimodal areal-feature-based registration and resampled to a common surface mesh, and finally smoothed with a 2 *mm* full-width half-maximum kernel using a surface-constrained method [135–138]. The surface and MNI outputs (masked to consider subcortical structures only) are combined to produce the final CIFTI greyordinates files, which serve as the input data format for subsequent analyses in this study. Participant-specific perfusion maps are originally produced as greyordinates files (including 91 282 vertices/voxels), which we then parcellate them by averaging all greyordinates belonging to the same region of our cortical atlas. Subsequently, we *z*-score the data of individual participants and then *z*-score each region across individuals. Based on normalised down-sampled data, we compute the regions-by-regions covariance matrix across participants.

#### Gene co-expression from microarray transcriptomics

Correlated gene expression represents the transcriptional similarity between pairs of cortical regions [24, 51]. Regional microarray expression data were obtained from 6 post-mortem brains (1 female, ages 24.0– 57.0, 42.50 ±13.38) provided by the Allen Human Brain Atlas (AHBA, https://human.brain-map.org [52]). We followed the same preprocessing as recently described [139]. Briefly, the Allen Human Brain Atlas (AHBA) is a publicly available transcriptional atlas containing gene expression data measured with DNA microarrays and sampled from hundreds of histologically validated neuroanatomical structures across six (five male and one female) normal postmortem human brains. We extracted and mapped gene expression data to the 68 cortical ROIs of the Desikan-Killiany atlas using the `abagen` toolbox (version 0.1.1; https://github.com/rmarkello/ abagen [140]). Data was pooled between homologous cortical regions to ensure adequate coverage of both left (data from six donors) and right hemisphere (data from two donors). Distances between samples were evaluated on the cortical surface with a 2mm distance threshold. Only probes where expression measures were above a background threshold in more than 50% of samples were selected. A representative probe for a gene was selected based on highest intensity. 15,633 genes survived these preprocessing and quality assurance steps. Finally, the region × region correlated gene expression matrix was constructed by correlating (Pearson’s *r*) the normalised gene expression profile at every pair of brain regions. We also replicate our main results using human gene expression data from an alternative modality, RNA-seq, which was available from 2/6 AHBA donors [52].

#### Receptor co-expression from Positron Emission Tomography

Receptor similarity indexes the degree to which the receptor density profiles at two cortical regions are correlated [24, 38]. Here we followed the same procedures as in [24]. Briefly, receptor densities were estimated using PET tracer studies for a total of 18 receptors and transporters, across 9 neurotransmitter systems, recently made available by Hansen and colleagues at https://github.com/netneurolab/hansen_receptors [38]. These include dopamine (D_1_ [141], D_2_ [142–145], DAT [146], noradrenaline (NAT [147–150], serotonin (5-HT_1A_ [151], 5-HT_1_B [151–156], 5-HT_2A_ [157], 5-HT_4_ [157], 5-HT_6_ [158, 159] 5-HTT [157]), acetylcholine (*α*_4_*β*_2_ [160, 161], M_1_ [162], VAChT [163, 164], glutamate (mGluR_5_ [165, 166] NMDA [167, 168], GABA (GABA_A_ [169])), histamine (H_3_ [170]), cannabinoid (CB_1_ [171–174]) and opioid (MOR [175]). Volumetric PET images were registered to the MNI-ICBM 152 nonlinear 2009 (version c, asymmetric) template, averaged across participants within each study, then parcellated and receptors/transporters with more than one mean image of the same tracer (5-HT_1_B, D_2_, VAChT) were combined using a weighted average [38]. A region-by-region receptor similarity matrix was constructed by correlating (Pearson’s *r*) receptor profiles at every pair of cortical regions.

#### Laminar similarity from BigBrain histology

Laminar similarity is estimated from histological data and aims to uncover how similar pairs of cortical regions are in terms of cellular distributions across the cortical laminae. Specifically, we use data from the Big-Brain, a high-resolution (20 *µ*m) histological reconstruction of a post-mortem brain from a 65 year old male [62, 64]. Cell-staining intensity profiles were sampled across 50 equivolumetric surfaces from the pial surface to the white matter surface to estimate laminar variation in neuronal density and soma size. Intensity profiles at various cortical depths can be used to approximately identify boundaries of cortical layers that separate supragranular (cortical layers I–III) granular (cortical layer IV), and infragranular (cortical layers V-VI) layers.

The data were obtained on *fsaverage* surface (164k vertices) from the `BigBrainWarp` toolbox [176] and were parcellated into 68 cortical regions according to the Desikan-Killianty atlas [75]. The region × region laminar similarity matrix was calculated as the partial correlation (Pearson’s *r*) of cell intensities between pairs of cortical regions, after correcting for the mean intensity across cortical regions. Laminar similarity was first introduced in Paquola et al. [64] and has also been referred to as “microstructure profile covariance”.

#### Meta-analytic cognitive similarity from NeuroSynth

Continuous measures of the association between voxels and cognitive categories were obtained from NeuroSynth, an automated term-based meta-analytic tool that synthesizes results from more than 14 000 published fMRI studies by searching for high-frequency key words (such as “pain” and “attention” terms) that are systematically mentioned in the papers alongside fMRI voxel co-ordinates (https://github.com/neurosynth/neurosynth), using the volumetric association test maps [65]. This coninuous measure of association strength is the tendency that a given term is reported in the functional neuroimaging study if there is activation observed at a given voxel. Note that NeuroSynth does not distinguish between areas that are activated or deactivated in relation to the term of interest, nor the degree of activation, only that certain brain areas are frequently reported in conjunction with certain words. Although more than a thousand terms are catalogued in the NeuroSynth engine, we refine our analysis by focusing on cognitive function and therefore we limit the terms of interest to cognitive and behavioural terms. To avoid introducing a selection bias, we opted for selecting terms in a data-driven fashion instead of selecting terms manually. Therefore, terms were selected from the Cognitive Atlas, a public ontology of cognitive science [177], which includes a comprehensive list of neurocognitive terms. This approach totaled to *t* = 123 terms, ranging from umbrella terms (“attention”, “emotion”) to specific cognitive processes (“visual attention”, “episodic memory”), behaviours (“eating”, “sleep”), and emotional states (“fear”, “anxiety”) (note that the 123 term-based meta-analytic maps from NeuroSynth do not explicitly exclude patient studies). The Cognitive Atlas subdivision has previously been used in conjunction with NeuroSynth [178–180], so we opted for the same approach to make our results comparable to previous reports. The probabilistic measure reported by NeuroSynth can be interpreted as a quantitative representation of how regional fluctuations in activity are related to psychological processes. NeuroSynth maps are provided as voxelwise volumetric maps in MNI standard space; as with the resting-state BOLD data, they were parcellated into 100 cortical regions according to the Schaefer atlas [95] (or 68 for the replication with Desikan-Killiany atlas), by averaging the values of all voxels belonging to the same parcel. Finally, we obtain a regions-by-regions matrix of meta-analytic similarity by correlating each pair of regions’ meta-analytic profiles, across terms. The full list of terms included in the present analysis is shown in Table 1.

#### Alternative meta-analytic similarity from BrainMap

Whereas NeuroSynth is an automated tool, Brain-Map is an expert-curated repository: it includes the brain coordinates that are significantly activated during thousands of different experiments from published neuroimaging studies [76, 181]. As a result, NeuroSynth terms and BrainMap behavioural domains differ considerably. Here, we used maps in the Desikan-Killiany anatomical atlas, pertaining to 66 unique behavioural domains (the same as in [179]), obtained from 8, 703 experiments. The full list of BrainMap terms included in the present analysis is shown in Table 2. Experiments conducted on unhealthy participants were excluded, as well as experiments without a defined behavioural domain.

**TABLE 2.**
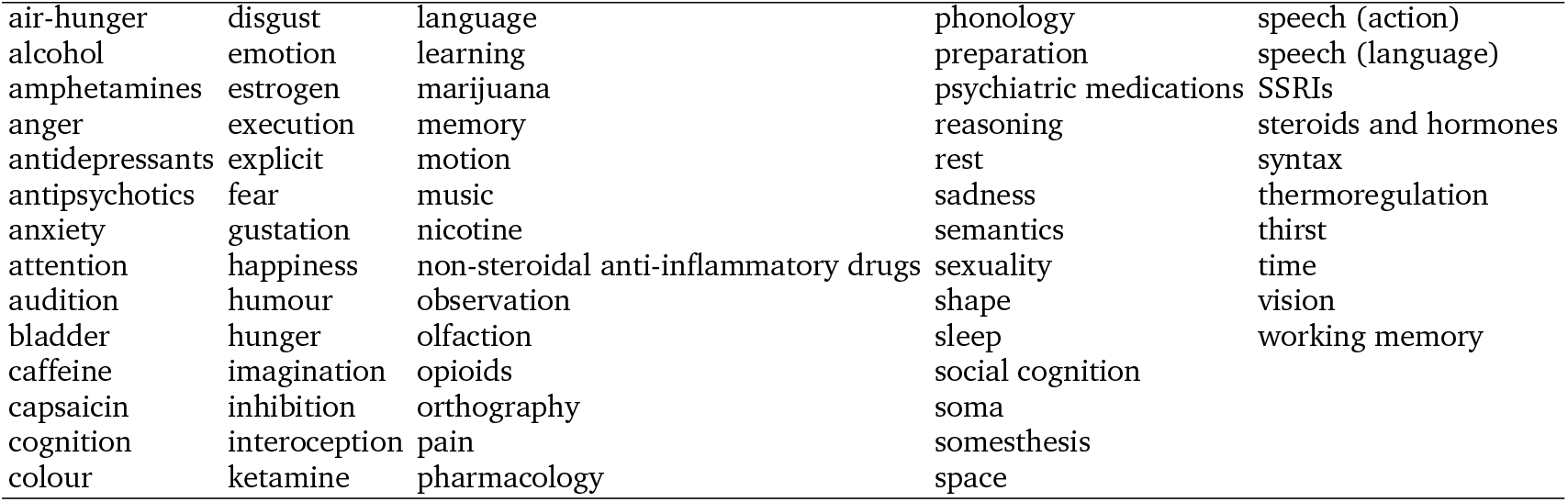
BrainMap terms. BrainMap terms are organized by behavioural domain. All 66 unique behavioural domains (excluding any undefined domains) used in analyses are shown here.

#### Similarity of subjective response to intracranial electrical stimulation

We obtained a matrix of inter-regional similarity of subjective experiences elicited by direct intracranial electrical stimulation (iES), by analysing a previously published dataset of surgical patients (*N* = 1, 537 cortical sites in 67 participants) [73]. Briefly, participants undergoing pre-surgical mapping self-reported their subjective experience upon direct electrical stimulation at different cortical locations. Integration of the first-person reports across the sample yields probabilistic maps of how direct stimulation elicits 8 broad categories of cognitive processes, including ‘emotion’, ‘language’, ‘memory’, and ‘vision’. Based on the intrinsic connectivity network that it belongs to, each cortical region has a ‘fingerprint’ given by its probability of eliciting each of the 8 classes of subjective experiences from [73] upon electrical stimulation. Correlation of these probability fingerprints produces a matrix of similarity of subjective elicitation between each pair of cortical regions.

#### Structural connectivity

We used diffusion MRI (dMRI) data from the same 100 unrelated participants (54 females and 46 males, mean age = 29.1 ± 3.7 years) of the HCP 900 participants data release [182], as described above. The diffusion weighted imaging (DWI) acquisition protocol is covered in detail elsewhere [183]. The diffusion MRI scan was conducted on a Siemens 3T Skyra scanner using a 2D spin-echo single-shot multiband EPI sequence with a multi-band factor of 3 and monopolar gradient pulse. The spatial resolution was 1.25 mm isotropic. TR=5500 ms, TE=89.50ms. The b-values were 1000, 2000, and 3000 s/mm^2^. The total number of diffusion sampling directions was 90, 90, and 90 for each of the shells in addition to 6 b0 images. We used the version of the data made available in DSI Studio-compatible format at http://brain.labsolver.org/ diffusion-mri-templates/hcp-842-hcp-1021 [184].

We adopted previously reported procedures to reconstruct the human connectome from DWI data. The minimally-preprocessed DWI HCP data [183] were corrected for eddy current and susceptibility artifact. DWI data were then reconstructed using q-space diffeomorphic reconstruction (QSDR [185]), as implemented in DSI Studio (www.dsi-studio.labsolver.org). QSDR is a model-free method that calculates the orientational distribution of the density of diffusing water in a standard space, to conserve the diffusible spins and preserve the continuity of fiber geometry for fiber tracking. QSDR first reconstructs diffusion-weighted images in native space and computes the quantitative anisotropy (QA) in each voxel. These QA values are used to warp the brain to a template QA volume in Montreal Neurological Institute (MNI) space using a nonlinear registration algorithm implemented in the statistical parametric mapping (SPM) software. A diffusion sampling length ratio of 2.5 was used, and the output resolution was 1 mm. A modified FACT algorithm [186] was then used to perform deterministic fiber tracking on the reconstructed data, with the following parameters [187]: angular cutoff of 55◦, step size of 1.0 mm, minimum length of 10 mm, maximum length of 400 mm, spin density function smoothing of 0.0, and a QA threshold determined by DWI signal in the cerebrospinal fluid. Each of the streamlines generated was automatically screened for its termination location. A white matter mask was created by applying DSI Studio’s default anisotropy threshold (0.6 Otsu’s threshold) to the spin distribution function’s anisotropy values. The mask was used to eliminate streamlines with premature termination in the white matter region. Deterministic fiber tracking was performed until 1, 000, 000 streamlines were reconstructed for each individual.

For each individual, their structural connectome was reconstructed by drawing an edge between each pair of regions *i* and *j* from the Desikan-Killiany cortical atlas [110] if there were white matter tracts connecting the corresponding brain regions end-to-end; edge weights were quantified as the number of streamlines connecting each pair of regions, normalised by ROI distance and size.

A group-consensus matrix *A* across participants was then obtained using the distance-dependent procedure of Betzel and colleagues, to mitigate concerns about inconsistencies in reconstruction of individual participants’ structural connectomes [188]. This approach seeks to preserve both the edge density and the prevalence and length distribution of inter- and intra-hemispheric edge length distribution of individual participants’ connectomes, and it is designed to produce a representative connectome [188, 189]. This procedure produces a binary consensus network indicating which edges to preserve. The weight of each non-zero edge is then computed as the mean of the corresponding non-zero edges across participants.

#### F-TRACT database of cortico-cortical evoked potentials

As an alternative to diffusion tractography, which is non-invasive but also not causal, we employ a recently assembled database that reports the amplitude of cortico-cortical evoked potentials between pairs of cortical regions, from the large multicentric F-TRACT consortium [77, 190, 191]. Briefly, cortico-cortical evoked potentials were quantified by stereo-EEG (SEEG) As part of a presurgical evaluation of patients’ drug-resistant epilepsy. The CCEPs were recorded using local clinical practice, following 1-Hz stimulations (99.5% of CCEPs), or rarely 2 or 3 Hz (0.5% of CCEPs), between two contiguous contacts located either in the grey matter (61% of contacts) or white matter (39% of contacts), using either monophasic (20% of CCEPs) or biphasic (80% of CCEPs) electrical pulses. The number of pulses in a row were up to 40 and only stimulation runs with at least 3 pulses were considered. On average, the patients were implanted with 320 contacts (min: 60, max: 618) and were stimulated 58 times (min: 1, max: 257). The evoked response was considered significant if its absolute value reached the threshold of z = 5 during the first 200 ms post-stimulation onset. This condition indicated the presence of a functional link between the stimulation and the recording sites [77, 190, 191]. Fully processed data at the group level are available for download on the atlas page F-TRACT website (f-tract.eu/atlas). In total, the database comprises data from *>* 34 354 direct electrical stimulations in 780 patients [77].

### Dominance analysis

To consider all networks of inter-regional interactions together and evaluate their respective contributions, we performed a dominance analysis with the vectorised brain networks of Euclidean distance, structural connectivity, gene co-expression, receptor similarity, laminar similarity, electrophysiological similarity, metabolic similarity, and meta-analytic cognitive co-activation as predictors; and each participant’s vectorised network of haemodynamic connectivity from fMRI as target. Dominance analysis seeks to determine the relative contribution (“dominance” of each independent variable to the overall fit (adjusted *R*^2^)) of the multiple linear regression model (https://github.com/dominance-analysis/dominance-analysis) [69]. This is done by fitting the same regression model on every combination of predictors (2^*p*^ − 1 submodels for a model with *p* predictors). Total dominance is defined as the average of the relative increase in *R*^2^ when adding a single predictor of interest to a submodel, across all 2^*p*^ − 1 submodels. The sum of the dominance of all input variables is equal to the total adjusted *R*^2^ of the complete model, making the percentage of relative importance an intuitive method that partitions the total effect size across predictors.

Subject-wise relative-importance values for each predictor were then entered into a linear mixed-effects modelling framework to test whether the contribution of each multimodal network to haemodynamic functional connectivity changed across consciousness states. For each predictor network, the dependent variable was the relative importance obtained from dominance analysis. We modelled these values as a function of experimental condition while controlling for head motion (mean framewise displacement) as a covariate of no interest.

### Supportvector machine

To assess whether subject-wise patterns of multimodal network contributions could distinguish normal from perturbed consciousness, we trained a linear support vector machine (SVM) implemented as MATLAB’s `fitcsvm` function with linear kernel, using each participant’s dominance-analysis profile as the feature vector. Each feature corresponded to the relative importance of one structural or functional predictor network for explaining that participant’s haemodynamic FC. Training data included healthy controls and DOC patients, as well as pre-anaesthetic baseline and anaesthesia scans from the pharmacological datasets; post-anaesthetic recovery scans were excluded from training and reserved for out-of-sample evaluation. Observations were labelled as baseline or perturbed consciousness, and classification performance was estimated using leave-one-subject-out cross-validation. Statistical significance was assessed by repeating the same cross-validation procedure after randomly permuting class labels (5,000 permutations). Finally, the classifier trained on all training data was applied to unseen recovery scans and independent DOC subgroups to evaluate model generalisation. McNemar’s test was used to compare the classification accuracies obtained from the SVM trained with dominance profiles as features, and the SVM trained with univariate correlations as features.

## Data and code availability

The original pharmacological and DOC fMRI data are available from the corresponding authors of the original publications referenced herein. Human networks of metabolic connectivity, electrophysiological connectivity, and laminar similarity are available on GitHub at https://github.com/netneurolab/hansen_many_networks/tree/v1.0.0. Diffusion MRI data for the Human Connectome Project in DSI Studio-compatible format are available at http://brain.labsolver.org/diffusion-mri-templates/hcp-842-hcp-1021. NeuroSynth meta-analytic maps are available at https://neurosynth.org/. Maps of subjective experience probabilities elicited by intracranial electrical stimulation are available from the supplementary materials of [73]. The F-TRACT database of cortico-cortical evoked potentials is available at f-tract.eu/atlas. The Human PET receptor and transporter maps are available at https://github.com/netneurolab/hansen_receptors. Human gene expression data from the Allen Human Brain Atlas are available at https://human.brain-map.org/. The `abagen` toolbox for processing of the AHBA human transcriptomic dataset is available at https://abagen.readthedocs.io/. Third-party Python software (version 3.8) for dominance analysis is freely available at https://github.com/dominance-analysis/dominance-analysis.

## Acknowledgments

A.I.L. acknowledges the support of the Natural Sciences and Engineering Research Council of Canada (NSERC), [funding reference number 202209BPF-489453-401636, Banting Postdoctoral Fellowship] and FRQNT Strategic Clusters Program (2020-RS4-265502 - Centre UNIQUE - Union Neuroscience & Artificial Intelligence - Quebec) via the UNIQUE Neuro-AI Excellence Award; CIHR (Funding reference ICT-201439); St John’s College, Cambridge; and a Wellcome Early Career Award (grant number 226924/Z/23/Z). B.M. acknowledges support from the Natural Sciences and Engineering Research Council of Canada (NSERC), Canadian Institutes of Health Research (CIHR), Brain Canada Foundation Future Leaders Fund, the Canada Research Chairs Program, the Michael J. Fox Foundation, and the Healthy Brains for Healthy Lives initiative. D.M. is supported by the MIT-Novo Nordisk Artificial Intelligence Postdoctoral Fellows Program. N.L.N.A. acknowledges support of the Belgian National Funds for Scientific Research (FRS-FNRS). A.M.O. acknowledges support from the Canadian Institutes of Health Research (CIHR) and is a Fellow of CIFAR Mind Brain and Consciousness program. This study was also supported by the University of Liège, the University Hospital of Liège and its Algology Interdisciplinary Center. For the purpose of open access, the authors have applied a Creative Commons Attribution (CC BY) licence to any Author Accepted Manuscript version arising from this submission.

## Supplementary Materials

## References

[1] Bharat B Biswal, Maarten Mennes, Xi-Nian Zuo, Suril Gohel, Clare Kelly, Steve M Smith, Christian F Beckmann, Jonathan S Adelstein, Randy L Buckner, Stan Colcombe, et al. Toward discovery science of human brain function. Proceedings of the national academy of sciences, 107(10):4734–4739, 2010.

[2] Bharat B Biswal and Lucina Q Uddin. The history and future of resting-state functional magnetic resonance imaging. Nature, 641(8065):1121–1131, 2025.

[3] Michael D Fox and Marcus E Raichle. Spontaneous fluctuations in brain activity observed with functional magnetic resonance imaging. Nature reviews neuroscience, 8 (9):700–711, 2007.

[4] Stephen M Smith, Thomas E Nichols, Diego Vidaurre, Anderson M Winkler, Timothy EJ Behrens, Matthew F Glasser, Kamil Ugurbil, Deanna M Barch, David C Van Essen, and Karla L Miller. A positive-negative mode of population covariation links brain connectivity, demographics and behavior. Nature neuroscience, 18(11): 1565–1567, 2015.

[5] Karla L Miller, Fidel Alfaro-Almagro, Neal K Bangerter, David L Thomas, Essa Yacoub, Junqian Xu, Andreas J Bartsch, Saad Jbabdi, Stamatios N Sotiropoulos, Jes-per LR Andersson, et al. Multimodal population brain imaging in the uk biobank prospective epidemiological study. Nature neuroscience, 19(11):1523–1536, 2016.

[6] Monica D Rosenberg, Emily S Finn, Dustin Scheinost, Xenophon Papademetris, Xilin Shen, R Todd Constable, and Marvin M Chun. A neuromarker of sustained attention from whole-brain functional connectivity. Nature neuroscience, 19(1):165–171, 2016.

[7] Xilin Shen, Emily S Finn, Dustin Scheinost, Monica D Rosenberg, Marvin M Chun, Xenophon Papademetris, and R Todd Constable. Using connectome-based predictive modeling to predict individual behavior from brain connectivity. nature protocols, 12(3):506–518, 2017.

[8] Emily S Finn, Xilin Shen, Dustin Scheinost, Monica D Rosenberg, Jessica Huang, Marvin M Chun, Xenophon Papademetris, and R Todd Constable. Functional connectome fingerprinting: identifying individuals using patterns of brain connectivity. Nature neuroscience, 18 (11):1664–1671, 2015.

[9] Michael D Fox. Mapping symptoms to brain networks with the human connectome. New England Journal of Medicine, 379(23):2237–2245, 2018.

[10] Zhen-Qi Liu, Andrea I Luppi, Justine Y Hansen, Ye Ella Tian, Andrew Zalesky, BT Thomas Yeo, Ben D Fulcher, and Bratislav Misic. Benchmarking methods for mapping functional connectivity in the brain. Nature Methods, 2025.

[11] Michael W Cole, Danielle S Bassett, Jonathan D Power, Todd S Braver, and Steven E Petersen. Intrinsic and task-evoked network architectures of the human brain. Neuron, 83(1):238–251, 2014. ISSN 10974199. doi: 10.1016/j.neuron.2014.05.014. PMID: 24991964 arXiv: NIHMS150003 Citation Key: Cole2014 ISBN: 1097-4199 (Electronic)\r0896-6273 (Linking).

[12] Angela R Laird, Simon B Eickhoff, Claudia Rottschy, Danilo Bzdok, Kimberly L Ray, and Peter T Fox. Networks of task co-activations. Neuroimage, 80:505–514, 2013.

[13] Stephen M Smith, Peter T Fox, Karla L Miller, David C Glahn, Peter Mickle Fox, Clare E Mackay, Nicola Filippini, Kate E Watkins, Roberto Toro, Angela R Laird, and Christian F Beckmann. Correspondence of the brain’s functional architecture during activation and rest. Proceedings of the National Academy of Sciences, 106(31):13040–13045, 2009. ISSN 0027-8424. doi: 10.1073/pnas.0905267106. PMID: 19620724 0905267106 Citation Key: Smith2009b ISBN: 1091-6490 (Electronic)\r1091-6490 (Linking).

[14] Nicolas A. Crossley, Andrea Mechelli, Petra E. Vértes, Toby T. Winton-Brown, Ameera X. Patel, Cedric E. Ginestet, Philip McGuire, and Edward T. Bullmore. Cognitive relevance of the community structure of the human brain functional coactivation network. Proceedings of the National Academy of Sciences, 110(28):11583–11588, 2013. doi:10.1073/pnas.1220826110.

[15] Roberto Toro, Peter T Fox, and Tomáš Paus. Functional coactivation map of the human brain. Cerebral cortex, 18(11):2553–2559, 2008.

[16] Evan Collins, Omar Chishti, Sami Obaid, Hari McGrath, Alex King, Xilin Shen, Jagriti Arora, Xenophon Papademetris, R Todd Constable, Dennis D Spencer, et al. Mapping the structure-function relationship along macroscale gradients in the human brain. Nature Communications, 15(1):7063, 2024.

[17] Pablo Barttfeld, Lynn Uhrig, Jacobo D Sitt, Mariano Sigman, Béchir Jarraya, and Stanislas Dehaene. Signature of consciousness in the dynamics of resting-state brain activity. Proceedings of the National Academy of Sciences, 112(3):887–892, 2015.

[18] Athena Demertzi, Enzo Tagliazucchi, Stanislas Dehaene, Gustavo Deco, Pablo Barttfeld, Federico Raimondo, Charlotte Martial, Davinia Fernández-Espejo, Benjamin Rohaut, HU Voss, et al. Human consciousness is supported by dynamic complex patterns of brain signal coordination. Science advances, 5(2):eaat7603, 2019.

[19] Andrea I. Luppi, Lynn Uhrig, Jordy Tasserie, Camilo M. Signorelli, Emmanuel A. Stamatakis, Alain Destexhe, Bechir Jarraya, and Rodrigo Cofre. Local orchestration of distributed functional patterns supporting loss and restoration of consciousness in the primate brain. Nature Communications, 15(1):2171, mar 11 2024. ISSN 2041-1723. doi:10.1038/s41467-024-46382-w. URL https://www.nature.com/articles/s41467-024-46382-w. publisher: Nature Publishing Group.

[20] Jordy Tasserie, Lynn Uhrig, Jacobo D Sitt, Dragana Manasova, Morgan Dupont, Stanislas Dehaene, and Béchir Jarraya. Deep brain stimulation of the thalamus restores signatures of consciousness in a nonhuman primate model. Sci. Adv, 8:5547, 2022. doi: 10.1126/SCIADV.ABL5547. URL https://www.science.org. [Online; accessed 2022-03-20].

[21] Daniel Gutierrez-Barragan, Neha Atulkumar Singh, Filomena Grazia Alvino, Ludovico Coletta, Federico Rocchi, Elizabeth De Guzman, Alberto Galbusera, Mauro Uboldi, Stefano Panzeri, and Alessandro Gozzi. Unique spatiotemporal fmri dynamics in the awake mouse brain. Current biology, 32(2):1–14, 1 2021. ISSN 1879-0445. doi:10.1016/J.CUB.2021.12.015. PMID: 34998465 publisher: Curr Biol.

[22] Andrea I Luppi, Fernando E Rosas, Pedro AM Mediano, Athena Demertzi, David K Menon, and Emmanuel A Stamatakis. Unravelling consciousness and brain function through the lens of time, space, and information. Trends in Neurosciences, 2024.

[23] Andrea I Luppi, Daniel Golkowski, Andreas Ranft, Rudi-ger Ilg, Denis Jordan, Danilo Bzdok, Adrian M Owen, Lorina Naci, Emmanuel A Stamatakis, Enrico Amico, et al. General anaesthesia decreases the uniqueness of brain functional connectivity across individuals and species. Nature Human Behaviour, pages 1–18, 2025.

[24] Justine Y Hansen, Golia Shafiei, Katharina Voigt, Emma X Liang, Sylvia ML Cox, Marco Leyton, Sharna D Jamadar, and Bratislav Misic. Integrating multimodal and multiscale connectivity blueprints of the human cerebral cortex in health and disease. PLoS biology, 21 (9):e3002314, 2023.

[25] Vincent Bazinet, Justine Y Hansen, and Bratislav Misic. Towards a biologically annotated brain connectome. Nature reviews neuroscience, 24(12):747–760, 2023.

[26] Laura E. Suárez, Ross D. Markello, Richard F. Betzel, and Bratislav Misic. Linking structure and function in macroscale brain networks. Trends in Cognitive Sciences, 24(4):302–315, 4 2020. ISSN 1879307X. doi:10.1016/j.tics.2020.01.008. PMID: 32160567 publisher: Elsevier Ltd.

[27] Panagiotis Fotiadis, Linden Parkes, Kathryn A Davis, Theodore D Satterthwaite, Russell T Shinohara, and Dani S Bassett. Structure–function coupling in macroscale human brain networks. Nature Reviews Neuroscience, 25(10):688–704, 2024.

[28] Andrea Avena-Koenigsberger, Bratislav Misic, and Olaf Sporns. Communication dynamics in complex brain networks. Nature Reviews Neuroscience, 19(1):17–33, 2018.

[29] Caio Seguin, Olaf Sporns, and Andrew Zalesky. Brain network communication: concepts, models and applications. Nature Reviews Neuroscience, pages 1–18, jul 12 2023. ISSN 1471-0048. doi:10.1038/s41583-023-00718-5. URL https://www.nature.com/articles/s41583-023-00718-5. publisher: Nature Publishing Group.

[30] Bertha Vázquez-Rodríguez, Laura E Suárez, Ross D Markello, Golia Shafiei, Casey Paquola, Patric Hagmann, Martijn P Van Den Heuvel, Boris C Bernhardt, R Nathan Spreng, and Bratislav Misic. Gradients of structure– function tethering across neocortex. Proceedings of the National Academy of Sciences, 116(42):21219–21227, 2019.

[31] Graham L Baum, Zaixu Cui, David R Roalf, Rastko Ciric, Richard F Betzel, Bart Larsen, Matthew Cieslak, Philip A Cook, Cedric H Xia, Tyler M Moore, et al. Development of structure–function coupling in human brain networks during youth. Proceedings of the National Academy of Sciences, 117(1):771–778, 2020.

[32] Maria Giulia Preti and Dimitri Van De Ville. Decoupling of brain function from structure reveals regional behavioral specialization in humans. Nature Communications, 10(1), 2019. ISSN 20411723. doi: 10.1038/s41467-019-12765-7. URL https://doi.org/10.1038/s41467-019-12765-7. 1905.07813.

[33] Selen Atasoy, Isaac Donnelly, and Joel Pearson. Human brain networks function in connectome-specific harmonic waves. Nature communications, 7(1):10340, 2016.

[34] Gustavo Deco, Adrián Ponce-Alvarez, Dante Mantini, Gian Luca Romani, Patric Hagmann, and Maurizio Corbetta. Resting-state functional connectivity emerges from structurally and dynamically shaped slow linear fluctuations. Journal of Neuroscience, 33(27):11239–11252, 2013. doi:10.1523/JNEUROSCI.1091-13.2013.

[35] Christopher J. Honey, Rolf Kötter, Michael Breakspear, and Olaf Sporns. Network structure of cerebral cortex shapes functional connectivity on multiple time scales. Proceedings of the National Academy of Sciences of the United States of America, 104(24):10240–10245, 6 2007. ISSN 00278424. doi:10.1073/pnas.0701519104. PMID: 17548818 publisher: National Academy of Sciences.

[36] C. J. Honey, O. Sporns, L. Cammoun, X. Gigandet, J. P. Thiran, R. Meuli, and P. Hagmann. Predicting human resting-state functional connectivity from structural connectivity. Proceedings of the National Academy of Sciences, 106(6):2035–2040, 2 2009. doi: 10.1073/pnas.0811168106. publisher: Proceedings of the National Academy of Sciences.

[37] Alexandros Goulas, Jean-Pierre Changeux, Konrad Wagstyl, Katrin Amunts, Nicola Palomero-Gallagher, and Claus C Hilgetag. The natural axis of transmitter receptor distribution in the human cerebral cortex. Proceedings of the National Academy of Sciences, 118(3): e2020574118, 2021.

[38] Justine Y Hansen, Golia Shafiei, Ross D Markello, Sylvia Cox, Kelly Smart, Etienne Aumont, Stijn Servaes, Stephanie Scala, Gabriel Wainstein, Gleb Bez-gin, Thomas Funck, W Schmitz, Marc-andre Bedard, R Nathan Spreng, Jean-paul Soucy, and Synthia Guimond. Mapping neurotransmitter systems to the structural and functional organization of the human neocortex. Nature Neuroscience, (accepted):1–26, 2022.

[39] Karl Zilles and Nicola Palomero-Gallagher. Multiple transmitter receptors in regions and layers of the human cerebral cortex. Frontiers in neuroanatomy, 11:78, 2017.

[40] Sean Froudist-Walsh, Ting Xu, Meiqi Niu, Lucija Rapan, Ling Zhao, Daniel S Margulies, Karl Zilles, Xiao-Jing Wang, and Nicola Palomero-Gallagher. Gradients of neurotransmitter receptor expression in the macaque cortex. Nature neuroscience, 26(7):1281–1294, 2023.

[41] Sharna D Jamadar, Phillip GD Ward, Emma X Liang, Edwina R Orchard, Zhaolin Chen, and Gary F Egan. Metabolic and hemodynamic resting-state connectivity of the human brain: a high-temporal resolution simultaneous bold-fmri and fdg-fpet multimodality study. Cerebral Cortex, 31(6):2855–2867, 2021.

[42] Katharina Voigt, Emma X Liang, Bratislav Misic, Phillip GD Ward, Gary F Egan, and Sharna D Jamadar. Metabolic and functional connectivity provide unique and complementary insights into cognition-connectome relationships. Cerebral Cortex, 33(4):1476–1488, 2023.

[43] Tommaso Volpi, Erica Silvestri, Marco Aiello, John J Lee, Andrei G Vlassenko, Manu S Goyal, Maurizio Corbetta, and Alessandra Bertoldo. The brain’s “dark energy” puzzle: How strongly is glucose metabolism linked to resting-state brain activity? Journal of Cerebral Blood Flow & Metabolism, 44(8):1433–1449, 2024.

[44] Golia Shafiei, Ben D Fulcher, Bradley Voytek, Theodore D Satterthwaite, Sylvain Baillet, and Bratislav Misic. Neurophysiological signatures of cortical microarchitecture. Nature communications, 14(1):6000, 2023.

[45] Golia Shafiei, Sylvain Baillet, and Bratislav Misic. Human electromagnetic and haemodynamic networks systematically converge in unimodal cortex and diverge in transmodal cortex. PLoS biology, 20(8):e3001735, 2022.

[46] Catherine Jean Chu, Naoaki Tanaka, J Diaz, Brian L Edlow, Ona Wu, M Hämäläinen, S Stufflebeam, Sydney S Cash, and Mark A Kramer. Eeg functional connectivity is partially predicted by underlying white matter connectivity. Neuroimage, 108:23–33, 2015.

[47] Matthew J Brookes, Joanne R Hale, Johanna M Zumer, Claire M Stevenson, Susan T Francis, Gareth R Barnes, Julia P Owen, Peter G Morris, and Srikantan S Nagarajan. Measuring functional connectivity using meg: methodology and comparison with fcmri. Neuroimage, 56(3):1082–1104, 2011.

[48] Biyu J He, Abraham Z Snyder, John M Zempel, Matthew D Smyth, and Marcus E Raichle. Electrophysiological correlates of the brain’s intrinsic large-scale functional architecture. Proceedings of the National Academy of Sciences, 105(41):16039–16044, 2008.

[49] Matthew J Brookes, Mark Woolrich, Henry Luckhoo, Darren Price, Joanne R Hale, Mary C Stephenson, Gareth R Barnes, Stephen M Smith, and Peter G Morris. Investigating the electrophysiological basis of resting state networks using magnetoencephalography. Proceedings of the National Academy of Sciences, 108(40): 16783–16788, 2011.

[50] James C Pang, Kevin M Aquino, Marianne Oldehinkel, Peter A Robinson, Ben D Fulcher, Michael Breakspear, and Alex Fornito. Geometric constraints on human brain function. Nature, 618(7965):566–574, 2023.

[51] Jonas Richiardi, Andre Altmann, Anna-Clare Milazzo, Catie Chang, M Mallar Chakravarty, Tobias Banaschewski, Gareth J Barker, Arun LW Bokde, Uli Bromberg, Christian Büchel, et al. Correlated gene expression supports synchronous activity in brain networks. Science, 348(6240):1241–1244, 2015.

[52] Michael J Hawrylycz, Ed S Lein, Angela L Guillozet-Bongaarts, Elaine H Shen, Lydia Ng, Jeremy A Miller, Louie N Van De Lagemaat, Kimberly A Smith, Amanda Ebbert, Zackery L Riley, et al. An anatomically comprehensive atlas of the adult human brain transcriptome. Nature, 489(7416):391, 2012.

[53] Ed S Lein, Michael J Hawrylycz, Nancy Ao, Mikael Ayres, Amy Bensinger, Amy Bernard, Andrew F Boe, Mark S Boguski, Kevin S Brockway, Emi J Byrnes, et al. Genomewide atlas of gene expression in the adult mouse brain. Nature, 445(7124):168–176, 2007.

[54] Tingting Bo, Jie Li, Ganlu Hu, Ge Zhang, Wei Wang, Qian Lv, Shaoling Zhao, Junjie Ma, Meng Qin, Xiaohui Yao, et al. Brain-wide and cell-specific transcriptomic insights into mri-derived cortical morphology in macaque monkeys. Nature Communications, 14(1):1499, 2023.

[55] Ao Chen, Yidi Sun, Ying Lei, Chao Li, Sha Liao, Juan Meng, Yiqin Bai, Zhen Liu, Zhifeng Liang, Zhiyong Zhu, et al. Single-cell spatial transcriptome reveals cell-type organization in the macaque cortex. Cell, 186(17): 3726–3743, 2023.

[56] Casey Paquola, Jakob Seidlitz, Oualid Benkarim, Jessica Royer, Petr Klimes, Richard AI Bethlehem, Sara Larivière, Reinder Vos de Wael, Raul Rodríguez-Cruces, Jeffery A Hall, et al. A multi-scale cortical wiring space links cellular architecture and functional dynamics in the human brain. PLoS biology, 18(11):e3000979, 2020.

[57] Matthew F Glasser and David C Van Essen. Mapping human cortical areas in vivo based on myelin content as revealed by t1-and t2-weighted mri. Journal of neuroscience, 31(32):11597–11616, 2011.

[58] Joshua B. Burt, Murat Demirtaş, William J. Eckner, Natasha M. Navejar, Jie Lisa Ji, William J. Martin, Alberto Bernacchia, Alan Anticevic, and John D. Murray. Hierarchy of transcriptomic specialization across human cortex captured by structural neuroimaging topography. Nature Neuroscience, 21(9):1251–1259, 9 2018. ISSN 15461726. doi:10.1038/s41593-018-0195-0. PMID: 30082915 publisher: Nature Publishing Group.

[59] Julia M Huntenburg, Pierre-Louis Bazin, Alexandros Goulas, Christine L Tardif, Arno Villringer, and Daniel S Margulies. A systematic relationship between functional connectivity and intracortical myelin in the human cerebral cortex. Cerebral Cortex, 27(2):981–997, 2017.

[60] Ben D Fulcher, John D Murray, Valerio Zerbi, and Xiao-Jing Wang. Multimodal gradients across mouse cortex. Proc Natl Acad Sci USA, 116(10):4689–4695, 2019.

[61] Isaac Sebenius, Jakob Seidlitz, Varun Warrier, Richard AI Bethlehem, Aaron Alexander-Bloch, Travis T Mallard, Rafael Romero Garcia, Edward T Bullmore, and Sarah E Morgan. Robust estimation of cortical similarity networks from brain mri. Nature neuroscience, 26(8):1461–1471, 2023.

[62] Katrin Amunts, Claude Lepage, Louis Borgeat, Hartmut Mohlberg, Timo Dickscheid, Marc-Étienne Rousseau, Sebastian Bludau, Pierre-Louis Bazin, Lindsay B Lewis, Ana-Maria Oros-Peusquens, et al. Bigbrain: an ultrahigh-resolution 3d human brain model. science, 340 (6139):1472–1475, 2013.

[63] Dragana Manasova, Laouen Mayal Louan Bel-loli, Martin Justinus Rosenfelder, Lina Willacker, Emilia Fló Rama Chiara Valota, Bertrand Hermann, Brigitte Charlotte Kaufmann, Alice Pirastru, Chiara Camilla Derchi, et al. Multimodal multicentre investigation of diagnostic and prognostic markers in disorders of consciousness. Brain, page awaf412, 2026.

[64] Casey Paquola, Reinder Vos De Wael, Konrad Wagstyl, Richard AI Bethlehem, Seok-Jun Hong, Jakob Seidlitz, Edward T Bullmore, Alan C Evans, Bratislav Misic, Daniel S Margulies, et al. Microstructural and functional gradients are increasingly dissociated in transmodal cortices. PLoS biology, 17(5):e3000284, 2019.

[65] Tal Yarkoni, Russell A Poldrack, Thomas E Nichols, David C Van Essen, and Tor D Wager. Large-scale automated synthesis of human functional neuroimaging data. Nature methods, 8(8):665–670, 2011.

[66] Andrea I Luppi, Michael M Craig, Ioannis Pappas, Paola Finoia, Guy B Williams, Judith Allanson, John D. Pickard, Adrian M. Owen, Lorina Naci, David K Menon, and Emmanuel A. Stamatakis. Consciousness-specific dynamic interactions of brain integration and functional diversity. Nature Communications, 10(1), 2019. ISSN 20411723. doi:10.1038/s41467-019-12658-9. PMID: 31601811.

[67] Andreas Ranft, Daniel Golkowski, Tobias Kiel, Valentin Riedl, Philipp Kohl, Guido Rohrer, Joachim Pientka, Sebastian Berger, Alexander Thul, Max Maurer, Christine Preibisch, Claus Zimmer, George A Mashour, Eberhard F Kochs, Denis Jordan, and Rüdiger Ilg. Neural correlates of sevoflurane-induced unconsciousness identified by simultaneous functional magnetic resonance imaging and electroencephalography. Anesthesiology, 125(5):861–872, 2016. ISSN 1528-1175. doi: 10.1097/ALN.0000000000001322. PMID: 27617689 Citation Key: Ranft2016 ISBN: 0000000000.

[68] Vincent Bonhomme, Audrey Vanhaudenhuyse, Athena Demertzi, Marie-Aurélie Bruno, Oceane Jaquet, Mohamed Ali Bahri, Alain Plenevaux, Melanie Boly, Pierre Boveroux, Andrea Soddu, Jean François Brichant, Pierre Maquet, and Steven Laureys. Resting-state network-specific breakdown of functional connectivity during ketamine alteration of consciousness in volunteers. Anesthesiology, 125(5):873–888, 11 2016. ISSN 0003-3022. doi:10.1097/ALN.0000000000001275. publisher: The American Society of Anesthesiologists Citation Key: Bonhomme 2016.

[69] Razia Azen and David V Budescu. The dominance analysis approach for comparing predictors in multiple regression. Psychological methods, 8(2):129, 2003.

[70] Justine Y Hansen, Golia Shafiei, Jacob W Vogel, Kelly Smart, Carrie E Bearden, Martine Hoogman, Barbara Franke, Daan Van Rooij, Jan Buitelaar, Carrie R McDonald, et al. Local molecular and global connectomic contributions to cross-disorder cortical abnormalities. Nature communications, 13(1):4682, 2022.

[71] Milan Van Maldegem, Jakub Vohryzek, Selen Atasoy, Naji Alnagger, Paolo Cardone, Vincent Bonhomme, Audrey Vanhaudenhuyse, Athena Demertzi, Oceane Jaquet, Mohamed Ali Bahri, et al. Connectome harmonic decomposition tracks the presence of disconnected consciousness during ketamine-induced unresponsiveness. British Journal of Anaesthesia, 2025.

[72] Valerie J Sydnor, Bart Larsen, Danielle S Bassett, Aaron Alexander-Bloch, Damien A Fair, Conor Liston, Allyson P Mackey, Michael P Milham, Adam Pines, David R Roalf, Jakob Seidlitz, Ting Xu, Armin Raznahan, and Theodore D Satterthwaite. Neurodevelopment of the association cortices: Patterns, mechanisms, and implications for psychopathology. Neuron, 2021. ISSN 08966273. doi:10.1016/j.neuron.2021.06.016. URL https://doi.org/10.1016/j.neuron.2021.06.016. [On-line; accessed 2021-07-26].

[73] Kieran CR Fox, Lin Shi, Sori Baek, Omri Raccah, Brett L Foster, Srijani Saha, Daniel S Margulies, Aaron Kucyi, and Josef Parvizi. Intrinsic network architecture predicts the effects elicited by intracranial electrical stimulation of the human brain. Nature human behaviour, 4(10): 1039–1052, 2020.

[74] Shan H. Siddiqi, Konrad P. Kording, Josef Parvizi, and Michael D. Fox. Causal mapping of human brain function. Nature Reviews Neuroscience, 23(6):361–375, 2022. doi:10.1038/s41583-022-00583-8.

[75] Rahul S Desikan, Florent Ségonne, Bruce Fischl, Brian T Quinn, Bradford C Dickerson, Deborah Blacker, Randy L Buckner, Anders M Dale, R Paul Maguire, Bradley T Hyman, et al. An automated labeling system for subdividing the human cerebral cortex on mri scans into gyral based regions of interest. NeuroImage, 31(3):968–980, 2006.

[76] Angela R Laird, Jack J Lancaster, and Peter T Fox. Brainmap: the social evolution of a human brain mapping database. Neuroinformatics, 3:65–77, 2005.

[77] Jean-Didier Lemaréchal, Maciej Jedynak, Lena Trebaul, Anthony Boyer, François Tadel, Manik Bhattacharjee, Pierre Deman, Viateur Tuyisenge, Leila Ayoubian, Etienne Hugues, et al. A brain atlas of axonal and synaptic delays based on modelling of cortico-cortical evoked potentials. Brain, 145(5):1653–1667, 2022.

[78] Karl J Friston. Functional and effective connectivity: a review. Brain connectivity, 1(1):13–36, 2011.

[79] Enrique CA Hansen, Demian Battaglia, Andreas Spiegler, Gustavo Deco, and Viktor K Jirsa. Functional connectivity dynamics: Modeling the switching behavior of the resting state. NeuroImage, 105:525–535, 2015. doi: 10.1016/j.neuroimage.2014.11.001.

[80] Sylvain Baillet. Magnetoencephalography for brain electrophysiology and imaging. Nature neuroscience, 20(3): 327–339, 2017.

[81] Jordy Tasserie, Lynn Uhrig, Jacobo D Sitt, Dragana Manasova, Morgan Dupont, Stanislas Dehaene, and Béchir Jarraya. Deep brain stimulation of the thalamus restores signatures of consciousness in a nonhuman primate model. Sci. Adv, 8:5547, 2022. doi: 10.1126/SCIADV.ABL5547.

[82] Lennart RB Spindler, Andrea I Luppi, Ram M Adapa, Michael M Craig, Peter Coppola, Alexander RD Peattie, Anne E Manktelow, Paola Finoia, Barbara J Sahakian, Guy B Williams, et al. Dopaminergic brainstem disconnection is common to pharmacological and pathological consciousness perturbation. Proceedings of the National Academy of Sciences, 118(30):e2026289118, 2021.

[83] James M. Shine, Laura D. Lewis, Douglas D. Garrett, and Kai Hwang. The impact of the human thalamus on brain-wide information processing. Nature Reviews. Neuroscience, 24(7):416–430, 7 2023. ISSN 1471-0048. doi:10.1038/s41583-023-00701-0. PMID: 37237103.

[84] Andrea I. Luppi, Lynn Uhrig, Jordy Tasserie, Camilo M. Signorelli, Emmanuel A. Stamatakis, Alain Destexhe, Bechir Jarraya, and Rodrigo Cofre. Local orchestration of distributed functional patterns supporting loss and restoration of consciousness in the primate brain. Nature Communications, 15(1):2171, mar 11 2024. ISSN 2041-1723. doi:10.1038/s41467-024-46382-w. publisher: Nature Publishing Group.

[85] Brian L Edlow, Mark Olchanyi, Holly J Freeman, Jian Li, Chiara Maffei, Samuel B Snider, Lilla Zöllei, J Eugenio Iglesias, Jean Augustinack, Yelena G Bodien, et al. Multimodal mri reveals brainstem connections that sustain wakefulness in human consciousness. Science Translational Medicine, 16(745):eadj4303, 2024.

[86] Eli J Müller, Brandon R Munn, Michelle J Redinbaugh, Joseph Lizier, Michael Breakspear, Yuri B Saalmann, and James M Shine. The non-specific matrix thalamus facilitates the cortical information processing modes relevant for conscious awareness. Cell reports, 42(8), 2023.

[87] Brandon R. Munn, Eli J. Müller, Jaan Aru, Christopher J. Whyte, Albert Gidon, Matthew E. Larkum, and James M. Shine. A thalamocortical substrate for integrated information via critical synchronous bursting. Proceedings of the National Academy of Sciences, 120(46):e2308670120, nov 14 2023. doi: 10.1073/pnas.2308670120. publisher: Proceedings of the National Academy of Sciences.

[88] Michelle J. Redinbaugh, J.M. Phillips, N.A. Kambi, S. Mohanta, S. Andryk, G.L. Dooley, Mohsen Afrasiabi, Aeyal Raz, and Yuri B. Saalmann. Thalamus modulates consciousness via layer-specific control of cortex. Neuron, 106:66–75.e12, 2020.

[89] André M. Bastos, Jacob A. Donoghue, Scott L. Brincat, Meredith Mahnke, Jorge Yanar, Josefina Correa, Ayan S. Waite, Mikael Lundqvist, Jefferson Roy, Emery N. Brown, and Earl K. Miller. Neural effects of propofol-induced unconsciousness and its reversal using thalamic stimulation. eLife, 10, 2021. ISSN 2050084X. doi:10.7554/ELIFE.60824. PMID: 33904411 publisher: eLife Sciences Publications Ltd.

[90] Mohsen Afrasiabi, Michelle J. Redinbaugh, Jessica M. Phillips, Niranjan A. Kambi, Sounak Mohanta, Aeyal Raz, Andrew M. Haun, and Yuri B. Saalmann. Consciousness depends on integration between parietal cortex, striatum, and thalamus. Cell Systems, 12(4): 363–373.e11, apr 21 2021. ISSN 2405-4720. doi: 10.1016/j.cels.2021.02.003. PMID: 33730543 PMCID: PMC8084606.

[91] George A. Mashour. Anesthesia and the neurobiology of consciousness. Neuron, 0(0), apr 4 2024. ISSN 0896-6273. doi:10.1016/j.neuron.2024.03.002. URL https://www.cell.com/neuron/abstract/S0896-6273(24)00156-9. publisher: Elsevier PMID: 38579714.

[93] Klaus H Maier-Hein, Peter F Neher, Jean-Christophe Houde, Marc-Alexandre Côté, Eleftherios Garyfallidis, Jidan Zhong, Maxime Chamberland, Fang-Cheng Yeh, Ying-Chia Lin, Qing Ji, et al. The challenge of mapping the human connectome based on diffusion tractography. Nature communications, 8(1):1349, 2017.

[94] Jesper Ekelund, Mark Slifstein, Raj Narendran, Olivier Guillin, Hemant Belani, Ning-Ning Guo, Yuying Hwang, Dah-Ren Hwang, Anissa Abi-Dargham, and Marc Laruelle. In vivo da d 1 receptor selectivity of nnc 112 and sch 23390. Molecular imaging and biology, 9(3):117–125, 2007.

[95] Alexander Schaefer, Ru Kong, Evan M Gordon, Timothy O Laumann, Xi-Nian Zuo, Avram J Holmes, Simon B Eickhoff, and BT Thomas Yeo. Local-global parcellation of the human cerebral cortex from intrinsic functional connectivity mri. Cerebral cortex, 28(9):3095–3114, 2018.

[96] Peter T. Fox and Jack L. Lancaster. Opinion: Mapping context and content: the BrainMap model. Nature Reviews. Neuroscience, 3(4):319–321, April 2002. ISSN 1471-003X. doi:10.1038/nrn789.

[97] Lorina Naci, Leah Sinai, and Adrian M Owen. Detecting and interpreting conscious experiences in behaviorally non-responsive patients. NeuroImage, 145:304–313, 2017. ISSN 10959572. doi: 10.1016/j.neuroimage.2015.11.059. PMID: 26679327 Citation Key: Naci2015.

[98] Lorina Naci, Amelie Haugg, Alex MacDonald, Mimma Anello, Evan Houldin, Shakib Naqshbandi, Laura E Gonzalez-Lara, Miguel Arango, Christopher Harle, Rhodri Cusack, and Adrian M Owen. Functional diversity of brain networks supports consciousness and verbal intelligence. Scientific Reports, 8(1):1–15, 2018. ISSN 20452322. doi:10.1038/s41598-018-31525-z. PMID: 30185912.

[99] Andrea I. Luppi, Jakub Vohryzek, Morten L. Kringelbach, Pedro A. M. Mediano, Michael M. Craig, Ram Adapa, Robin L. Carhart-Harris, Leor Roseman, Ioannis Pappas, Alexander R. D. Peattie, Anne E. Manktelow, Barbara J. Sahakian, Paola Finoia, Guy B. Williams, Judith Allanson, John D. Pickard, David K. Menon, Selen Atasoy, and Emmanuel A. Stamatakis. Distributed harmonic patterns of structure-function dependence orchestrate human consciousness. Communications Biology, 6(1):1– 19, 1 2023. ISSN 2399-3642. doi:10.1038/s42003-023-04474-1. number: 1 publisher: Nature Publishing Group.

[100] Marie-Aurélie Bruno, Audrey Vanhaudenhuyse, Aurore Thibaut, Gustave Moonen, and Steven Laureys. From unresponsive wakefulness to minimally conscious plus and functional locked-in syndromes: Recent advances in our understanding of disorders of consciousness. Journal of Neurology, 258:1373–1384, a2011.

[101] Sophie Wannez et al. Prevalence of coma-recovery scale-revised signs of consciousness in patients in minimally conscious state. Neuropsychological Rehabilitation, 28: 1350–1359, 2018.

[102] Anne E Manktelow, David K Menon, Barbara J Sahakian, and Emmanuel A Stamatakis. Working memory after traumatic brain injury: The neural basis of improved performance with methylphenidate. Frontiers in Behavioral Neuroscience, 11, 2017. ISSN 1662-5153. doi:10.3389/fnbeh.2017.00058. URL http://journal.frontiersin.org/article/10.3389/fnbeh.2017.00058/full. PMID: 28424597 Citation Key: Manktelow2017.

[103] Susan Whitfield-Gabrieli and Alfonso Nieto-Castanon. Conn: A Functional Connectivity Toolbox for Correlated and Anticorrelated Brain Networks. Brain Connectivity, 2(3):125–141, 2012. ISSN 21580022. doi: 10.1089/brain.2012.0073. Citation Key: Whitfield-Gabrieli.

[104] Behzadi Y, Restom K, Liau J, and Liu TT. A component based noise correction method (compcor) for bold and perfusion based fmri. NeuroImage, 37:90–101, 2007.

[105] Jonathan D Power, Kelly A Barnes, Abraham Z Snyder, Bradley L Schlaggar, and Steven E Petersen. Spurious but systematic correlations in functional connectivity mri networks arise from subject motion. NeuroImage, 59(3):2142–2154, 2012. ISSN 10538119. doi: 10.1016/j.neuroimage.2011.10.018. PMID: 22019881 arXiv: NIHMS150003 Citation Key: Power2012 ISBN: 1095-9572 (Electronic)\r1053-8119 (Linking).

[106] Jonathan D Power, Anish Mitra, Timothy O Laumann, Abraham Z Snyder, Bradley L Schlaggar, and Steven E Petersen. Methods to detect, characterize, and remove motion artifact in resting state fmri. NeuroImage, 84:320–341, 2014. ISSN 10538119. doi: 10.1016/j.neuroimage.2013.08.048. PMID: 23994314.

[107] Kevin Murphy and Michael D. Fox. Towards a consensus regarding global signal regression for resting state functional connectivity mri. NeuroImage, 154:169–173, 7 2017. ISSN 10959572. doi: 10.1016/j.neuroimage.2016.11.052. PMID: 27888059 publisher: Neuroimage.

[108] Jingwei Li, Taylor Bolt, Danilo Bzdok, Jason S Nomi, BT Thomas Yeo, R Nathan Spreng, and Lucina Q Ud-din. Topography and behavioral relevance of the global signal in the human brain. Scientific reports, 9(1):14286, 2019.

[109] Sean Tanabe, Zirui Huang, Jun Zhang, Yali Chen, Stuart Fogel, Julien Doyon, Jinsong Wu, Jianghui Xu, Jianfeng Zhang, Pengmin Qin, Xuehai Wu, Ying Mao, George A. Mashour, Anthony G. Hudetz, and Georg Northoff. Altered global brain signal during physiologic, pharmacologic, and pathologic states of unconsciousness in humans and rats. Anesthesiology, pages 1392–1406, 2020. ISSN 15281175. doi: 10.1097/ALN.0000000000003197.

[110] Rahul S Desikan, Florent Ségonne, Bruce Fischl, Brian T Quinn, Bradford C Dickerson, Deborah Blacker, Randy L Buckner, Anders M Dale, R Paul Maguire, Bradley T Hyman, et al. An automated labeling system for subdividing the human cerebral cortex on mri scans into gyral based regions of interest. Neuroimage, 31(3):968–980, 2006.

[111] Mark Jenkinson, Peter Bannister, Michael Brady, and Stephen Smith. Improved optimization for the robust and accurate linear registration and motion correction of brain images. Neuroimage, 17(2):825–841, 2002.

[112] Theodore D. Satterthwaite, Mark A. Elliott, Raphael T. Gerraty, Kosha Ruparel, James Loughead, Monica E. Calkins, Simon B. Eickhoff, et al. An improved framework for confound regression and filtering for control of motion artifact in the preprocessing of resting-state functional connectivity data. NeuroImage, 64:240–256, 2013. doi:10.1016/j.neuroimage.2012.08.052.

[113] Rémi Patriat, Richard C. Reynolds, and Rasmus M. Birn. An improved model of motion-related signal changes in fmri. NeuroImage, 144:74–82, 2017. doi: 10.1016/j.neuroimage.2016.08.051.

[114] Sharna D Jamadar, Phillip GD Ward, Emma X Liang, Edwina R Orchard, Zhaolin Chen, and Gary F Egan. Metabolic and hemodynamic resting-state connectivity of the human brain: a high-temporal resolution simultaneous bold-fmri and fdg-fpet multimodality study. Cerebral Cortex, 31(6):2855–2867, 2021.

[115] Sharna D Jamadar, Phillip GD Ward, Emma X Liang, Edwina R Orchard, Zhaolin Chen, and Gary F Egan. Metabolic and hemodynamic resting-state connectivity of the human brain: a high-temporal resolution simultaneous bold-fmri and fdg-fpet multimodality study. Cerebral Cortex, 31(6):2855–2867, 2021.

[116] Katharina Voigt, Emma X Liang, Bratislav Misic, Phillip G D Ward, Gary F Egan, and Sharna D Jamadar. Metabolic and functional connectivity provide unique and complementary insights into cognition-connectome relationships. Cerebral Cortex, 2022.

[117] Ninon Burgos, M Jorge Cardoso, Kris Thielemans, Marc Modat, Stefano Pedemonte, John Dickson, Anna Barnes, Rebekah Ahmed, Colin J Mahoney, Jonathan M Schott, et al. Attenuation correction synthesis for hybrid pet-mr scanners: application to brain studies. IEEE transactions on medical imaging, 33(12):2332–2341, 2014.

[118] Sharna D Jamadar, Phillip GD Ward, Thomas G Close, Alex Fornito, Malin Premaratne, Kieran O’Brien, Daniel Stäb, Zhaolin Chen, N Jon Shah, and Gary F Egan. Simultaneous bold-fmri and constant infusion fdg-pet data of the resting human brain. Scientific data, 7(1):1–12, 2020.

[119] David C Van Essen, Stephen M Smith, Deanna M Barch, Timothy EJ Behrens, Essa Yacoub, Kamil Ugurbil, Wu-Minn HCP Consortium, et al. The wu-minn human connectome project: an overview. Neuroimage, 80:62–79, 2013.

[120] François Tadel, Sylvain Baillet, John C Mosher, Dimitrios Pantazis, and Richard M Leahy. Brainstorm: a user-friendly application for meg/eeg analysis. Computational intelligence and neuroscience, 2011, 2011.

[121] Giles L Colclough, Mark W Woolrich, PK Tewarie, Matthew J Brookes, Andrew J Quinn, and Stephen M Smith. How reliable are meg resting-state connectivity metrics? Neuroimage, 138:284–293, 2016.

[122] Meher R Juttukonda, Binyin Li, Randa Almaktoum, Kimberly A Stephens, Kathryn M Yochim, Essa Yacoub, Randy L Buckner, and David H Salat. Characterizing cerebral hemodynamics across the adult lifespan with arterial spin labeling mri data from the human connectome project-aging. Neuroimage, 230:117807, 2021.

[123] Thomas F. Kirk, Flora A. Kennedy McConnell, Jack Toner, Martin S. Craig, Davide Carone, Xiufeng Li, Yuriko Suzuki, Timothy S. Coalson, Michael P. Harms, Matthew F. Glasser, and Michael A. Chappell. Arterial spin labelling perfusion mri analysis for the human connectome project lifespan ageing and development studies. Imaging Neuroscience, 3, 01 2025. ISSN 2837-6056. doi:10.1162/imag_a_00444.

[124] Michael P Harms, Leah H Somerville, Beau M Ances, Jesper Andersson, Deanna M Barch, Matteo Bastiani, Susan Y Bookheimer, Timothy B Brown, Randy L Buckner, Gregory C Burgess, et al. Extending the human connectome project across ages: Imaging protocols for the lifespan development and aging projects. Neuroimage, 183:972–984, 2018.

[125] Leah H Somerville, Susan Y Bookheimer, Randy L Buckner, Gregory C Burgess, Sandra W Curtiss, Mirella Dapretto, Jennifer Stine Elam, Michael S Gaffrey, Michael P Harms, Cynthia Hodge, et al. The lifespan human connectome project in development: A large-scale study of brain connectivity development in 5–21 year olds. Neuroimage, 183:456–468, 2018.

[126] Susan Y Bookheimer, David H Salat, Melissa Terpstra, Beau M Ances, Deanna M Barch, Randy L Buckner, Gregory C Burgess, Sandra W Curtiss, Mirella Diaz-Santos, Jennifer Stine Elam, et al. The lifespan human connectome project in aging: an overview. Neuroimage, 185: 335–348, 2019.

[127] Jie Lisa Ji, Jure Demšar, Clara Fonteneau, Zailyn Tamayo, Lining Pan, Aleksij Kraljič, Andraž Matkovič, Nina Purg, Markus Helmer, Shaun Warrington, et al. Qunex—an integrative platform for reproducible neuroimaging analytics. Frontiers in Neuroinformatics, 17: 1104508, 2023.

[128] Yuriko Suzuki, Thomas W Okell, Michael A Chappell, and Matthias JP van Osch. A framework for motion correction of background suppressed arterial spin labeling perfusion images acquired with simultaneous multi-slice epi. Magnetic resonance in medicine, 81(3):1553–1565, 2019.

[129] Michael A Chappell, Thomas F Kirk, Martin S Craig, Flora A Kennedy McConnell, Moss Y Zhao, Bradley J MacIntosh, Thomas W Okell, and Mark W Woolrich. Basil: A toolbox for perfusion quantification using arterial spin labelling. Imaging Neuroscience, 1:1–16, 2023.

[130] Richard B Buxton, Lawrence R Frank, Eric C Wong, Bettina Siewert, Steven Warach, and Robert R Edelman. A general kinetic model for quantitative perfusion imaging with arterial spin labeling. Magnetic resonance in medicine, 40(3):383–396, 1998.

[131] Michael A Chappell, Adrian R Groves, Bradley J MacIntosh, Manus J Donahue, Peter Jezzard, and Mark W Woolrich. Partial volume correction of multiple inversion time arterial spin labeling mri data. Magnetic resonance in medicine, 65(4):1173–1183, 2011.

[132] Joana Pinto, Michael A Chappell, Thomas W Okell, Melvin Mezue, Andrew R Segerdahl, Irene Tracey, Pedro Vilela, and Patrícia Figueiredo. Calibration of arterial spin labeling data—potential pitfalls in postprocessing. Magnetic resonance in medicine, 83(4):1222–1234, 2020.

[133] Bruce Fischl. Freesurfer. Neuroimage, 62(2):774–781, 2012.

[134] Thomas F Kirk, Timothy S Coalson, Martin S Craig, and Michael A Chappell. Toblerone: surface-based partial volume estimation. IEEE transactions on medical imaging, 39(5):1501–1510, 2019.

[135] Matthew F Glasser, Stamatios N Sotiropoulos, J Anthony Wilson, Timothy S Coalson, Bruce Fischl, Jesper L Andersson, Junqian Xu, Saad Jbabdi, Matthew Webster, Jonathan R Polimeni, et al. The minimal preprocessing pipelines for the human connectome project. Neuroimage, 80:105–124, 2013.

[136] Emma C Robinson, Saad Jbabdi, Matthew F Glasser, Jesper Andersson, Gregory C Burgess, Michael P Harms, Stephen M Smith, David C Van Essen, and Mark Jenkinson. Msm: a new flexible framework for multimodal surface matching. Neuroimage, 100:414–426, 2014.

[137] Matthew F Glasser, Timothy S Coalson, Emma C Robinson, Carl D Hacker, John Harwell, Essa Yacoub, Kamil Ugurbil, Jesper Andersson, Christian F Beckmann, Mark Jenkinson, et al. A multi-modal parcellation of human cerebral cortex. Nature, 536(7615):171–178, 2016.

[138] Emma C Robinson, Kara Garcia, Matthew F Glasser, Zhengdao Chen, Timothy S Coalson, Antonios Makropoulos, Jelena Bozek, Robert Wright, Andreas Schuh, Matthew Webster, et al. Multimodal surface matching with higher-order smoothness constraints. Neuroimage, 167:453–465, 2018.

[139] Andrea I Luppi, Zhen-Qi Liu, Justine Y Hansen, Rodrigo Cofre, Meiqi Niu, Elena Kuzmin, Seán Froudist-Walsh, Nicola Palomero-Gallagher, and Bratislav Misic. Benchmarking macaque brain gene expression for horizontal and vertical translation. Science Advances, 11(9): eads6967, 2025.

[140] Ross D Markello, Aurina Arnatkeviciute, Jean-Baptiste Poline, Ben D Fulcher, Alex Fornito, and Bratislav Misic. Standardizing workflows in imaging transcriptomics with the abagen toolbox. Elife, 10:e72129, 2021.

[141] Simon Kaller, Michael Rullmann, Marianne Patt, Georg Becker, Julia Luthardt, Johanna Girbardt, Philipp Meyer, Peter Werner, Henryk Barthel, Anke McLeod, Thomas Fritz, Swen Hesse, and Osama Sabri. Test–retest measurements of dopamine d1-type receptors using simultaneous pet/mri imaging. European Journal of Nuclear Medicine and Molecular Imaging, 44, 6 2017. doi: 10.1007/s00259-017-3645-0.

[142] Christine M. Sandiego, Jean-Dominique Gallezot, Keunpoong Lim, Jim Ropchan, Shu-fei Lin, Hong Gao, Evan D. Morris, and Kelly P. Cosgrove. Reference region modeling approaches for amphetamine challenge studies with [11c]flb 457 and pet. Journal of Cerebral Blood Flow and Metabolism: Official Journal of the International Society of Cerebral Blood Flow and Metabolism, 35(4):623–629, 3 2015. ISSN 1559-7016. doi:10.1038/jcbfm.2014.237. PMID: 25564239 PMCID: PMC4420880.

[143] Mark Slifstein, Elsmarieke van de Giessen, Jared Van Snellenberg, Judy L. Thompson, Rajesh Narendran, Roberto Gil, Elizabeth Hackett, Ragy Girgis, Najate Ojeil, Holly Moore, Deepak D’Souza, Robert T. Malison, Yiyun Huang, Keunpoong Lim, Nabeel Nabulsi, Richard E. Carson, Jeffrey A. Lieberman, and Anissa Abi-Dargham. Deficits in prefrontal cortical and extrastriatal dopamine release in schizophrenia: a positron emission tomographic functional magnetic resonance imaging study. JAMA psychiatry, 72(4):316–324, 4 2015. ISSN 2168-6238. doi:10.1001/jamapsychiatry.2014.2414. PMID: 25651194 PMCID: PMC4768742.

[144] Christopher T. Smith, Jennifer L. Crawford, Linh C. Dang, Kendra L. Seaman, M. Danica San Juan, Aishwarya Vijay, Daniel T. Katz, David Matuskey, Ronald L. Cowan, Evan D. Morris, David H. Zald, and Gregory R. Samanez-Larkin. Partial-volume correction increases estimated dopamine d2-like receptor binding potential and reduces adult age differences. Journal of Cerebral Blood Flow and Metabolism: Official Journal of the International Society of Cerebral Blood Flow and Metabolism, 39(5):822–833, 5 2019. ISSN 1559-7016. doi:10.1177/0271678X17737693. PMID: 29090626 PMCID: PMC6498753.

[145] Yasmin Zakiniaeiz, Ansel T. Hillmer, David Matuskey, Nabeel Nabulsi, Jim Ropchan, Carolyn M. Mazure, Marina R. Picciotto, Yiyun Huang, Sherry A. McKee, Evan D. Morris, and Kelly P. Cosgrove. Sex differences in amphetamine-induced dopamine release in the dorsolateral prefrontal cortex of tobacco smokers. Neuropsychopharmacology: Official Publication of the American College of Neuropsychopharmacology, 44(13):2205– 2211, 12 2019. ISSN 1740-634X. doi:10.1038/s41386-019-0456-y. PMID: 31269510 PMCID: PMC6897943.

[146] Juergen Dukart, Štefan Holiga, Christopher Chatham, Peter Hawkins, Anna Forsyth, Rebecca McMillan, Jim Myers, Anne R. Lingford-Hughes, David J. Nutt, Emilio Merlo-Pich, Celine Risterucci, Lauren Boak, Daniel Um-bricht, Scott Schobel, Thomas Liu, Mitul A. Mehta, Fernando O. Zelaya, Steve C. Williams, Gregory Brown, Martin Paulus, Garry D. Honey, Suresh Muthukumaraswamy, Joerg Hipp, Alessandro Bertolino, and Fabio Sambataro. Cerebral blood flow predicts differential neurotransmitter activity. Scientific Reports, 8(1): 4074, 3 2018. ISSN 2045-2322. doi:10.1038/s41598-018-22444-0. PMID: 29511260 PMCID: PMC5840131.

[147] Renata Belfort-DeAguiar, Jean-Dominique Gallezot, Janice J Hwang, Ahmed Elshafie, Catherine W Yeckel, Owen Chan, Richard E Carson, Yu-Shin Ding, and Robert S Sherwin. Noradrenergic activity in the human brain: A mechanism supporting the defense against hypoglycemia. The Journal of Clinical Endocrinology and Metabolism, 103(6):2244–2252, 6 2018. ISSN 0021-972X. doi:10.1210/jc.2017-02717.

[148] Yu-Shin Ding, Tarun Singhal, Beata Planeta-Wilson, Jean-Dominique Gallezot, Nabeel Nabulsi, David Labaree, Jim Ropchan, Shannan Henry, Wendol Williams, Richard E. Carson, Alexander Neumeister, and Robert T. Malison. Pet imaging of the effects of age and cocaine on the norepinephrine transporter in the human brain using (s,s)-[(11)c]o-methylreboxetine and hrrt. Synapse (New York, N.Y.), 64(1):30–38, 1 2010. ISSN 1098-2396. doi:10.1002/syn.20696. PMID: 19728366 PMCID: PMC3727644.

[149] Chiang-shan R. Li, Marc N. Potenza, Dianne E. Lee, Beata Planeta, Jean-Dominique Gallezot, David Labaree, Shannan Henry, Nabeel Nabulsi, Rajita Sinha, Yu-Shin Ding, Richard E. Carson, and Alexander Neumeister. Decreased norepinephrine transporter availability in obesity: Positron emission tomography imaging with (s,s)-[(11)c]o-methylreboxetine. NeuroImage, 86:306–310, 2 2014. ISSN 1095-9572. doi: 10.1016/j.neuroimage.2013.10.004. PMID: 24121204 PMCID: PMC3947246.

[150] Elizabeth Sanchez-Rangel, Jean-Dominique Gallezot, Catherine W. Yeckel, Wai Lam, Renata Belfort-DeAguiar, Ming-Kai Chen, Richard E. Carson, Robert Sherwin, and Janice J. Hwang. Norepinephrine transporter availability in brown fat is reduced in obesity: a human pet study with [11c] mrb. International Journal of Obesity, 44(4):964–967, 4 2020. ISSN 1476-5497. doi: 10.1038/s41366-019-0471-4. number: 4 publisher: Nature Publishing Group.

[151] Markus Savli, Andreas Bauer, Markus Mitterhauser, Yu-Shin Ding, Andreas Hahn, Tina Kroll, Alexander Neumeister, Daniela Haeusler, Johanna Ungersboeck, Shannan Henry, Sanaz Attaripour Isfahani, Frank Rattay, Wolfgang Wadsak, Siegfried Kasper, and Rupert Lanzenberger. Normative database of the serotonergic system in healthy subjects using multi-tracer pet. NeuroImage, 63(1):447–459, 10 2012. ISSN 1053-8119. doi: 10.1016/j.neuroimage.2012.07.001.

[152] Jean-Dominique Gallezot, Nabeel Nabulsi, Alexander Neumeister, Beata Planeta-Wilson, Wendol A. Williams, Tarun Singhal, Sunhee Kim, R. Paul Maguire, Timothy McCarthy, J. James Frost, Yiyun Huang, Yu-Shin Ding, and Richard E. Carson. Kinetic modeling of the serotonin 5-ht(1b) receptor radioligand [(11)c]p943 in humans. Journal of Cerebral Blood Flow and Metabolism: Official Journal of the International Society of Cerebral Blood Flow and Metabolism, 30(1):196–210, 1 2010. ISSN 1559-7016. doi:10.1038/jcbfm.2009.195. PMID: 19773803 PMCID: PMC2949107.

[153] David Matuskey, Zubin Bhagwagar, Beata Planeta, Brian Pittman, Jean-Dominique Gallezot, Jason Chen, Jane Wanyiri, Soheila Najafzadeh, Jim Ropchan, Paul Geha, Yiyun Huang, Marc N. Potenza, Alexander Neumeister, Richard E. Carson, and Robert T. Malison. Reductions in brain 5-ht1b receptor availability in primarily cocaine-dependent humans. Biological Psychiatry, 76(10):816–822, 11 2014. ISSN 1873-2402. doi: 10.1016/j.biopsych.2013.11.022. PMID: 24433854 PMCID: PMC4037398.

[154] James W. Murrough, Christoph Czermak, Shannan Henry, Nabeel Nabulsi, Jean-Dominique Gallezot, Ralitza Gueorguieva, Beata Planeta-Wilson, John H. Krystal, John F. Neumaier, Yiyun Huang, Yu-Shin Ding, Richard E. Carson, and Alexander Neumeister. The effect of early trauma exposure on serotonin type 1b receptor expression revealed by reduced selective radioligand binding. Archives of General Psychiatry, 68(9):892–900, 9 2011. ISSN 1538-3636. doi: 10.1001/archgenpsychiatry.2011.91. PMID: 21893657 PMCID: PMC3244836.

[155] Christopher Pittenger, Thomas G. Adams, Jean-Dominique Gallezot, Michael J. Crowley, Nabeel Nabulsi, null James Ropchan, Hong Gao, Stephen A. Kichuk, Ryan Simpson, Eileen Billingslea, Jonas Hannestad, Michael Bloch, Linda Mayes, Zubin Bhagwagar, and Richard E. Carson. Ocd is associated with an altered association between sensorimotor gating and cortical and subcortical 5-ht1b receptor binding. Journal of Affective Disorders, 196:87–96, 5 2016. ISSN 1573-2517. doi: 10.1016/j.jad.2016.02.021. PMID: 26919057 PMCID: PMC4808438.

[156] Aybala Saricicek, Jason Chen, Beata Planeta, Barbara Ruf, Kalyani Subramanyam, Kathleen Maloney, David Matuskey, David Labaree, Lorenz Deserno, Alexander Neumeister, John H. Krystal, Jean-Dominique Gallezot, Yiyun Huang, Richard E. Carson, and Zubin Bhagwagar. Test-retest reliability of the novel 5-ht1b receptor pet radioligand [11c]p943. European Journal of Nuclear Medicine and Molecular Imaging, 42(3):468–477, 3 2015. ISSN 1619-7089. doi:10.1007/s00259-014-2958-5. PMID: 25427881.

[157] Vincent Beliveau, Melanie Ganz, Ling Feng, Brice Ozenne, Liselotte Højgaard, Patrick M. Fisher, Claus Svarer, Douglas N. Greve, and Gitte M. Knudsen. A high-resolution in vivo atlas of the human brain’s serotonin system. The Journal of Neuroscience, 37(1):120, 1 2017. doi:10.1523/JNEUROSCI.2830-16.2016. PMID: 28053035 publisher: Society for Neuroscience.

[158] Rajiv Radhakrishnan, Nabeel Nabulsi, Edward Gaiser, Jean-Dominique Gallezot, Shannan Henry, Beata Planeta, Shu-fei Lin, Jim Ropchan, Wendol Williams, Evan Morris, Deepak Cyril D’Souza, Yiyun Huang, Richard E. Carson, and David Matuskey. Age-related change in 5-ht6 receptor availability in healthy male volunteers measured with 11c-gsk215083 pet. Journal of Nuclear Medicine, 59(9):1445–1450, 9 2018. ISSN 0161-5505. doi:10.2967/jnumed.117.206516. PMID: 29626125 PMCID: PMC6126437.

[159] Rajiv Radhakrishnan, David Matuskey, Nabeel Nabulsi, Edward Gaiser, Jean-Dominique Gallezot, Shannan Henry, Beata Planeta, Shu-Fei Lin, Jim Ropchan, Yiyun Huang, Richard E. Carson, and Deepak Cyril D’Souza. In vivo 5-ht6 and 5-ht2a receptor availability in antipsychotic treated schizophrenia patients vs. unmedicated healthy humans measured with [11c]gsk215083 pet. Psychiatry Research. Neuroimaging, 295:111007, 1 2020. ISSN 1872-7506. doi: 10.1016/j.pscychresns.2019.111007. PMID: 31760336.

[160] Stephen R. Baldassarri, Ansel T. Hillmer, Jon Mikael Anderson, Peter Jatlow, Nabeel Nabulsi, David Labaree, Kelly P. Cosgrove, Stephanie S. O’Malley, Thomas Eissenberg, Suchitra Krishnan-Sarin, and Irina Esterlis. Use of electronic cigarettes leads to significant beta2-nicotinic acetylcholine receptor occupancy: Evidence from a pet imaging study. Nicotine and Tobacco Research: Official Journal of the Society for Research on Nicotine and Tobacco, 20(4):425–433, 3 2018. ISSN 1469-994X. doi:10.1093/ntr/ntx091. PMID: 28460123 PMCID: PMC5896427.

[161] A. T. Hillmer, I. Esterlis, J. D. Gallezot, F. Bois, M. Q. Zheng, N. Nabulsi, S. F. Lin, R. L. Papke, Y. Huang, O. Sabri, R. E. Carson, and K. P. Cosgrove. Imaging of cerebral alpha4beta2* nicotinic acetylcholine receptors with (-)-[(18)f]flubatine pet: Implementation of bolus plus constant infusion and sensitivity to acetylcholine in human brain. NeuroImage, 141:71–80, 11 2016. ISSN 1095-9572. doi:10.1016/j.neuroimage.2016.07.026. PMID: 27426839 PMCID: PMC5026941.

[162] Mika Naganawa, Nabeel Nabulsi, Shannan Henry, David Matuskey, Shu-Fei Lin, Lawrence Slieker, Adam J. Schwarz, Nancy Kant, Cynthia Jesudason, Kevin Ru-ley, Antonio Navarro, Hong Gao, Jim Ropchan, David Labaree, Richard E. Carson, and Yiyun Huang. First-in-human assessment of 11c-lsn3172176, an m1 muscarinic acetylcholine receptor pet radiotracer. Journal of Nuclear Medicine: Official Publication, Society of Nuclear Medicine, 62(4):553–560, 4 2021. ISSN 1535-5667. doi: 10.2967/jnumed.120.246967. PMID: 32859711 PMCID: PMC8049371.

[163] M. Aghourian, C. Legault-Denis, J.-P. Soucy, P. Rosa-Neto, S. Gauthier, A. Kostikov, P. Gravel, and M.-A. Bédard. Quantification of brain cholinergic denervation in alzheimer’s disease using pet imaging with [18f]-feobv. Molecular Psychiatry, 22(11):1531–1538, 11 2017. ISSN 1476-5578. doi:10.1038/mp.2017.183. PMID: 28894304.

[164] Marc-Andre Bedard, Meghmik Aghourian, Camille Legault-Denis, Ronald B. Postuma, Jean-Paul Soucy, Jean-François Gagnon, Amélie Pelletier, and Jacques Montplaisir. Brain cholinergic alterations in idiopathic rem sleep behaviour disorder: a pet imaging study with 18f-feobv. Sleep Medicine, 58:35–41, 6 2019. ISSN 1878-5506. doi:10.1016/j.sleep.2018.12.020. PMID: 31078078.

[165] Jonathan M. DuBois, Olivier G. Rousset, Jared Rowley, Manuel Porras-Betancourt, Andrew J. Reader, Aurelie Labbe, Gassan Massarweh, Jean-Paul Soucy, Pedro Rosa-Neto, and Eliane Kobayashi. Characterization of age/sex and the regional distribution of mglur5 availability in the healthy human brain measured by high-resolution [(11)c]abp688 pet. European Journal of Nuclear Medicine and Molecular Imaging, 43(1):152–162, 1 2016. ISSN 1619-7089. doi:10.1007/s00259-015-3167-6. PMID: 26290423.

[166] Kelly Smart, Sylvia M. L. Cox, Stephanie G. Scala, Maria Tippler, Natalia Jaworska, Michel Boivin, Jean R. Séguin, Chawki Benkelfat, and Marco Leyton. Sex differences in [11c]abp688 binding: a positron emission tomography study of mglu5 receptors. European Journal of Nuclear Medicine and Molecular Imaging, 46(5):1179–1183, 2019. ISSN 1619-7070. doi:10.1007/s00259-018-4252-4. PMID: 30627817 PMCID: PMC6451701.

[167] Marian Galovic, Adam Al-Diwani, Umesh Vivekananda, Francisco Torrealdea, Kjell Erlandsson, Tim D. Fryer, Young T. Hong, Benjamin A. Thomas, Colm J. McGinnity, Evan Edmond, Kerstin Sander, Erik Årstad, Ilijas Jelcic, Franklin I. Aigbirhio, Ashley M. Groves, Kris Thielemans, Brian Hutton, Alexander Hammers, John S. Duncan, Jonathan P. Coles, Anna Barnes, Charlotte J. Stagg, Matthew C. Walker, Sarosh R. Irani, Matthias J. Koepp, and for the NEST Investigators. In vivo nmda receptor function in people with nmda receptor antibody encephalitis. medRxiv, 12 2021. doi:10.1101/2021.12.04.21267226. URL https://www.medrxiv.org/content/10.1101/2021.12.04.21267226v1. page: 2021.12.04.21267226.

[168] Marian Galovic, Kjell Erlandsson, Tim D. Fryer, Young T. Hong, Roido Manavaki, Hasan Sari, Sarah Chetcuti, Benjamin A. Thomas, Martin Fisher, Selena Sephton, Roberto Canales, Joseph J. Russell, Kerstin Sander, Erik Årstad, Franklin I. Aigbirhio, Ashley M. Groves, John S. Duncan, Kris Thielemans, Brian F. Hutton, Jonathan P. Coles, Matthias J. Koepp, and NEST investigators. Validation of a combined image derived input function and venous sampling approach for the quantification of [18f]ge-179 pet binding in the brain. NeuroImage, 237:118194, 8 2021. ISSN 1095-9572. doi: 10.1016/j.neuroimage.2021.118194. PMID: 34023451.

[169] Martin Nørgaard, Vincent Beliveau, Melanie Ganz, Claus Svarer, Lars Pinborg, Sune Keller, Peter Jensen, Douglas Greve, and Gitte Knudsen. A high-resolution in vivo atlas of the human brain’s benzodiazepine binding site of gaba a receptors. NeuroImage, 232:117878, 2021. doi: 10.1101/2020.04.10.035352. ISBN: 4535456720.

[170] Jean-Dominique Gallezot, Beata Planeta, Nabeel Nabulsi, Donna Palumbo, Xiaoxi Li, Jing Liu, Carolyn Rowinski, Kristin Chidsey, David Labaree, Jim Ropchan, Shu-Fei Lin, Aarti Sawant-Basak, Timothy J. McCarthy, Anne W. Schmidt, Yiyun Huang, and Richard E. Carson. Determination of receptor occupancy in the presence of mass dose: [11c]gsk189254 pet imaging of histamine h3 receptor occupancy by pf-03654746. Journal of Cerebral Blood Flow and Metabolism: Official Journal of the International Society of Cerebral Blood Flow and Metabolism, 37(3):1095–1107, 3 2017. ISSN 1559-7016. doi:10.1177/0271678X16650697. PMID: 27207170 PMCID: PMC5363483.

[171] Deepak Cyril D’Souza, Jose A. Cortes-Briones, Mohini Ranganathan, Halle Thurnauer, Gina Creatura, Toral Surti, Beata Planeta, Alexander Neumeister, Brian Pittman, Marc Normandin, Michael Kapinos, Jim Ropchan, Yiyun Huang, Richard E. Carson, and Patrick D. Skosnik. Rapid changes in cb1 receptor availability in cannabis dependent males after abstinence from cannabis. Biological Psychiatry. Cognitive Neuroscience and Neuroimaging, 1(1):60–67, 1 2016. ISSN 2451-9022. doi:10.1016/j.bpsc.2015.09.008. PMID: 26858993 PMCID: PMC4742341.

[172] Alexander Neumeister, Marc D. Normandin, James W. Murrough, Shannan Henry, Christopher R. Bailey, David A. Luckenbaugh, Keri Tuit, Ming-Qiang Zheng, Isaac R. Galatzer-Levy, Rajita Sinha, Richard E. Carson, Marc N. Potenza, and Yiyun Huang. Positron emission tomography shows elevated cannabinoid cb1 receptor binding in men with alcohol dependence. Alcoholism, Clinical and Experimental Research, 36 (12):2104–2109, 12 2012. ISSN 1530-0277. doi: 10.1111/j.1530-0277.2012.01815.x. PMID: 22551199 PMCID: PMC3418442.

[173] Marc D. Normandin, Ming-Qiang Zheng, Kuo-Shyan Lin, N. Scott Mason, Shu-Fei Lin, Jim Ropchan, David Labaree, Shannan Henry, Wendol A. Williams, Richard E. Carson, Alexander Neumeister, and Yiyun Huang. Imaging the cannabinoid cb1 receptor in humans with [11c]omar: assessment of kinetic analysis methods, testretest reproducibility, and gender differences. Journal of Cerebral Blood Flow and Metabolism: Official Journal of the International Society of Cerebral Blood Flow and Metabolism, 35(8):1313–1322, 8 2015. ISSN 1559-7016. doi:10.1038/jcbfm.2015.46. PMID: 25833345 PMCID: PMC4528005.

[174] Mohini Ranganathan, Jose Cortes-Briones, Rajiv Radhakrishnan, Halle Thurnauer, Beata Planeta, Patrick Skosnik, Hong Gao, David Labaree, Alexander Neumeister, Brian Pittman, Toral Surti, Yiyun Huang, Richard E. Carson, and Deepak Cyril D’Souza. Reduced brain cannabinoid receptor availability in schizophrenia. Biological Psychiatry, 79(12):997–1005, 6 2016. ISSN 1873-2402. doi:10.1016/j.biopsych.2015.08.021. PMID: 26432420 PMCID: PMC4884543.

[175] Tatu Kantonen, Tomi Karjalainen, Janne Isojärvi, Pirjo Nuutila, Jouni Tuisku, Juha Rinne, Jarmo Hietala, Valtteri Kaasinen, Kari Kalliokoski, Harry Scheinin, Jussi Hirvonen, Aki Vehtari, and Lauri Nummenmaa. Interindividual variability and lateralization of mu-opioid receptors in the human brain. NeuroImage, 217:116922, 8 2020. ISSN 1053-8119. doi: 10.1016/j.neuroimage.2020.116922.

[176] Casey Paquola, Jessica Royer, Lindsay B Lewis, Claude Lepage, Tristan Glatard, Konrad Wagstyl, Jordan DeKraker, Paule-J Toussaint, Sofie L Valk, Louis Collins, et al. The bigbrainwarp toolbox for integration of bigbrain 3d histology with multimodal neuroimaging. Elife, 10:e70119, 2021.

[177] Russell A Poldrack, Aniket Kittur, Donald Kalar, Eric Miller, Christian Seppa, Yolanda Gil, D Stott Parker, Fred W Sabb, and Robert M Bilder. The cognitive atlas: toward a knowledge foundation for cognitive neuroscience. Frontiers in neuroinformatics, 5:17, 2011.

[178] Aaron F. Alexander-Bloch, Haochang Shou, Siyuan Liu, Theodore D. Satterthwaite, David C. Glahn, Russell T. Shinohara, Simon N. Vandekar, and Armin Raznahan. On testing for spatial correspondence between maps of human brain structure and function. NeuroImage, 178:540–551, 9 2018. ISSN 10959572. doi: 10.1016/j.neuroimage.2018.05.070. PMID: 29860082 publisher: Academic Press Inc.

[179] Justine Y. Hansen, Ross D. Markello, Jacob W. Vogel, Jakob Seidlitz, Danilo Bzdok, and Bratislav Misic. Mapping gene transcription and neurocognition across human neocortex. Nature Human Behaviour, pages 1–11, 3 2021. ISSN 2397-3374. doi:10.1038/s41562-021-01082-z. publisher: Nature Publishing Group.

[180] Ross D Markello and Bratislav Misic. Comparing spatial null models for brain maps. NeuroImage, 236:118052, 2021.

[181] Peter T Fox and Jack L Lancaster. Mapping context and content: the brainmap model. Nature Reviews Neuroscience, 3(4):319–321, 2002.

[182] David C Van Essen, Stephen M Smith, Deanna M Barch, Timothy E.J. Behrens, Essa Yacoub, and Kamil Ugurbil. The wu-minn human connectome project: An overview. NeuroImage, 80:62–79, 2013. ISSN 10538119. doi: 10.1016/j.neuroimage.2013.05.041. PMID: 23684880.

[183] Matthew F Glasser, Stamatios N Sotiropoulos, Anthony Wilson, Timothy S Coalson, Bruce Fischl, Jesper L Andersson, Junqian Xu, Saad Jbabdi, Matthew Webster, Jonathan R Polimeni, David C Van Essen, and Mark Jenkinson. The minimal preprocessing pipelines for the human connectome project. Neuroimage, 80:105–124, 2013. doi:10.1016/j.neuroimage.2013.04.127.

[184] Fang-Cheng Yeh, Sandip Panesar, David Fernandes, Antonio Meola, Masanori Yoshino, Juan C Fernandez-Miranda, Jean M Vettel, and Timothy Verstynen. Population-averaged atlas of the macroscale human structural connectome and its network topology. NeuroImage, 178:57–68, 2018. ISSN 10959572. doi: 10.1016/j.neuroimage.2018.05.027. PMID: 29758339.

[185] Fang-Cheng Yeh, Van Jay Wedeen, and Wen-Yih Isaac Tseng. Estimation of fiber orientation and spin density distribution by diffusion deconvolution. NeuroImage, 55(3):1054–1062, 4 2011. ISSN 1053-8119. doi: 10.1016/J.NEUROIMAGE.2010.11.087. publisher: Academic Press.

[186] Fang-Cheng Yeh, T D Verstynen, Y Wang, J C Fernández-Miranda, and W-Yi Tseng. Deterministic diffusion fiber tracking improved by quantitative anisotropy. PLoS ONE, 8(11):80713, 2013. doi: 10.1371/journal.pone.0080713.

[187] Andrea I Luppi and Emmanuel A Stamatakis. Combining network topology and information theory to construct representative brain networks. Network Neuroscience, 5(1):96–124, 2021. ISSN 2472-1751. doi: 10.1162/netn_a_00170.

[188] Richard F Betzel, Alessandra Griffa, Patric Hagmann, and Bratislav Mišić. Distance-dependent consensus thresholds for generating group-representative structural brain networks. Network neuroscience, 3(2):475–496, 2019.

[189] Bratislav Mišić, Richard F Betzel, Azadeh Nematzadeh, Joaquin Goni, Alessandra Griffa, Patric Hagmann, Alessandro Flammini, Yong-Yeol Ahn, and Olaf Sporns. Cooperative and competitive spreading dynamics on the human connectome. Neuron, 86(6):1518–1529, 2015.

[190] Olivier David, Anne-Sophie Job, Luca De Palma, Dominique Hoffmann, Lorella Minotti, and Philippe Kahane. Probabilistic functional tractography of the human cortex. Neuroimage, 80:307–317, 2013.

[191] Lena Trebaul, Pierre Deman, Viateur Tuyisenge, Maciej Jedynak, Etienne Hugues, David Rudrauf, Manik Bhattacharjee, François Tadel, Blandine Chanteloup-Foret, Carole Saubat, et al. Probabilistic functional tractography of the human cortex revisited. Neuroimage, 181: 414–429, 2018.

