## Supplementary Figures for "The blueprint of human functional architecture shifts from cognition to anatomy during perturbations of consciousness"

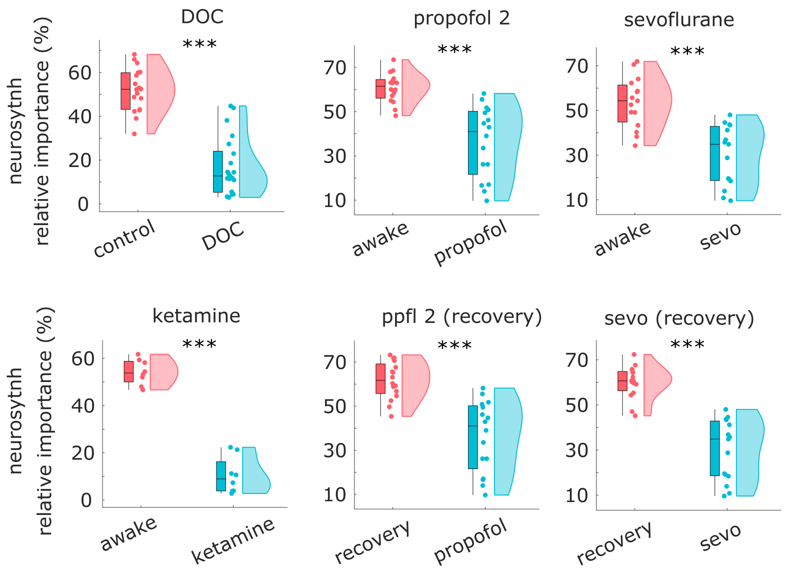


Figure S1. **Statistical comparisons of contributions (relative importance) from NeuroSynth meta-analysis |** Box-plots: center line, median; box limits, upper and lower quartiles; whiskers, $1.5\times$ interquartile range. *, $p < 0.05$; **, $p < 0.01$; ***, $p < 0.001$.

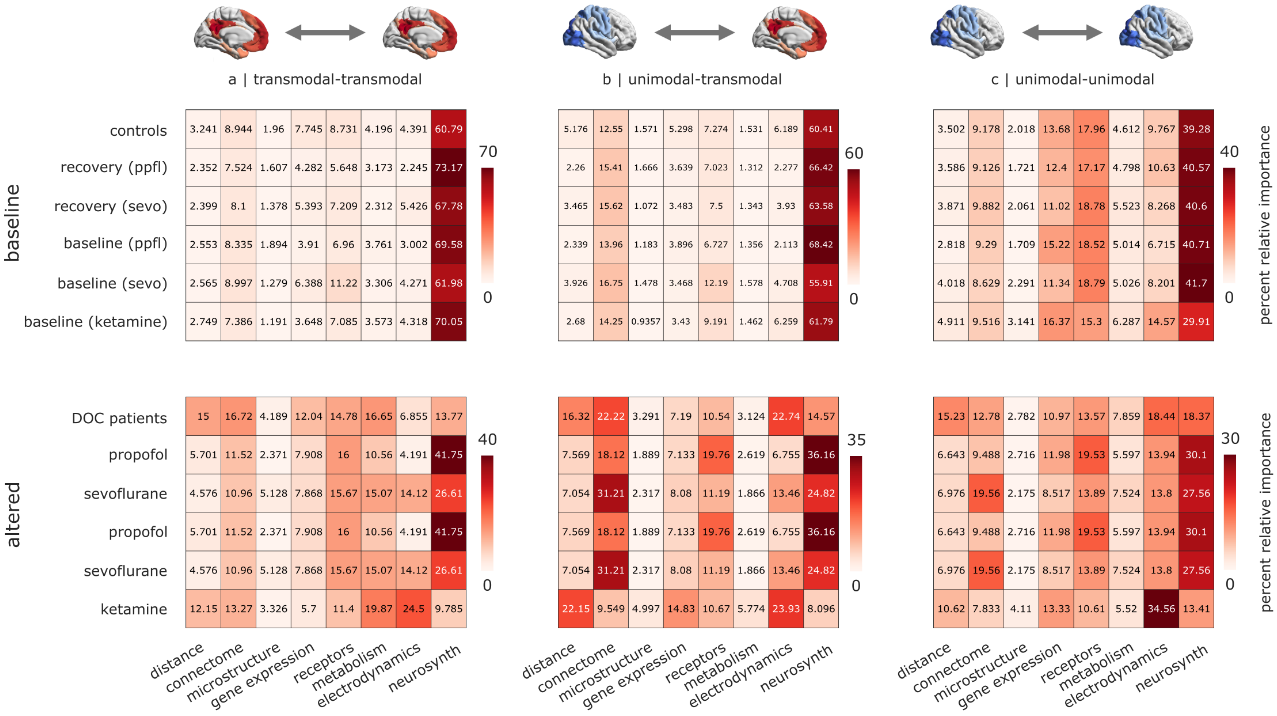


Figure S2. **Cognitively-relevant co-activation is the dominant predictor of haemodynamic functional connectivity across unimodal and transmodal cortices |** The relative importance of meta-analytic co-activation is largest for connections between transmodal regions (a), intermediate for connections between unimodal and transmodal (b), and lowest for connections between unimodal regions (c).

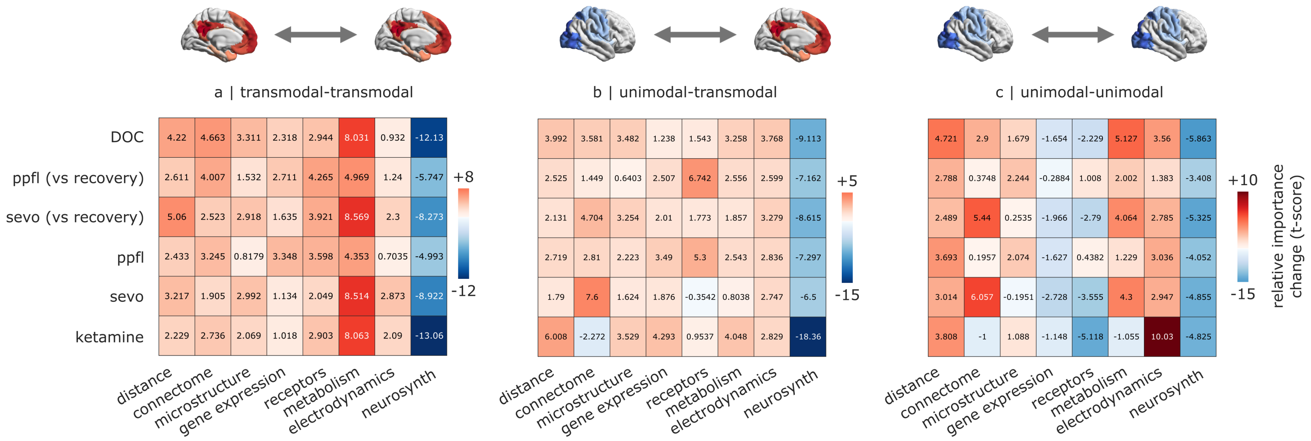


Figure S3. **Change in relative importance of each predictor network as a function of dataset and cortical type | (a)** Transmodal-transmodal. **(b)** Unimodal-transmodal. **(c)** Unimodal-unimodal (c).

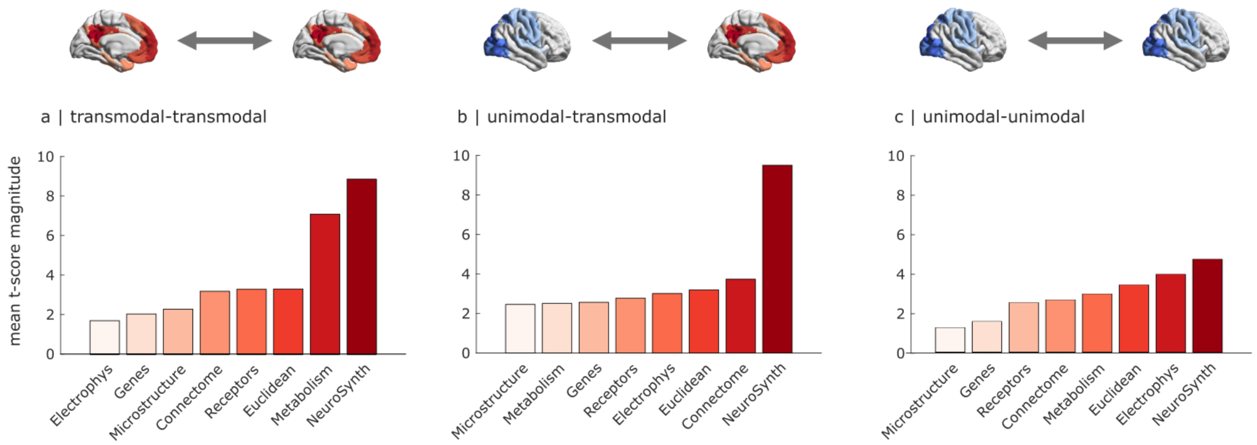


Figure S4. **Meta-analytic co-activation is the most affected predictor by anaesthesia and DOC, across cortical types | (a)** Transmodal-transmodal. **(b)** Unimodal-transmodal. **(c)** Unimodal-unimodal.

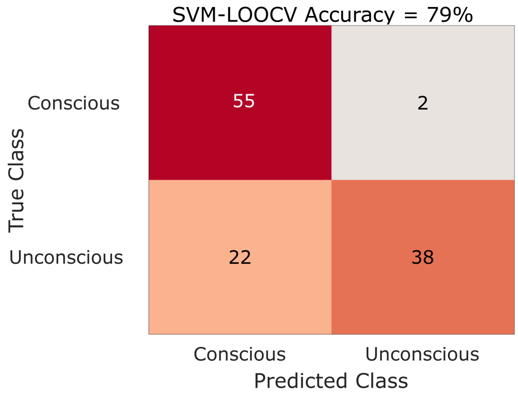


Figure S5. **Machine learning classification of consciousness from univariate network contributions** | Confusion matrix for the classification of pharmacological and pathological perturbations of consciousness from univariate correlations with brain networks, achieving 79% accuracy after training a support vector machine using leave-one-out cross-validation.

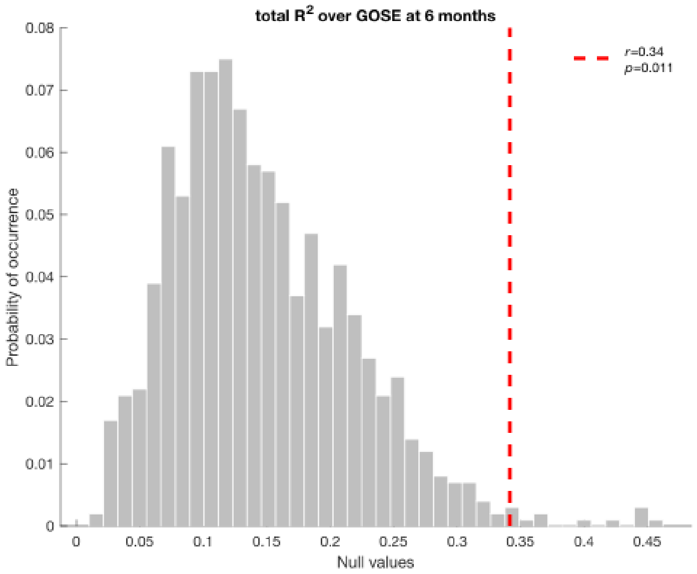


Figure S6. **Machine learning prognosis of 6-month GOSE scores |** Total $R^2$ from multiple regression, using relative importance of different networks to predict 6-month Glasgow Outcome Scale–Extended (GOSE) scores for 48/51 DOC patients in the Paris dataset for whom such scores are available. Significance is assessed against a null distribution of total $R^2$ obtained from 1000 models with reshuffled labels.

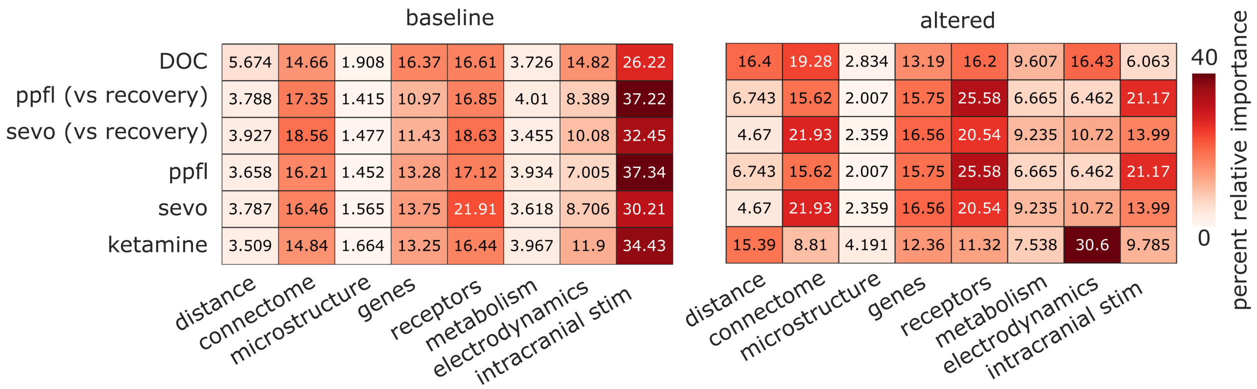


Figure S7. **Contribution of multimodal networks to haemodynamic functional connectivity under wakefulness and under perturbed consciousness is replicated with the network of similarity of subjective experiences elicited by intracranial electrical stimulation | (a)** Wakefulness, including pre-anaesthesia as well as post-anaesthetic recovery of responsiveness, and healthy controls for the DOC dataset. **(b)** Altered consciousness, including anaesthesia and patients with disorders of consciousness. For comparison, the re-test condition of the test-retest dataset is also included. Note that the first scan of the test-retest dataset is used as the control group for the DOC patients.

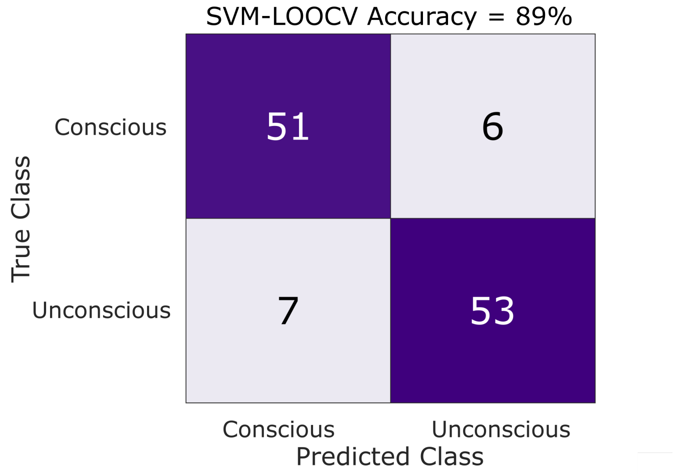


Figure S8. **Machine learning classification of consciousness from multimodal network contributions is replicated with the network of similarity of subjective experiences elicited by intracranial electrical stimulation |** Confusion matrix for the classification of pharmacological and pathological perturbations of consciousness using the relative importance of multimodal networks, achieving 89% accuracy (p < 0.001) after training a support vector machine using leave-one-out cross-validation.

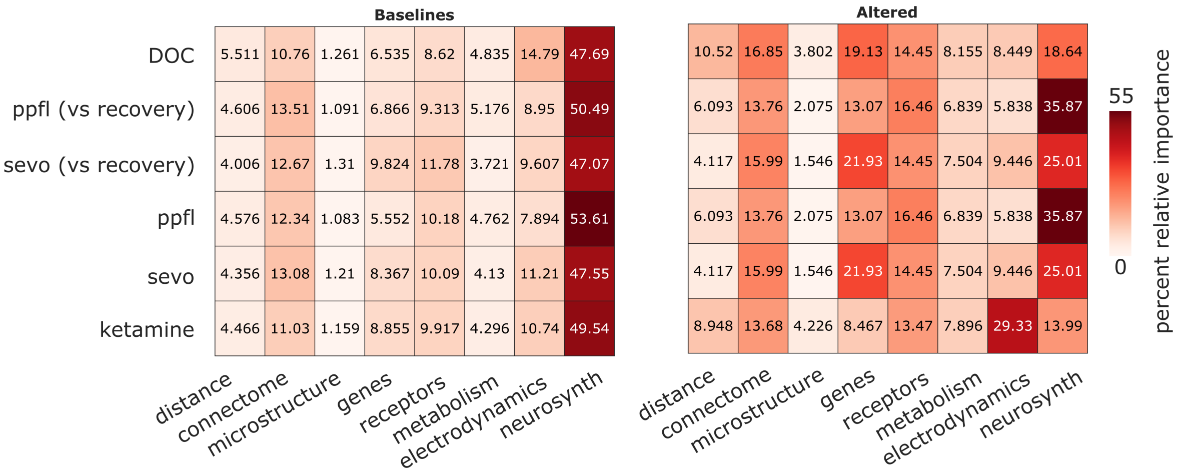


Figure S9. **Contribution of multimodal networks to haemodynamic functional connectivity under wakefulness and under perturbed consciousness is replicated with the anatomical Desikan-Killiany atlas | (a)** Wakefulness, including pre-anaesthesia as well as post-anaesthetic recovery of responsiveness, and healthy controls for the DOC dataset. **(b)** Altered consciousness, including anaesthesia and patients with disorders of consciousness. For comparison, the re-test condition of the test-retest dataset is also included. Note that the first scan of the test-retest dataset is used as the control group for the DOC patients.

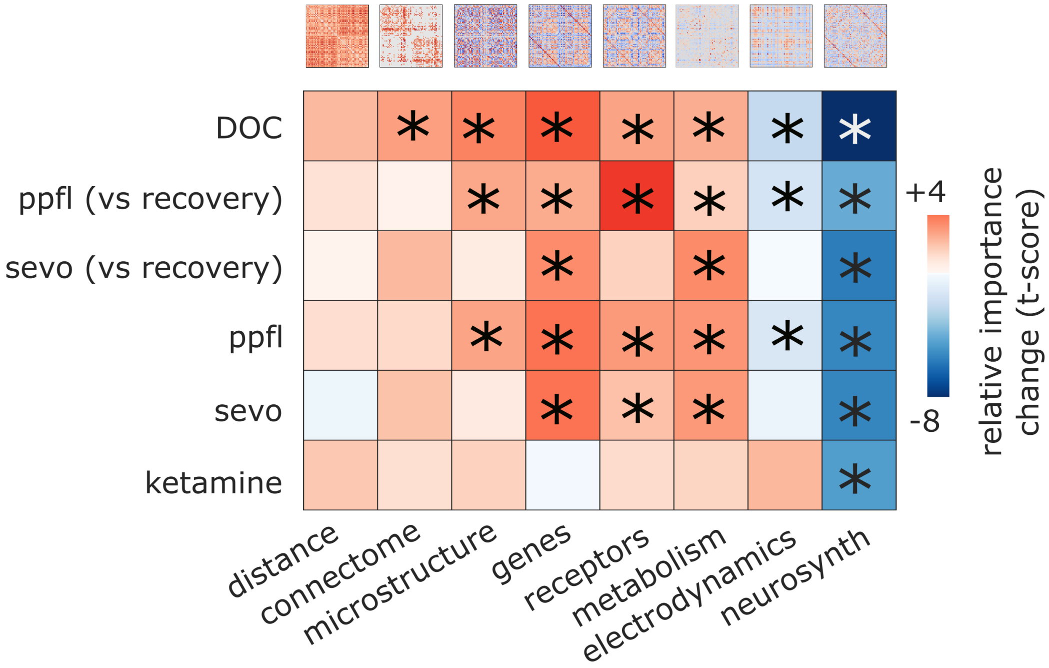


Figure S10. **Reduced contribution of meta-analytic co-activation to haemodynamic functional connectivity is replicated with the anatomical Desikan-Killiany atlas |** *, $p < 0.05$ after controlling for head motion. Red color-scale indicates altered > baseline. Blue color-scale indicates baseline > altered.

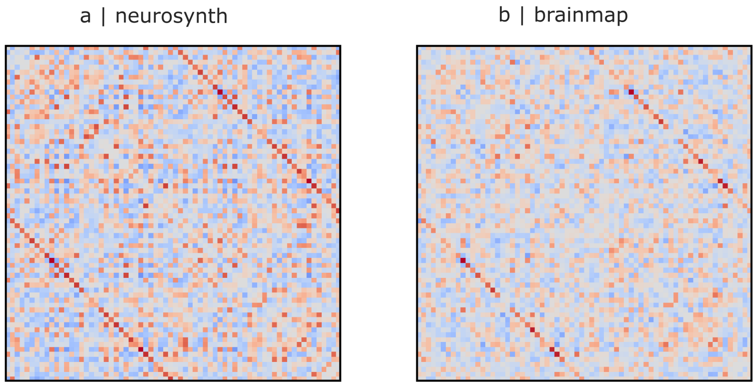


Figure S11. **Alternative quantifications of meta-analytic co-activation |** (**a**) NeuroSynth automated meta-analysis. (**b**) BrainMap expert-curated meta-analysis.

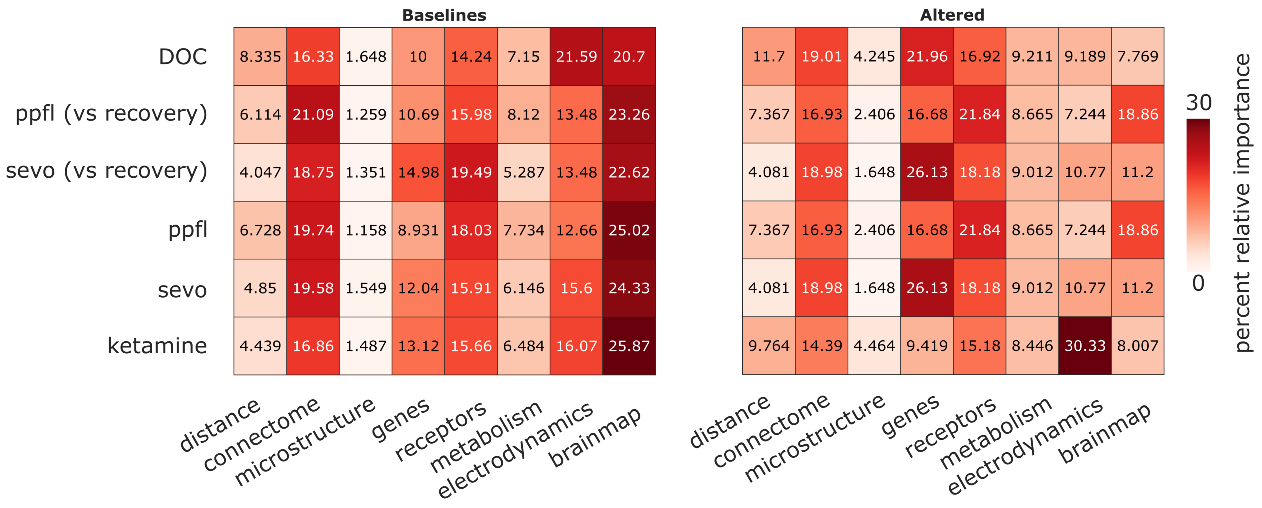


Figure S12. **Contribution of multimodal networks to haemodynamic functional connectivity under wakefulness and under perturbed consciousness is replicated with BrainMap meta-analytic database | (a)** Wakefulness, including pre-anaesthesia as well as post-anaesthetic recovery of responsiveness, and healthy controls for the DOC dataset. **(b)** Altered consciousness, including anaesthesia and patients with disorders of consciousness. For comparison, the re-test condition of the test-retest dataset is also included. Note that the first scan of the test-retest dataset is used as the control group for the DOC patients.

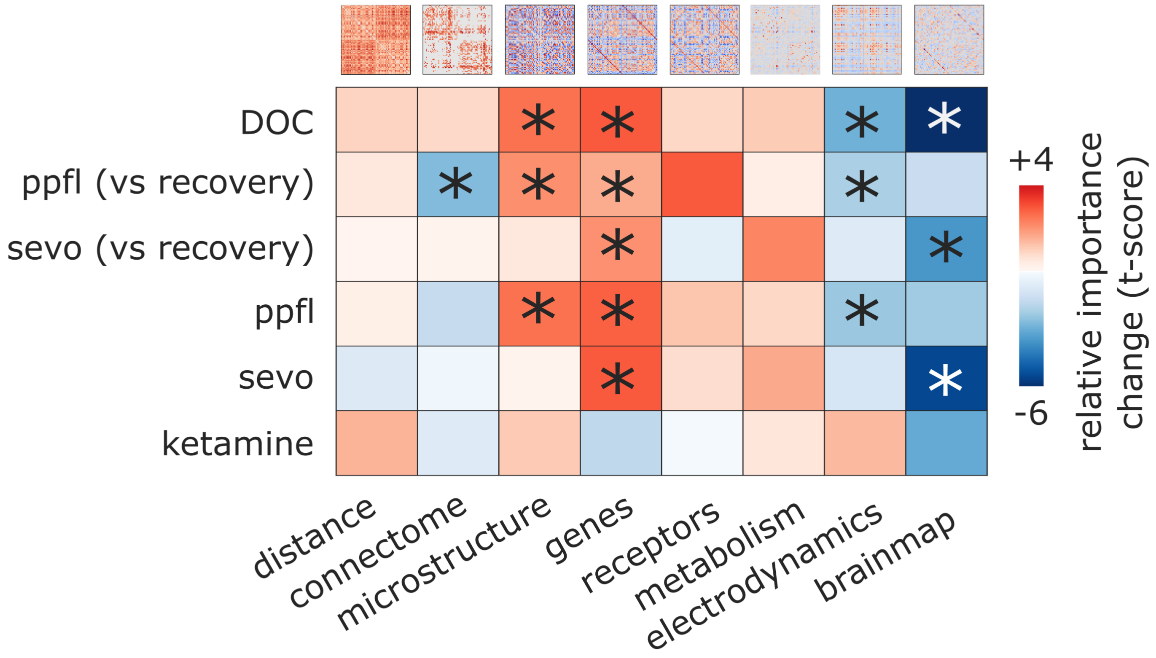


Figure S13. **Reduced contribution of meta-analytic co-activation to haemodynamic functional connectivity is replicated with BrainMap meta-analytic database |** *, $p < 0.05$ after controlling for head motion. Red color-scale indicates altered > baseline. Blue color-scale indicates baseline > altered.

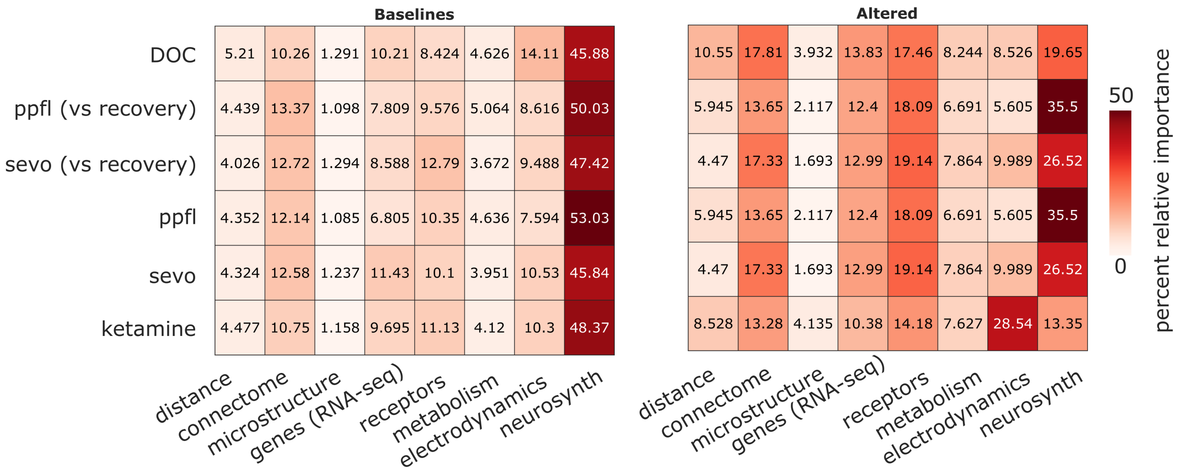


Figure S14. **Contribution of multimodal networks to haemodynamic functional connectivity under wakefulness and under perturbed consciousness is replicated with RNAseq gene expression | (a)** Wakefulness, including pre-anaesthesia as well as post-anaesthetic recovery of responsiveness, and healthy controls for the DOC dataset. **(b)** Altered consciousness, including anaesthesia and patients with disorders of consciousness. For comparison, the re-test condition of the test-retest dataset is also included. Note that the first scan of the test-retest dataset is used as the control group for the DOC patients.

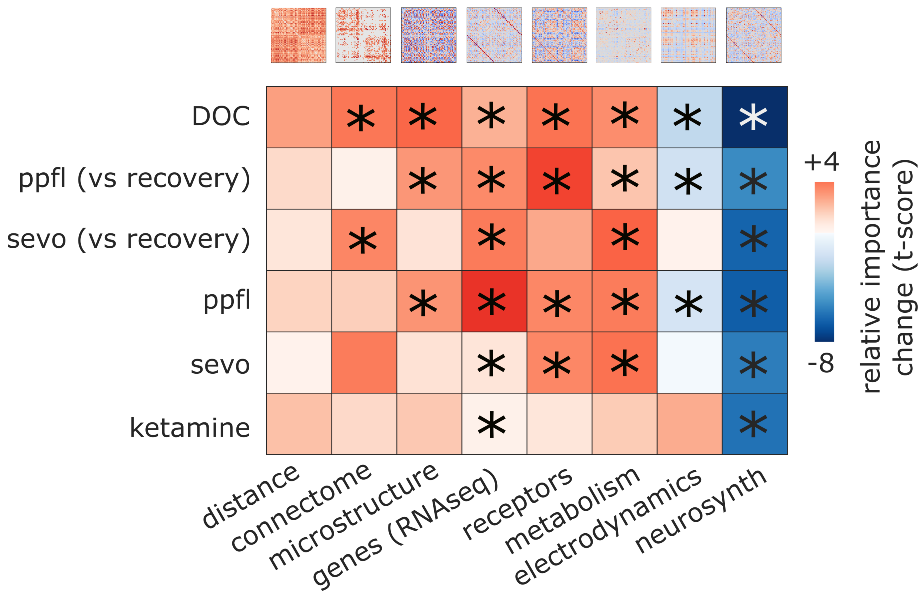


Figure S15. **Reduced contribution of meta-analytic co-activation to haemodynamic functional connectivity is replicated with RNAseq gene expression |** *, $p < 0.05$ after controlling for head motion. Red color-scale indicates altered > baseline. Blue color-scale indicates baseline > altered.

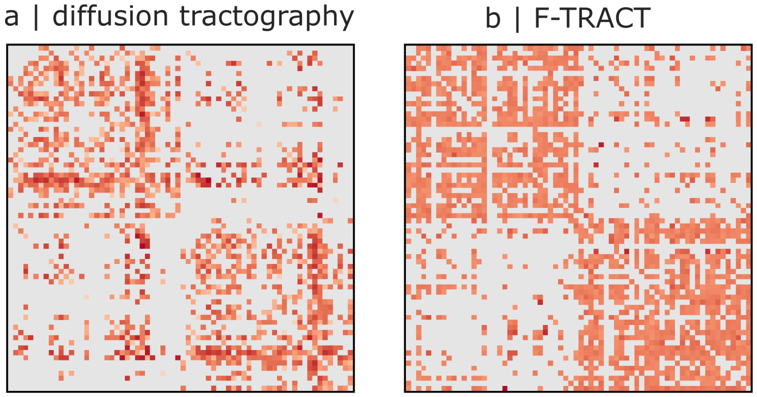


Figure S16. **Alternative quantifications of anatomical connectivity | (a)** Structural connectome from diffusion tractography. **(b)** F-TRACT connectome from cortico-cortical evoked potentials.

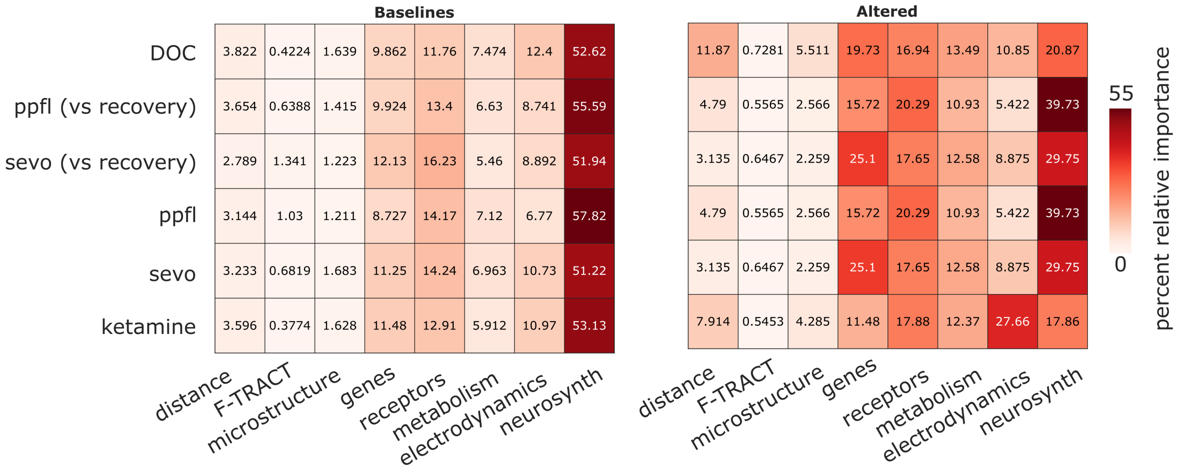


Figure S17. **Contribution of multimodal networks to haemodynamic functional connectivity under wakefulness and under perturbed consciousness is replicated with connectome estimated from cortico-cortical evoked potentials | (a)** Wakefulness, including pre-anaesthesia as well as post-anaesthetic recovery of responsiveness, and healthy controls for the DOC dataset. **(b)** Altered consciousness, including anaesthesia and patients with disorders of consciousness. For comparison, the re-test condition of the test-retest dataset is also included. Note that the first scan of the test-retest dataset is used as the control group for the DOC patients.

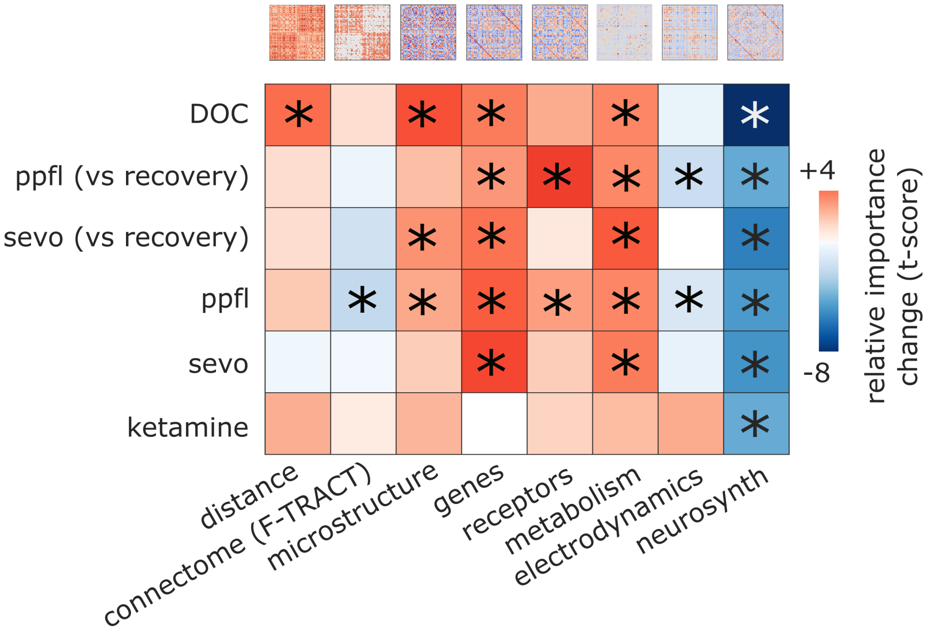


Figure S18. **Reduced contribution of meta-anaytic co-activation to haemodynamic functional connectivity is replicated with connectome estimated from cortico-cortical evoked potentials** | *, $p < 0.05$ after controlling for head motion. Red color-scale indicates altered > baseline. Blue color-scale indicates baseline > altered.

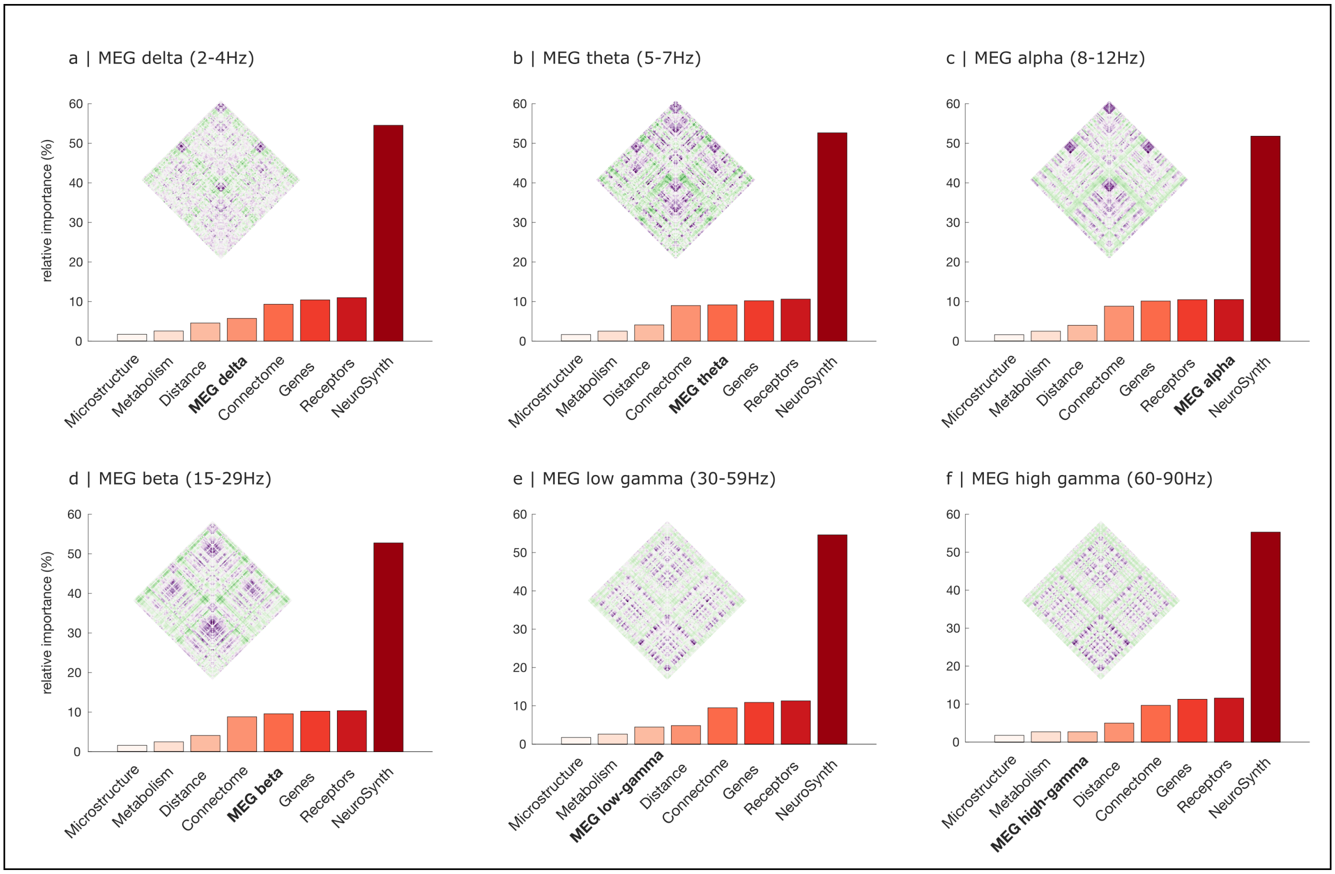


Figure S19. **Cognitively-relevant co-activation is the dominant predictor of haemodynamic functional connectivity regardless of MEG frequency band choice |** (**a**) MEG delta band (2-4Hz). (**b**) MEG theta band (5-7Hz). (**c**) MEG alpha band (8-12Hz). (**d**) MEG beta band (15-29Hz). (**e**) MEG low-gamma band (30-59Hz). (**f**) MEG high-gamma band (60-90Hz).

**Baselines**

| | Distance | Connectome | Microstructure | Genes | Receptors | Electrophys | Metabolism | NeuroSynth | Perfusion |
|---|---|---|---|---|---|---|---|---|---|
| DOC | 4.146 | 6.546 | 1.579 | 7.123 | 7.176 | 8.64 | 1.924 | 38.3 | 24.56 |
| ppfl (vs recovery) | 3.524 | 7.092 | 1.56 | 4.923 | 6.632 | 4.443 | 1.828 | 46.07 | 23.93 |
| sevo (vs recovery) | 3.655 | 7.894 | 1.418 | 5.052 | 7.358 | 5.376 | 1.635 | 46.42 | 21.19 |
| ppfl | 3.281 | 6.896 | 1.586 | 5.775 | 6.869 | 3.6 | 1.872 | 44.99 | 25.13 |
| sevo | 4.221 | 7.284 | 1.342 | 5.992 | 9.147 | 4.965 | 1.934 | 37.74 | 27.37 |
| ketamine | 4.367 | 7.194 | 1.23 | 6.142 | 7.61 | 7.443 | 2.092 | 39.77 | 24.15 |

**Altered**

| | Distance | Connectome | Microstructure | Genes | Receptors | Electrophys | Metabolism | NeuroSynth | Perfusion |
|---|---|---|---|---|---|---|---|---|---|
| DOC | 12.93 | 13.92 | 2.259 | 8.581 | 10.33 | 14.29 | 7.017 | 12.4 | 18.27 |
| ppfl (vs recovery) | 5.321 | 9.318 | 1.743 | 8.671 | 14.76 | 4.792 | 4.303 | 26.53 | 24.57 |
| sevo (vs recovery) | 4.704 | 13.66 | 1.742 | 9.226 | 11.04 | 8.449 | 5.818 | 21.49 | 23.87 |
| ppfl | 5.321 | 9.318 | 1.743 | 8.671 | 14.76 | 4.792 | 4.303 | 26.53 | 24.57 |
| sevo | 4.704 | 13.66 | 1.742 | 9.226 | 11.04 | 8.449 | 5.818 | 21.49 | 23.87 |
| ketamine | 13.02 | 7.131 | 3.507 | 9.672 | 8.716 | 28.74 | 6.339 | 7.789 | 15.09 |

percent relative importance: 0 – 45

Figure S20. **Contribution of multimodal networks to functional connectivity under wakefulness and under perturbed consciousness is replicated when adding blood co-perfusion as predictor | (a)** Wakefulness, including pre-anaesthesia as well as post-anaesthetic recovery of responsiveness, and healthy controls for the DOC dataset. **(b)** Altered consciousness, including anaesthesia and patients with disorders of consciousness. For comparison, the re-test condition of the test-retest dataset is also included. Note that the first scan of the test-retest dataset is used as the control group for the DOC patients.

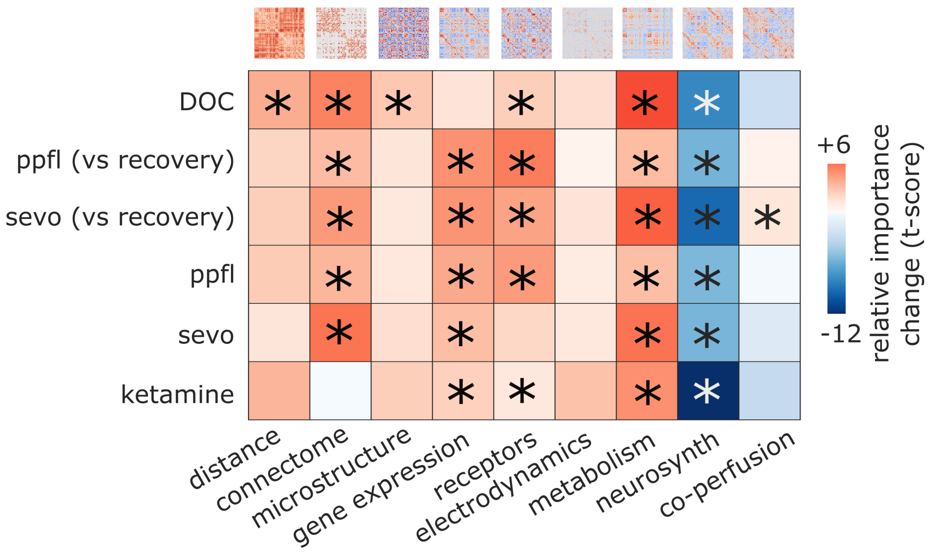


Figure S21. **Reduced contribution of meta-analytic co-activation to functional connectivity is replicated when adding co-perfusion as predictor |** *, $p < 0.05$ after controlling for head motion. Red color-scale indicates altered > baseline. Blue color-scale indicates baseline > altered.
